# Elucidating tissue and sub-cellular specificity of the entire SUMO network reveals how stress responses are fine-tuned in a eukaryote

**DOI:** 10.1101/2025.04.08.647708

**Authors:** Jason Banda, Shraboni Ghosh, Dipan Roy, Kishor D. Ingole, Lisa Clark, Eshan Sharma, Sumesh Kakkunath, Kawinnat Sue ob, Rahul Bhosale, Leah Band, Srayan Ghosh, Darren Wells, Jonathan Atkinson, Malcolm J. Bennett, Kathryn Lilley, Andrew Jones, Miguel De Lucas, Anthony Bishopp, Ari Sadanandom

## Abstract

SUMOylation is essential in plant and animal cells, but it remains unknown how SUMO components act in concert to modify specific targets in response to environmental stresses. In this study we characterise every SUMO component in the Arabidopsis root to create the first complete SUMO Cell Atlas in eukaryotes. This unique resource reveals wide spatial variation, where SUMO proteins and proteases have sub-functionalized in both their expression and subcellular localisation. During stress SUMO conjugation is mainly driven by tissue specific regulation of the SUMO E2 ligase. Stress-specific modulation of the SUMO pathway reveals unique combinations of Proteases being targeted for regulation in distinct root tissues by salt, osmotic and biotic signals. Our SUMO Cell Atlas resources reveal how this PTM influences cellular and tissue scale adaptations during root development and stress responses. We provide the first comprehensive study elucidating how multiple stress inputs can regulate an entire PTM system.

---

Post-translational modifications (PTMs) are crucial in orchestrating biological complexity and modulate nearly every biological process. This is particularly evident in multicellular organisms where development requires precise coordination of cell fates.^1^ There are >100 different PTMs, including ubiquitination and phosphorylation.^1,2,3^ Unravelling the mechanisms regulating these PTMs can be challenging due to the very large number of components involved. However, compared with the many hundreds of proteins involved in ubiquitination, experimentally characterising the 30-40 components required for the SUMO (Small Ubiquitin-like Modifier) pathway is far more tractable. SUMOylation is a key mechanism for rapid control of protein degradation, activity, localisation and binding partners,^4,5^ regulating an array of complex biological processes, from stress perception to chromatin modification and transcription. ^6–8^ Given its importance in cell function and its tractable genetics, SUMO allows us to delve deeper into how multicellular organisms use PTMs to integrate stress signals by modifying key substrates to coordinate downstream effects on adaptive responses.

SUMO is ∼11 kDa, slightly larger than ubiquitin and is found in all eukaryotes. The SUMO modifier protein has a tertiary structure consisting of the β-grasp orientation structure,^9^ which is also common to other protein conjugation systems like Ubiquitin and Nedd8. SUMO genes are translated as premature peptides with C-terminal extensions to form a diglycine motif. The C-terminal end is processed to expose a di-glycine termini, which is then conjugated to lysine residues in target substrates through a cascading collection of enzymes sequentially named E1, E2, E3, and E4. SUMO is covalently linked to its substrate as single or multiple monomers, or as polymers composed of different SUMO peptides catalysed by the E4 enzyme.^10^ The SUMO E2 enzyme can also facilitate SUMO conjugation directly to its substrate protein or to the SUMO E3 Ligase for conjugation onto substrates. The Ubiquitin-like Protease (ULP) class of SUMO proteases can process premature SUMO and remove SUMO from substrates, while DeSUMOylating Isopeptidases (DeSI) exclusively remove SUMO from targets. Together, these proteases regulate the pool of free SUMO within cells. **(Figure S1)**.

Families of all major components of the SUMO cycle, except USPL1 proteases, are present in multiple eukaryotic kingdoms, suggesting similar functions. However, there are some noticeable differences between the animal and plant SUMO systems. Animal SUMO systems have a significantly increased number of E3 ligases, eight genes, whereas in plants only two have been confirmed to date.^11,12^ Conversely, plants have a more extensive number of SUMO peptides and proteases than animals.^9^ The expansion of SUMO ligase genes in animals and protease genes in plants may provide insights into how a single PTM system diverged to suit each kingdom’s different developmental and adaptive needs.

Despite SUMO modification enzymes being well studied in both plant and animal kingdoms, it is unknown how SUMO components act in concert to modify their targets in response to changing environments. Furthermore, despite their critical importance in numerous cellular processes, our understanding of the rules governing SUMO target specificity remains rudimentary. We use the root of the plant Arabidopsis thaliana as a model to address this. We report the spatial expression and protein localization of every SUMO system component to create the first complete SUMO Cell Atlas (SCA) and reveal how SUMO influences cellular and tissue scale adaptations during plant development and stress responses. In doing so, we provide a comprehensive network elucidating how multiple inputs can regulate a single PTM.

To understand the role of SUMO in regulating cellular states, fates and molecular processes, it is crucial to map the spatio-temporal distribution of the SUMO-machinery components **(Figure S1)** within a model organ. Given its simple and robust cellular organisation, the *Arabidopsis* primary root is an ideal organ for developing a SUMO Cell Atlas (SCA). The *Arabidopsis* primary root is derived from the root apical meristem which is made up of proximal and distal stem cells separated by an organising centre (the quiescent centre, QC). The proximal portion of the root comprises a series of distinct cell types (epidermis, cortex, endodermis and pericycle) arranged in concentric layers around a central vascular cylinder (comprising phloem, xylem and procambial cells).^13^ As new cells divide in the meristematic stem cell niche, mature cells are displaced away from this zone. These displaced cells then undergo a period of expansion before differentiating into their final cell fates. The timings of each phase of root cell development (division, elongation, differentiation) are tightly controlled by genetic and environmental factors, resulting in three distinct zones within the root tip; meristem, elongation and differentiation zones.^13^

We initial analysed published single-cell RNA-Seq data and observed differential expression for all SUMO components in different cell types **(Figure S2)**. For example, *SUMO3* is expressed in epidermal cells yet not in the vasculature and quiescent centre cells. *SUMO3* expression correlated with SUMO proteases SPF1, OTS2 and DeSi3B, with high expression only in the meristem zone (MZ), indicating a potential functional relationship between these components. To validate the observed co-expression-based functional association between the SUMO components across different cell types, we needed to simultaneously analyse gene expression, protein abundance and subcellular localisation of each component.

To generate an expression atlas for every Arabidopsis SUMO component, we synthesised transcriptional and translational fluorescent fusions for all 32 SUMO cycle genes using an innovative reporter design. This dual transcriptional and translational reporter contained a promoter region followed by the genomic sequence fused to an HA-tagged mVenus-3xHA-2A-mTurquoise-N7 dual reporter **(Figure 1A**). The 2A peptide induces ribosomal skipping during translation creating a SUMO-mVenus-HA translational reporter and a co-transcribed nuclear localised mTurquoise reporter, both driven by the same promoter.^14–16^ The translational reporter will denote the spatial domain, at both (sub)cellular and tissue scales, in which the protein is located. The second peptide, mTurquoise-N7 defines the spatial domain where transcription/translation of the gene occurs. Any differences in the spatial location of the two fluorescent signals will reveal whether SUMO proteins exhibit cell-to-cell movement or whose abundance are regulated via degradation.

**Figure 1.**
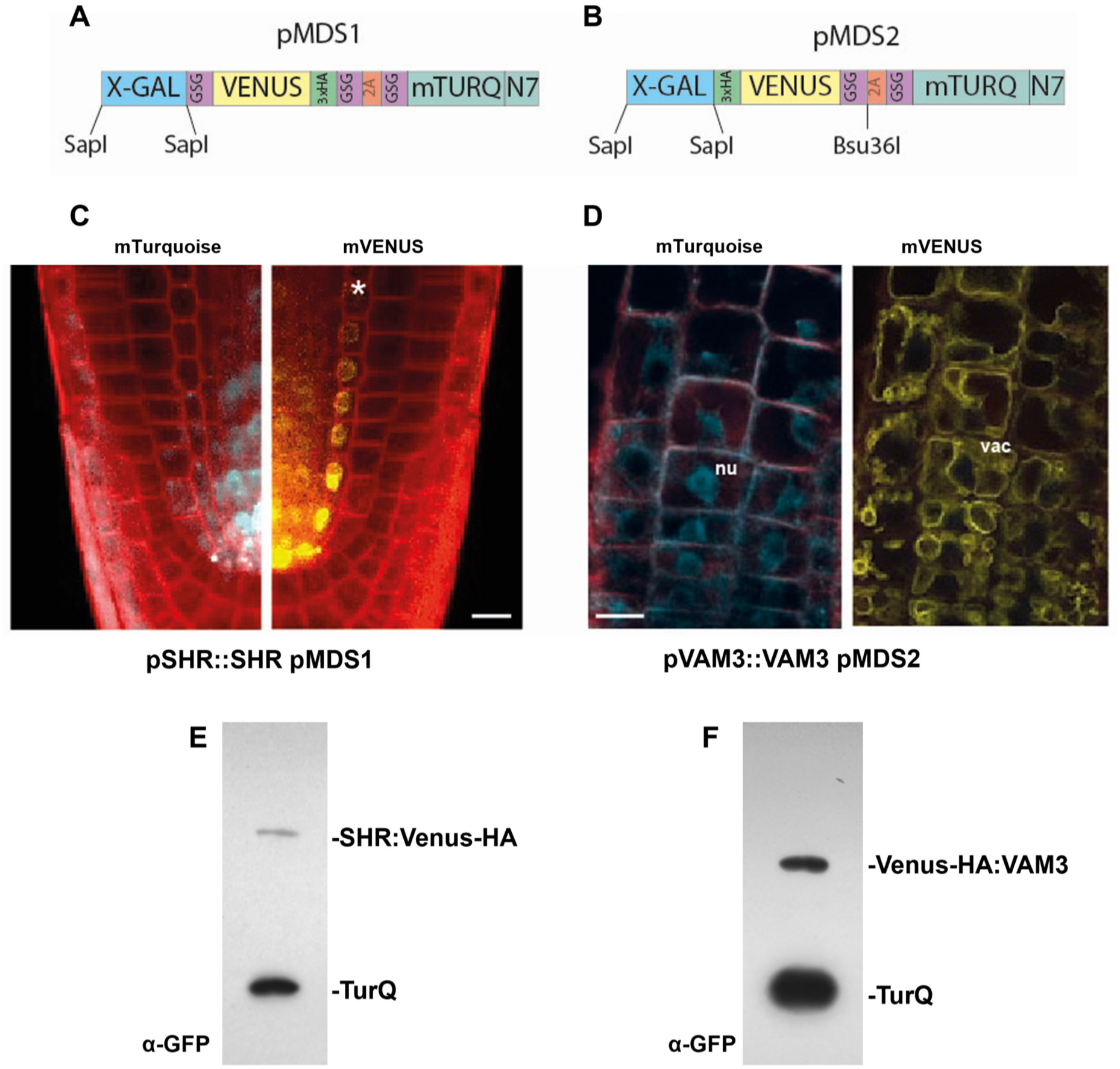
Description and validation of pMDS plasmid system for dual analysis of transcription and translation in plants. (A) Organization of pMDS1 vector showing reporters for transcription (mTurQ), translation (C-terimal mVenus) and a 2A self-cleaving peptide. (B) Organization of pMDS2 vector showing reporters for transcription (mTurQ), translation (N-terminal mVenus) and a 2A self-cleaving peptide. (C) Confocal image of pMDS1_*SHR*pro:*SHR:mVenus:mTurQ* showing gene expression (mTurQ) in the stele region and protein (mVenus) translocating to endodermis in root meristem. (D) Confocal image of pMDS2_*VAM3*pro:mVenus:*VAM3:mTurQ* showing sub-cellular expression (mTurQ) in nucleus and protein (mVenus) moving to vacuole in root epidermis. Red channel shows mCherry expression. Nu (Nucleus), Vac (Vacuole), * Endodermis of root meristem, Scale bar=10µM

To validate the functionality of our polycistronic reporter approach, we analysed the transcriptional and translational behaviour of two well-characterised genes: SHORTROOT (SHR) and VACUOLAR MORPHOLOGY 3 (VAM3). SHR is a transcription factor that regulates root radial patterning via its non-cell autonomous behaviour.^17^ The SHR gene is initially transcribed in the vascular cells of the root apical meristem (RAM), but its protein moves into adjacent endodermal and cortex initials.^17^ The *SHR* promoter and genomic coding region was cloned into the vector (termed pMDS1; Figure 1A) to create a C-terminal reporter fusion. Transgenic plants exhibited a mTurquoise-N7 signal in the stele (denoting the SHR transcription domain) while the mVenus signal (denoting SHR protein distribution) was expanded to include endodermal and cortex initials **(Figure 1A and C**), demonstrating SHR protein movement and validating our dual reporter approach. Furthermore, efficient 2A skipping of the construct was verified using VAM3, a Q-SNARE receptor that locates to vacuolar and pre-vacuolar compartments cloned into a vector variant termed pMDS2. In transgenic roots, VAM3 gene expression (visualised with mTurquoise-N7) was present in the nucleus of epidermal cells, while the VAM3 protein was only visible in the vacuole of these cells **(Figure 1B and D**). Additionally, western blot analysis of protein extracts from VAM3 pMDS2 transgenic lines demonstrated efficient ribosome skipping conferred by the 2A signal **(Figure 1E and F**), as no hybrid peptide at higher molecular weight was detected. These outcomes indicate that the pMDS vector system works efficiently and allows us to simultaneously visualise expression and protein localisation. This will enable us to ascertain whether rapid changes in gene expression or protein levels of SUMO components is due to induction and/or degradation.

To create a SUMO cell atlas for a plant organ we characterised each component’s expression and subcellular localisation in the key Arabidopsis root cell types (i.e. epidermis, cortex, endodermis and vasculature) and zones (i.e. meristem and elongation; **Figure 2A and B**). We quantified each SUMO component’s expression at both translation (using mTurquoise) and protein (using mVenus) levels in each root cell type and zone, then collated these values into five categories of expression **Figure 2C**) (low-high). This approach allows us to define levels of each SUMO component per tissue and zone. To validate that the genomic fragments of the SUMO components cloned into the pMDS vectors faithfully replicated endogenous genes expression, we performed Q-RTPCR to compare transgene and endogenous SUMO gene expression. Two independent transgenic lines for each SUMO component exhibiting similar gene expression patterns to the corresponding endogenous gene were taken forward for further analysis (Figure S3). Furthermore, efficient 2A skipping of each of the SUMO component constructs was verified using immunoblot analysis **(Figure S4-7)**. Finally, to validate that SUMO components tagged with mVENUS remain functional, we successfully complemented mutants in SUMO components with known phenotypes (Figure S8 and 9).

**Figure 2:**
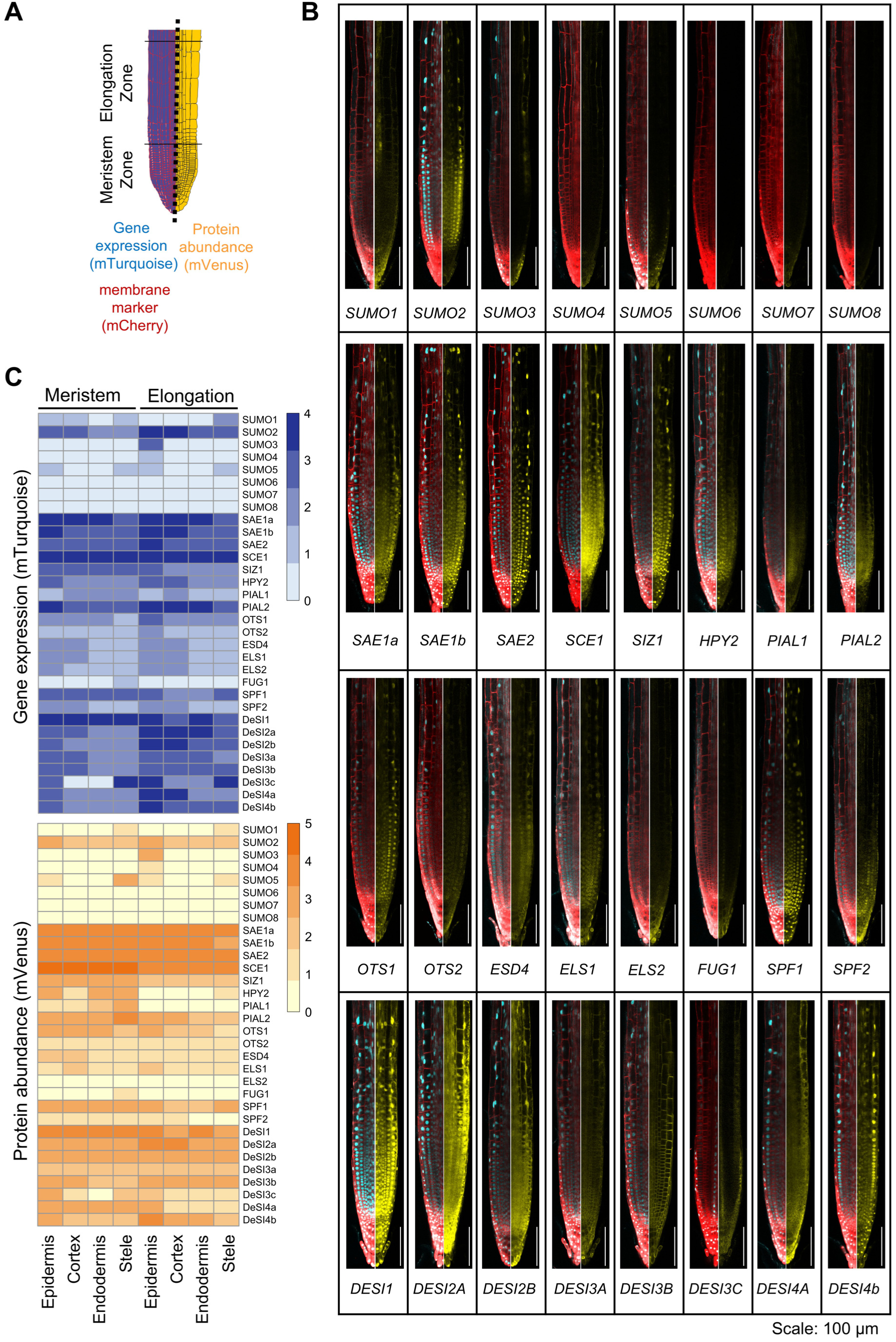
SUMO components exhibit distinct domains of expression and protein accumulation in Arabidopsis root tips. (A) Schematic representation of the root tip zones and tissues measured in the Arabidopsis root. (B) Confocal images displaying gene expression (cyan) and protein localisation (yellow) of all 32 SUMO components in the root tip of *Arabidopsis thaliana*. The cell outline is shown in red by a genetic membrane marker. Scale bar, 100 μm. (C) Heatmap representing the semi-quantitative level of gene expression and protein abundance of all SUMO components in the varying root zones and tissues. 0 indicated no fluorescence observed, while level 4 or 5 defines high level of fluorescence.

The SUMO Cell atlas revealed several surprising insights. Earlier work assumed SUMO1 and SUMO2 were ubiquitously expressed within root tissues,^18^ but using our reporter lines, we observed discrete expression patterns for these two modifiers. SUMO2 was ubiquitously expressed in all root cell types and zones measured. In striking contrast, SUMO1 exhibited a much more restricted pattern, with high levels of expression in a subset of vasculature cells in the elongation zone, which we confirmed as phloem companion cells after analysing root cross sections **(Figure S10)**. Single-cell Seq datasets supported both patterns. These differences in transcriptional regulation of SUMO1 and 2 provide a basis for sub-functionalization, where these modifiers mediate responses to different stimuli. Discrete expression patterns were also observed for other SUMO peptides in our reporter lines. SUMO3 was detected in epidermal cells within the elongation zone. SUMO4 was expressed at very low levels in the late elongation zone and SUMO5 demonstrated very low expression in vasculature and epidermis. The remaining SUMO6, SUMO7 and SUMO8 were not detected in the root tip yet could be involved in older root tissue or in shoot organs. This wide variation in the spatial expression of SUMO genes could enable cell-type specific conjugation of targets.

E1 – E4 enzymes form the core SUMO conjugation machinery. The genes encoding these key components exhibited the highest levels of expression and protein abundance in the SUMO cycle, especially SAE1a, SAE1b, SAE2 and SCE1. The E3, HPY2 and E4, PIAL1 had the lowest expression and the E3, SIZ1 was intermediate in expression in all root tissues and zones analysed. HPY2 is expressed in both root zones, but the protein is only visible in the meristematic zone. This observation is consistent with the reported role for HPY2 as a regulator of cell proliferation in the apical meristem where it functions as a repressor of the endocycle **(Figure 2B**).^19,20^ The SUMO chain-forming E4 ligase PIAL1 is expressed at low levels in multiple root cell-types, but its protein is only found in early meristematic tissue. Interestingly, a number of the E1-E4 enzymes are localised in both the nucleus and cytosol. Particularly SCE1, PIAL1, PIAL2 and SAE1a exhibit strong localisation to the cytosol, whereas SAE1b and HPY2 show minor protein abundance in the cytosol. This suggests that SUMOylation of targets could occur outside the nucleus, explaining the SUMOylation of plasma membrane targets like FLS2.^21^

ULP proteases perform a dual function; SUMO maturation through cleaving the pre-protein to release the peptide and SUMO removal from its target substrates. In general, this group of proteases displayed low expression levels and protein abundance. Most ULPs are expressed in all root cell types, however FUG1 is exclusively observed in pericycle cells (a layer of cells surrounding the vascular tissues that gives rise to lateral roots), where FUG1 might play a role in epigenetic gene silencing.^22^ SUMO maturation is considered to occur only in the nucleus, however the OTS2 protease is localised exclusively to the nuclear envelope. Previous work demonstrated binding of human SENPs to nuclear pore complexes in the nuclear envelope, where the proteases are involved in localisation and transport kinetics.^23^ Hence, OTS2 may perform similar roles in plants. ELS1 and ESD4 are also located in both nucleus and cytoplasm. They may fulfil a dual role between peptidase activity in the nucleus and isopeptidase activity in the cytoplasm.

In contrast to the ULP class, DeSI proteases contain an isopeptidase domain that cleaves SUMO off substrates but does not function during SUMO maturation. This distinct family of DeSI proteases is more highly expressed and abundant than its ULP peptidase counterpart. DESI’s exhibit expression in all root tissues and zones with the exception of DESI3c which is limited to the lateral root cap and vasculature, with weak expression in the epidermis. Interestingly, DeSI showed a wide variety of subcellular locations **(Figure S11)**. DESI1 and DES4b were strongly localised to the nucleus with weak cytosolic abundance. DESI2a and DESI2b displayed similar protein levels in both the nucleus and cytosol. DESI3a and DESI3b are localised to the plasma membrane, while Desi3c contains both membrane and cytosolic localisation. The outlier is DESI4a, which is specifically localised to the vacuole membrane in the elongation zone as vesicles start fusing to form one main vacuole **(Figure 2B**). Little is known about the molecular targets of these DESIs, but our SUMO Cell Atlas reveals a strong subcellular variation, indicating that individual DESIs may target a discrete set of targets in different cellular components. Our SUMO system wide approach raises the possibility of a much more complex regulatory landscape for SUMO modification than previously known.

The 32 dual reporter lines within our Cell Atlas resource were next employed to establish how the SUMO system modulates the SUMOylated proteome upon different environmental cues. We studied three model abiotic and biotic stresses; salt, osmotic and bacterial elicitor, flagellin. For salt and flagellin treatment, seedlings were exposed to 150mM NaCl or 1uM flagellin. For osmotic stress, transgenic seedlings were transferred to agar plates containing either 300mM of mannitol or control plates.

We firstly verified that SUMO systems respond after 3-hour of exposure to model stimuli. Immunoblot analysis confirmed SUMO conjugate accumulation 3-hour post-treatment (Figure S12). Next, we quantified changes in SUMO components gene expression 3h after treatment using bulk RNA Seq on whole roots. The salt and mannitol treatments have similar numbers of DEGs **(Figure S13)**. Flagellin treatment, however, only triggered relatively mild transcriptomic changes. In the salt and mannitol treatment, GO analysis revealed up-regulated genes were enriched in osmotic homeostasis signalling pathways (response to salt, response to water deprivation, etc.) **(Figure S13B)**. Flagellin treatment up-regulated known defence response genes, but down-regulated glucosinolate and carbohydrate metabolism.

The same treatments at 3-hour time point were used to visualise the response of the 32 SUMO component reporters. Seedlings were scanned on the confocal microscope to visualise both mTurquoise and mVenus patterns. The abundance of every SUMO components was quantified in multiple cell-types, including epidermis, cortex, endodermis and stele; and within two root zones, meristem and elongation zone. The changes induced by each stress within these cell types were visualised as expression **(Figure 4-6, panel A)** and protein abundance **(Figure 4-6, panel B)** heatmaps to reveal common and contrasting spatial and quantitative regulatory effects on each SUMO component.

Salt stress profoundly impacts root growth and development.^24,25^ It is well documented that salt stress induces SUMO conjugation to substrate proteins in roots **(Figure S12)**.^26,27^ However, it remains unclear how the SUMO system responds to salt stress. SUMO cell atlas resources revealed that after 3h of salt stress, SUMO1 and 2 showed were downregulated (both gene expression and protein abundance) while SUMO3 and 5 showed slight upregulation **(Figure 3**). The SUMO1 protein was strongly inhibited in the stele of the meristem, whilst SUMO2 was reduced across all cell types in the elongation zone **(Figure 3A and B**). This spatial specificity in downregulation suggests specific developmental roles for these two highly similar SUMO isoforms in salt response. The observation that levels of SUMO3 and SUMO5 are increased after salt stress suggests a novel role for these modifiers in salt stress responses.

**Figure 3:**
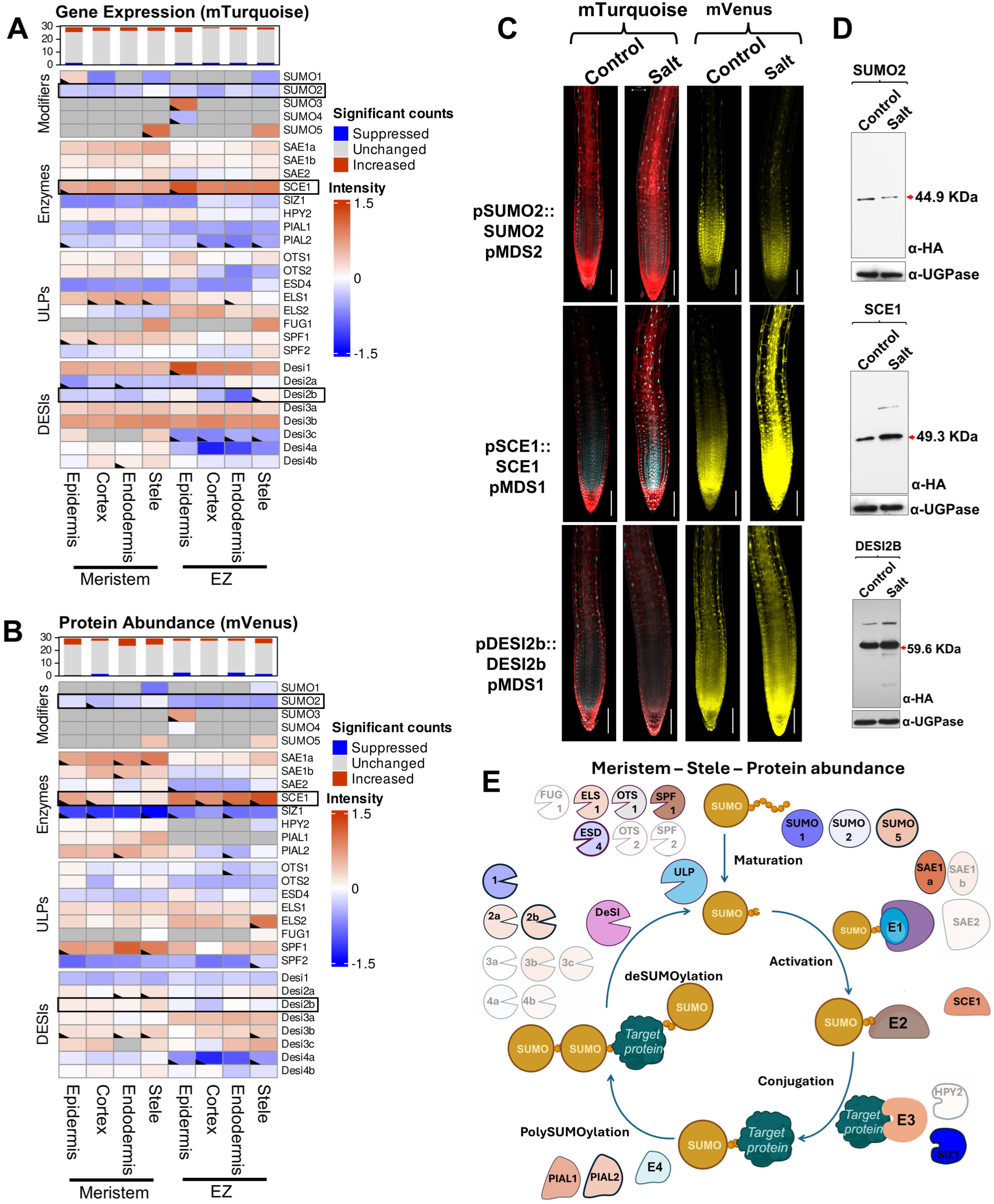
Salt stress modulates the SUMO system in a tissue-specific manner. (A) Heatmap representing the log 2-fold changes in gene expression within the SUMO components 3 hrs after 150 mM salt stress in several root tissues and zones. The top bar graph displays the number of SUMO genes per tissue type in a specific zone that change in a statistically significant manner (p=<0.05). Dark grey boxes indicate combinations that were not analysed as no expression or protein was observed in this tissue. Note that boxes with a marked corner indicate significant changes. Examples of the most extreme changes are outlined and shown in panels C and D. (B) Similar heatmap as in panel A representing the log 2-fold changes in protein abundance within the SUMO components 3 hrs after 150 mM salt stress in several root tissues and zones. (C) Confocal microscopy images showing changes in expression (mTurquoise - cyan) and protein abundance (mVenus – yellow) of 3 major changers during salt stress. Scale bar, 100 μm. (D) Western blot of 3 major SUMO changers during salt stress confirming trend as seen in image analysis. UGPase was used as a loading control. (E) Schematic representation of the SUMO cycle and its components. The cycle represents the changes in protein abundance in the meristematic stele. Note that components with modest changes (fold change <0.5) are shown as transparent while those with the largest changes (log change >0.5) are shown in solid colour based on the scale of the heatmap in panel B. Components with statistically significant changes are also outlined.

Amongst the SUMO conjugation enzymes, the E2 enzyme, SCE1 showed the greatest induction after salt stress **(Figure 3C and D**). We observed significantly increased levels of SCE1 protein in the elongation zone compared to the meristem, with the highest expression in the epidermis and protein abundance in the stele. The epidermis is where salt is first perceived in the roots, so SCE1 induction in this tissue suggests a direct role for salt perception by the SUMO system. The E3 ligase SIZ1 was strongly downregulated across all cell types and zones, while HPY2 levels remain stable. These results suggest that during salt stress, SUMOylation relies more on increased levels of E2, with minimal contribution from E3 ligases. Interestingly, PIAL1 and 2 showed decreased expression, but increased protein abundance. These proteins are likely stabilised during salt stress, leading to increased poly-SUMO formation.

Among the SUMO proteases, OTS1 and OTS2 showed a clear reduction in protein level consistent with earlier studies on this subset of proteases, indicating that they are degraded after salt stress.^26^ ELS1 and SPF1 showed increased gene expression and protein abundance. As SPF1 gene expression is not strongly induced, this increase in protein abundance is likely regulated through increased protein stability under these conditions. SPF1 was strongly induced specifically in the endodermis and stele tissues. This implies a possible new role for this ULP protease in regulating salt responses in inner root tissues. Amongst the DESI-type proteases, the highly expressed DESI1 transcripts were upregulated in both meristem and elongation zones, primarily in the epidermal layer, although its protein levels decreased under salt stress. Likewise, DESI4a protein also showed a strong reduction in cortex and endodermis. This suggests that ULP and DESI proteases play specialised roles in different cell types under salt stress to shape their SUMOylated proteomes **(Figure 3E**).

Osmotic stress treatment upregulated both SUMO peptide and SUMO ligase components (Figure 4 A and B), consistent with previous work demonstrating increased SUMOylation in response to abiotic stresses.^21,28–30^ Both salt and osmotic stresses induce expression of *SAE1a, SCE1, SPF1 and DeSI1*. The increase in SCE1 protein abundance stands out in both datasets. However, where protein levels of SUMO peptides are generally downregulated in salt stress, during osmotic stress their abundance is increased **(Figure 4C, D)**. This trend could be the basis for the increase in SUMO conjugation seen upon osmotic stress treatment **(Figure S12)**. Interestingly, in SUMO2 and SCE1 the greatest changes in expression occur in inner tissues (endodermis and stele) of the elongation zone, suggesting a local function responding to osmotic changes within these root tissues.

**Figure 4:**
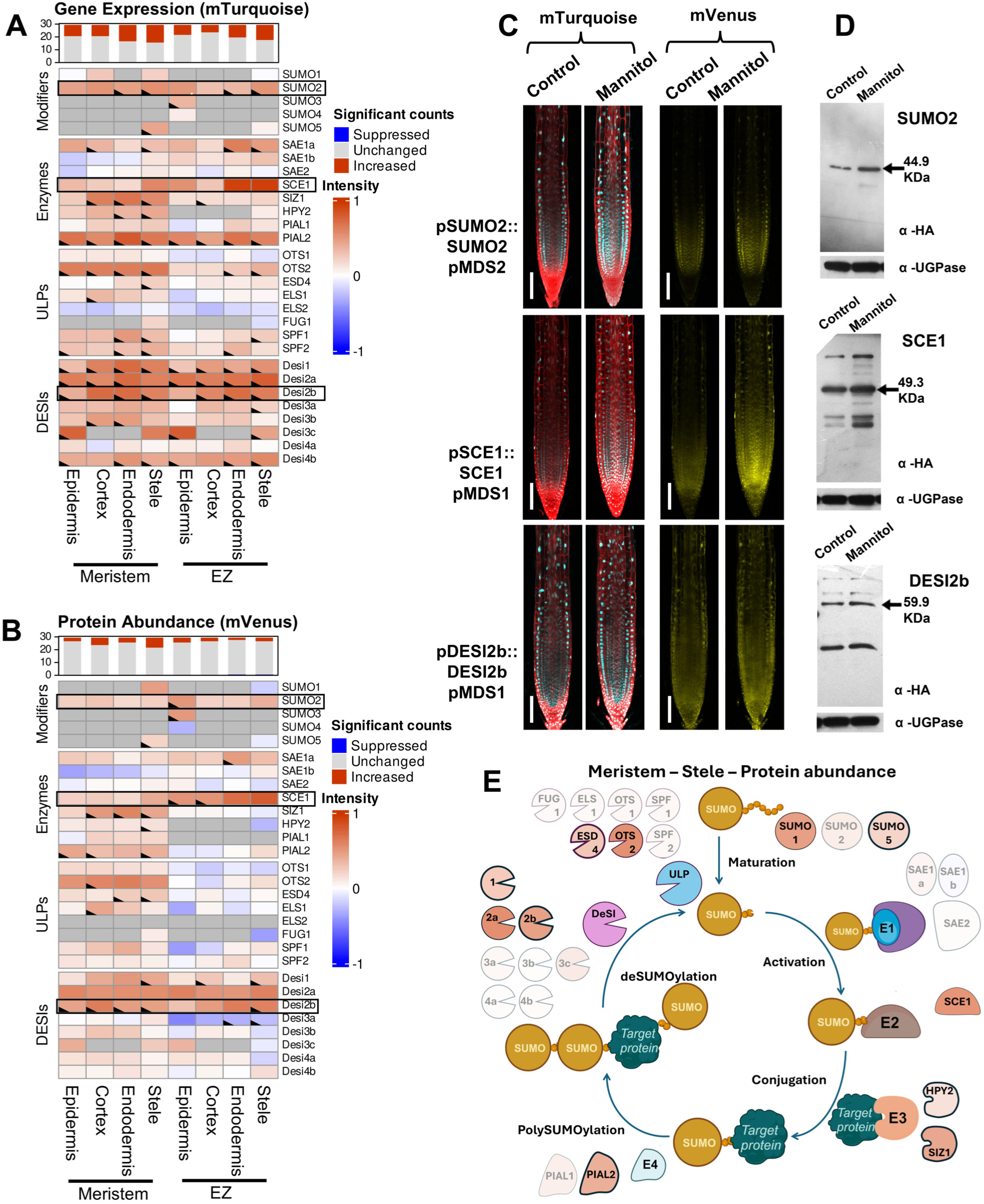
Osmotic stress regulates a distinct set of genes compared to salt stress. (A) Heatmap representing the log 2-fold changes in gene expression within the SUMO components 3 hrs after 300mM mannitol stress in several root tissues and zones. The top bar graph displays the number of SUMO genes per tissue type in a specific zone that change in a statistically significant manner (p=<0.05). Dark grey boxes indicate combinations that were not analysed as no expression or protein was observed in this tissue. Note that boxes with a marked corner indicate significant changes. Examples of the most extreme changes are outlined and shown in panels C and D. (B) Similar heatmap as in panel A representing the log 2-fold changes in protein abundance within the SUMO components 3 hrs after 300 mM mannitol stress in several root tissues and zones. (C) Confocal microscopy images showing changes in expression (mTurquoise - cyan) and protein abundance (mVenus – yellow) of 3 major changers during salt stress. Scale bar, 100 μm. (D) Western blot of 3 major SUMO changers during salt stress confirming trend as seen in image analysis. UGPase was used as a loading control. (E) Schematic representation of the SUMO cycle and its components. The cycle represents the changes in protein abundance in the meristematic stele. Note that components with modest changes (fold change <0.5) are shown as transparent while those with the largest changes (log change >0.5) are shown in solid colour based on the scale of the heatmap in panel B. Components with statistically significant changes are also outlined.

Strong increases in the expression and protein abundance of DeSI proteases 2a and 2b were also observed (Figure 4A and B). Whereas DeSI2a displays an increase in all tissues and zones, Desi2b increases mostly in endodermis and stele cell types, similar to SUMO2 and SCE1. This result reveals potential new roles for SUMO2 and SCE1 during osmotic stress by upregulating the SUMOylation of a wide group of targets, while Desi2a and Desi2b deSUMOylate specific targets to modulate stress response pathways.

Transcript and protein heatmaps reveal interesting zone-specific responses to osmotic stress for selected SUMO components (Figure 4A and B). For example, protein abundance of many ULP and DeSi proteases in the meristem is stable or increased after 3 h of osmotic stress. However, in the EZ a drop is visible for proteases such as OTS1, FUG1, SPF1 and DeSi3a (Figure 4B). In the case of OTS1 and FUG1, this can be directly ascribed to a decrease in gene expression (Figure 4A). Such contrasting behaviour suggests active breakdown of the protein in the EZ specifically during osmotic stress. In both SPF1 and OTS1 this trend is clearest in epidermal cells, which would perceive the stress first. The striking differences between root meristem and EZ highlights the importance of these two proteases in rapid osmotic stress responses.

Following osmotic stress treatment, most SUMO component expression changes are found in inner root tissues. Around 50% of the SUMO machinery genes showed significantly increased expression levels in the endodermis and stele of the root meristem versus ∼40% in the EZ. Significant changes in protein abundance were rarer, with the highest level of change in the meristem stele cells where 26% (7 out of 26) proteins increased versus two proteins increased (DeSi1 and DeSi2b) and one (DeSi3a) suppressed in the EZ. The meristem, therefore, appears to be the most reactive tissue when focussing on SUMO components. Focusing specifically on which proteins change within the stele, we observe changes in almost all E3 and E4 ligases (HPY2, SIZ1 and PIAL2) and changes in SUMO1, SUMO5 and SCE1. These changes are likely to cause an increased level of SUMO conjugation within stele cells. In contrast, levels of many ULP and DeSi proteases remain unchanged (Figure 4E). Only DeSI1, DeSi2b and ESD4 show significant increases. The elevation of SUMOylation versus deSUMOylation components likely explains the increased levels of SUMOylation observed after dehydration and osmotic stress. However, deSUMOylation of specific targets mediated by the increased abundance of DeSi components could play a vital role in activating specific stress pathways.

In parallel with our abiotic stress treatments, we observed that the biotic stress elicitor flg22 triggers dynamic changes of specific SUMO components primarily in the meristematic zone in a cell-type-specific manner **(Figure 5A and B**). Unlike abiotic stress treatments, changes in SUMO cycle machinery after flg22 application involved a smaller set of classes of components. The expression and protein abundance levels of all the SUMO modifiers remain unchanged, except for SUMO2, whose expression and protein abundance levels decreased in the meristematic zone **Figure 5C and D**). Flagellin treatment decreased both transcription and protein abundance levels of SAE1a, particularly at the meristematic zone. In contrast, the expression and protein abundance levels of SAE1b and SAE2 largely remained unchanged. However, we observed an increase in the levels of the E2 SUMO conjugating enzyme, SCE1, in the cell in the meristem more prominently than elongation zone **(Figure 5A and B**). Moreover, the protein abundance levels of the E3 ligase, HPY2, and the E4 ligase PIAL1 showed a slight but significant reduction at the meristematic zone.

**Figure 5:**
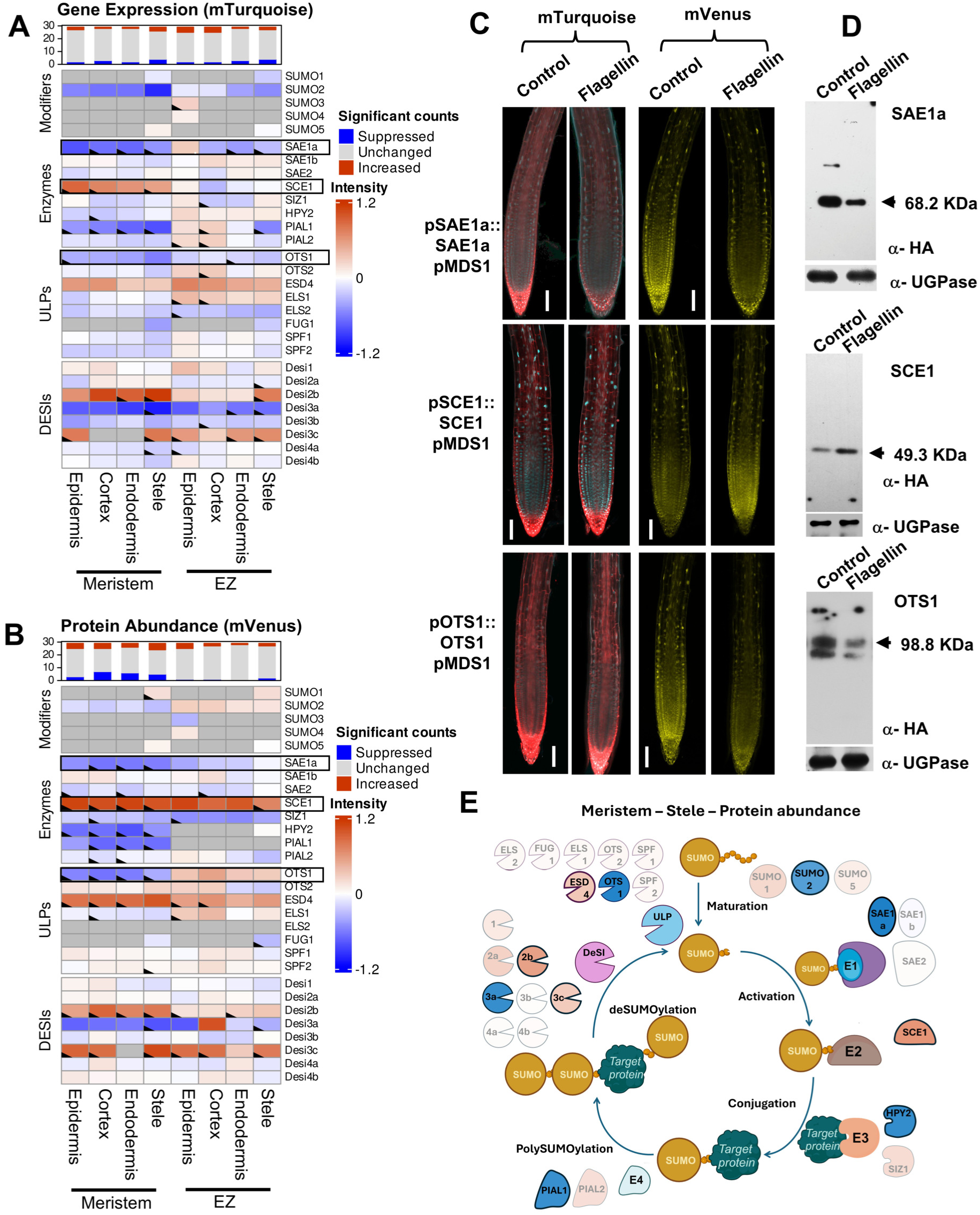
Biotic stress targets a distinct subset of SUMO system genes compared to abiotic stress. (A) Heatmap representing the log 2-fold changes in gene expression within the SUMO components 3 hrs after 1uM flagellin treatment in several root tissues and zones. The top bar graph displays the number of SUMO genes per tissue type in a specific zone that change in a statistically significant manner (p=<0.05). Dark grey boxes indicate combinations that were not analysed as no expression or protein was observed in this tissue. Note that boxes with a marked corner indicate significant changes. Examples of the most extreme changes are outlined and shown in panels C and D. (B) Similar heatmap as in panel A representing the log 2-fold changes in protein abundance within the SUMO components 3 hrs after 1uM flagellin treatment in several root tissues and zones. (C) Confocal microscopy images showing changes in expression (mTurquoise - cyan) and protein abundance (mVenus – yellow) of 3 major changers during salt stress. Scale bar, 100 μm. (D) Western blot of 3 major SUMO changers during salt stress confirming trend as seen in image analysis. UGPase was used as a loading control. (E) Schematic representation of the SUMO cycle and its components. The cycle represents the changes in protein abundance in the meristematic stele. Note that components with modest changes (fold change <0.5) are shown as transparent while those with the largest changes (log change >0.5) are shown in solid colour based on the scale of the heatmap in panel B. Components with statistically significant changes are also outlined.

Among the ULP proteases, there was a significant decrease in protein abundance of OTS1 particularly in the meristematic zone, while ESD4 was upregulated both at the transcriptional and translational level **(Figure 5E**). Amongst the components constituting the DESI SUMO proteases, there was strong upregulation at the transcriptional and translational level for DESI2B across all the examined tissue types in the meristematic zone. DESI3C also constituted a candidate DESI protease component that exhibited strong upregulation, particularly at the protein abundance level upon flagellin treatment across all examined tissue types in the meristematic and elongation zone except endodermis. Consistent with previous reports,^31^ flagellin treatment caused a strong decrease in protein levels of DESI3A. The response status of DESI3A upon flagellin treatment served as an internal positive reference control for our study confirming the efficiency of our experimental setup to elicit a biochemical and immune response in treated plants.

## Abiotic & Biotic stresses trigger differential tissue specific SCE1 interactome changes

The Arabidopsis genome contains a single SUMO E2 conjugation enzyme termed SCE1 **(Figure S1)**. Our cell atlas data revealed SCE1 is the only component of the SUMO system that is significantly increased under all three stresses **(Figure 3-5**). Hence, we hypothesise that this E2 is the main driver modulating stress-dependent increases in SUMO conjugation **(Figure S12)**.

We designed IP-MS experiments to identify SCE1 interacting partners under the three stresses to determine whether each treatment resulted in a distinct SUMOylated proteome. We performed these assays using root cultures expressing genomic SCE1 using the pMDS1 vector system **(Figure 1**). The interactome pull down assays were facilitated by the SCE1 protein being tagged at its C-terminus with both 3xHA and mVenus tags. After treatment of the roots with NaCl (150 mM), mannitol (300 mM), or flg22 (1 uM) for three hours, Immunoprecipitation was performed on extracted proteins using magnetic anti-GFP and anti-HA beads, then the interactomes were analysed using LC-MS/MS analysis (**Figure 6A and Figure S14)**.

**Figure 6.**
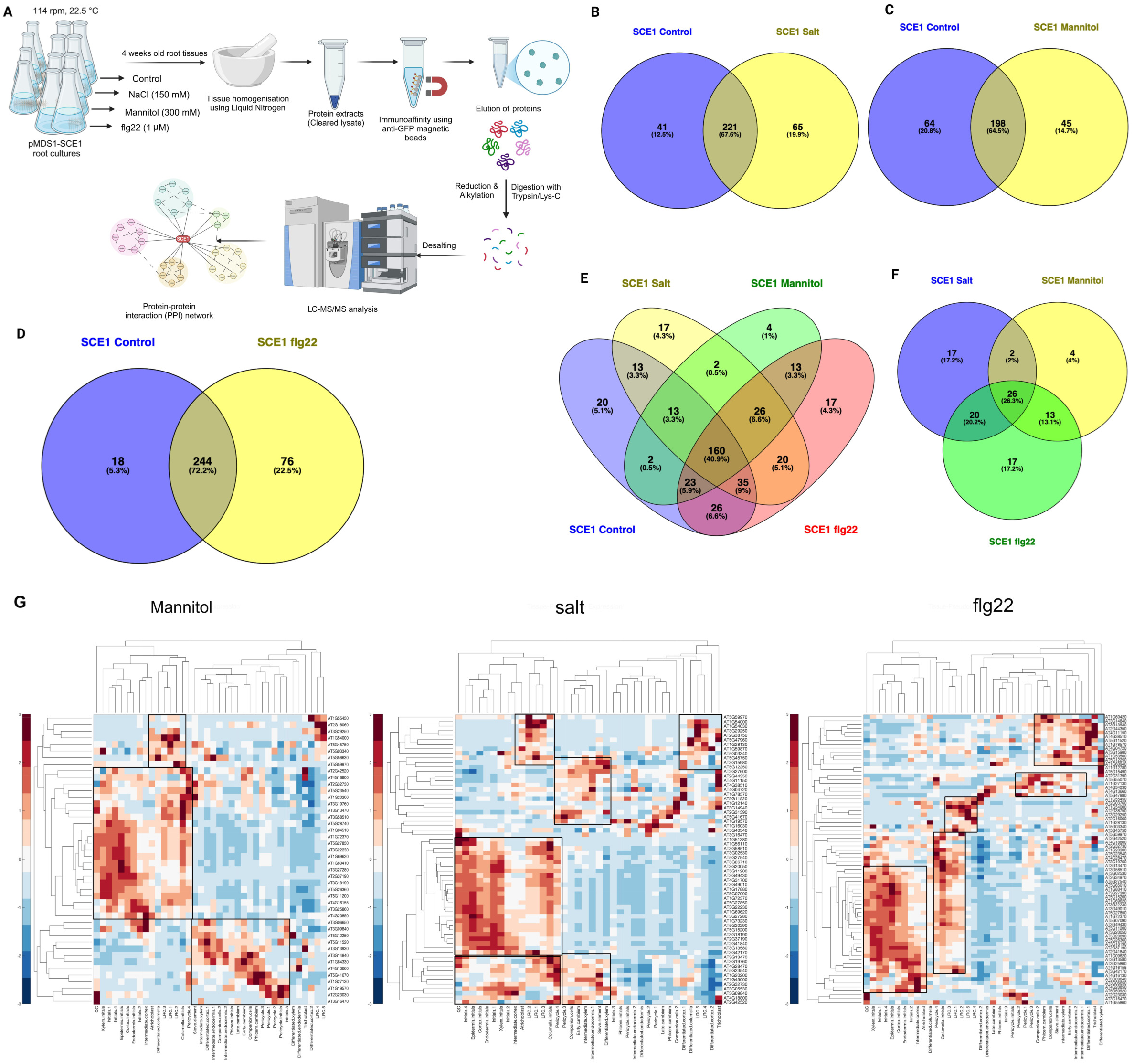
Stress-Specific SCE1-Interactome Characterized by IP-MS Analysis. A) Schematic representation of the experimental design and proteomics workflow employed for SCE1-IP-MS analysis under distinct stress conditions. B, C, D) Venn diagrams illustrating overlapping and stress-specific SCE1 interactors identified under salt, mannitol, and flg22 treatments, respectively. E) Venn diagram summarizing the overlap of SCE1 interactors across all tested stress conditions relative to the control. F) Venn diagram focusing on the overlap and uniqueness of stress-specific SCE1 candidates under salt, mannitol, and flg22 stresses. G) Expression analysis of SCE interactors identified from salt, mannitol, and Flg22 stress experiments across root tissues and their pseudo-time trajectories. The heatmap displays the proportion Z-score (-3 to 3) of standardized gene expression profiles across tissue types and zones. The red-blue colormap represents relative expression levels, with red indicating upregulation (+1 to +3 standard deviations above the mean) and blue indicating downregulation (-1 to -3 standard deviations below the mean), with the mean represented as 0. Black boxes highlight qualitative cluster groups, and dendrograms depict hierarchical clustering of genes (rows) and zones (columns) based on expression similarity.

LC-MS analysis identified 292, 286, 243, and 320 putative interactors of SCE1-GFP-HA under control, salt, mannitol, and flg22 treatments, respectively **(Figure 6B-F**). Deeper analysis of the stress-specific interactors indicated that 64 were unique to salt, 45 to mannitol, and 76 to flg22 treatments compared to unstressed controls **(Tables S5-7)**. Protein-protein interaction network analysis revealed that these stress-induced interacting targets are involved in various biological processes, highlighting their potential roles in response to distinct stress conditions (Figure S15). Of the stress-induced targets identified, 26 targets (26.3%) were common across different stress conditions, indicating that SCE1 interacts with these proteins regardless of the type of stress. In contrast, 17, 4, and 17 interactors were unique to salt, mannitol, and flg22 treatments, respectively **(Figure 6F**), suggesting these proteins might shape stress-specific SUMOylated proteomes. The interactors identified under these conditions are less likely to be the SUMOylated targets themselves as the pulldowns were performed under non-denaturing conditions where highly promiscuous protease activity dominates to remove SUMO from target substrates. However, the interactors of SCE1 may shape how the SUMO E2 is regulated to drive the formation of SUMOylated protein. Given that SCE1 protein abundance changes in different cell types during these stresses, we ascertained where the interactors of SCE1 are predominantly expressed within the root cell types by cross-analysis of SCE1 imaging data against publicly available scRNA-Seq data.^32^ This analysis reveals that under salt and mannitol stress SCE1 interacts more with proteins expressed in epidermis/atrichoblast cells, while under flg22 treatment, it interacts predominantly with proteins expressed in the endodermis/xylem tissue **(Figure 6G**). This data suggests that during abiotic stresses, the plant root SUMOylome is mainly derived from epidermis, while during flg22 treatment the SUMOylome is generated within endodermis and xylem cells suggesting that these inner tissues are the primary targets of the SUMO system for generating adaptive responses during biotic stress.

PTMs act at the core of biological processes by creating new proteoforms to transduce environmental cues into molecular responses. In this regard they amplify the genomic potential of organisms by creating multiple functionally different variants from a single gene product. SUMO is an essential PTM in eukaryotes.^9^ Despite its importance in cell function, it has not been possible to visualise the activity of an entire SUMO system in multicellular organisms in response to external cues until now. Here, we describe the construction of a Cell Atlas for the entire Arabidopsis SUMO machinery, the first of its kind in any eukaryote. To achieve this, we tagged all 32 components of the SUMO cycle involved in activating, conjugating and removing the SUMO peptide. In doing so, we provide a resource for all researchers working on plant SUMOylation which is located on the ePlant server (see resources availability statement). This web app visualises the expression, and localisation of all SUMO components in unstressed and stressed plants demonstrated as a heatmap. Seeds of all transgenic lines have been deposited in the Nottingham Arabidopsis Stock Centre. Additionally, through the pMDS1 and pMDS2 vectors we provide tools where others can simultaneously visualize gene expression and protein localisation of their gene of interest. These resources will provide molecular, spatio-temporal and regulatory information about the plant SUMO machinery that will allow researchers across different fields to explore the role of SUMOylation in a broad array of developmental programs and stresses and will be beneficial for a variety of abiotic and biotic stresses as well as those working in different tissues and different developmental stages.

Understanding how a PTM system is spatially resolved will provide fundamental information on how multicellular eukaryotes utilise proteoforms for cell signalling to produce an integrated response to changing environments. This study represents the first time a whole PTM has been spatially resolved in any eukaryote, and this has provided unique insights into how the SUMO system is organised and what drives its specificity. In Arabidopsis, the E1-E4 SUMO enzymes show broad expression across multiple cell types in the root tip. However, the E3, HPY2, and E4s PIAL1 and PIAL2 are enriched in the meristematic zone. Based on these proteins’ spatial specificity, a model is likely where SUMO is broadly conjugated to its targets across multiple cell types. Although most E1-E4 enzymes of the conjugation system are predominantly nuclear localised, there is clear evidence that the E1 (SAE complex) and E2 (SCE1) are present in significant levels in the cytoplasm suggesting that protein SUMOylation outside the nuclear compartment is mainly driven by SCE1.

The SUMO peptides show considerable variation in their cell type specific patterns. The Arabidopsis genome contains 8 SUMO paralogs.^33,34^ Five of these showed expression in the root tip, specifically SUMO1, SUMO2, SUMO3, SUMO4 and SUMO5, while no expression was found in SUMO6, SUMO7 and SUMO8. SUMO1 and SUMO2 are thought to be the main drivers of SUMOylation in plants as they demonstrate the highest levels of expression,^35^ and the *sumo1-1 sumo2-1* double knock-out is embryo lethal.^18^ SUMO1 and SUMO2 share an 83% amino acid sequence similarity, whereas SUMO1 and SUMO3, SUMO4 and SUMO5 is considerably less (46%, 36% and 37%, respectively). In the processed mature forms SUMO1 and SUMO2 are so similar that MS based analysis for SUMO conjugated targets may not be able to distinguish between these isoforms, causing researchers to assume both genes were broadly expressed and function in a similar manner. Here, we demonstrate a clear distinction in spatial expression domains. SUMO2 displays a broad zone of expression and protein abundance within root tip tissues, while SUMO1 is expressed in phloem pole companion cells starting from the root elongation zone. The contrasting expression pattern of SUMO1 and SUMO2 would enable conjugation to be controlled in a cell-type specific manner under inductive conditions. However, *sumo1-1* and *sumo2-1* single mutants display no obvious phenotypic defects in unstressed conditions,^18,36^ suggesting that they normally act redundantly. Hence, the phenotypes of these single mutant lines await characterisation under a range of stress conditions to reveal SUMO1 and SUMO2 specific roles.

Other SUMOs expressed in roots (3, 4 and 5) likely play more specialised roles in specific tissues. Of these three, only *sumo3* mutants have been studied to date, revealing a small delay in flowering time and a role in plant defence response downstream of salicylic acid.^36^ Unlike SUMO1/2, SUMO3 cannot form poly-SUMO chains and likely has unique target substrates.^37^ Expression of SUMO3 primarily in the epidermis is in accordance with its role in plant defence during root growth in soil. SUMO4 was previously thought not to be expressed in root tissue.^33^ Still, our study reveals weak expression in the late elongation and maturation zone tissues, whilst SUMO5 localises strongly to the stele and meristematic epidermal cells. This is the first observation of SUMO5 tissue and cell-specific expression in plants. The high level of spatial variation in expression between SUMO1-5 suggests SUMO2 drives the main developmental SUMO system as it is widely expressed. In contrast, cell-type specific expression of SUMO1, 3, 4 and 5 allows for fine-tuning of SUMO-mediated responses in a tissue or cell type-specific manner.

Our SUMO Cell Atlas also revealed that DeSI proteases exhibit variable subcellular localisation and cell-type specificity. This class of SUMO protease was first identified in mammalian cells as an evolutionarily distinct group from ULP proteases.^38,39^ DeSIs have isopeptidase activity, deconjugating SUMO from substrates, but cannot process SUMO, whereas the ULP class of proteases catalyze both processes. Interestingly, ULP proteases are widely expressed in different cell-types and localise largely to the nucleus, potentially explained by their critical role in SUMO maturation. In plants, the only non-nuclear localised SUMO protease to be described to date was DeSI3a, which localises to the plasma membrane.^31^ This localization is key to its role in deSUMOylation of the membrane-bound FLS receptor. Our current study reveals that the DeSI family exhibits great variability in subcellular localization, ranging from the cytoplasm, plasma membrane and vacuolar membranes. DeSI3a,b and c are all membrane localised, however, the mVenus signal in DeSI3a displays speckles on the membrane, suggesting highly specific binding to a select group of membrane proteins. This indicates the DeSIs control deSUMOylation of a small subset of substrates locally involved in fine-tuning stress responses. In contrast, ULP proteases appear to play a more general role in SUMO maturation and nuclear processes across multiple root tissues. Collectively, these data show sub-functionalisation of the SUMO signalling machinery often at a sub-cellular scale, providing a basis for individual stress responses to modulate bespoke downstream processes.

SUMOylation represents a vital PTM to modify cellular targets in response to changing environmental signals and stresses. In this study, we tested how the SUMO system in Arabidopsis root tissues reacted to three model environmental stresses; salt, osmotic and biotic signals. Each model stress triggers a distinct response in terms of the SUMO components being regulated and which root tissues respond. For example, each of the stresses regulate SUMO modifiers. Osmotic stress causes levels of SUMO2 and 3 to increase; under salt stress SUMO2 levels go down; whilst after flagellin treatment only SUMO3 levels increase in epidermal cells. Our results also reveal that biotic stress targets much fewer components of the SUMO system compared to salt and osmotic stresses which induce drastic changes in the transcriptional and translational status of the majority of the SUMO system components. Nevertheless, our observations reveal that the SUMOylated proteome in response to different stresses is shaped mostly by SUMO2 and 3. Intriguingly, the SUMO system distinguishes between different abiotic stresses at the tissue-specific level and highly probably at the cell-specific level to generate spatially resolved SUMOylated proteomes. This evidence hints at the possibility of these root tissues being key for enacting adaptive responses mediated by SUMO. Isolating tissue specific SUMOylomes promise to unlock the molecular pathways that integrate root growth and development with its changing environment.

It had previously been documented that SUMO conjugates accumulate upon different stresses in eukaryotes. Our Cell atlas has allowed us to demonstrate that, in plant roots, this increase in SUMO conjugation is primarily driven by increase in protein abundance of SCE1, the E2 conjugating enzyme, rather than E3s. The SCE1 interactome supports this observation as distinct interacting partners were observed for each stress applied. We postulate that specific tissues respond by increasing SCE1 levels and its interactome to shape the SUMOylated proteome upon stress perception. Tissue-specific change in the SUMOylated proteome landscape enacts the adaptive responses we observe – such as hydropatterning in roots.^21^ The Cell atlas provides researchers with new resources to extend this novel observation beyond the root to ascertain if SCE1 is also the principal driver of increased SUMO conjugation in other plant organs, developmental processes and stress responses.

SUMO modification status reflects the balance between the activity of the E1-E4 conjugation system and deconjugation controlled by ULP and DESI classes of SUMO proteases. Gene families of these SUMO proteases have expanded as plants evolved to adapt to diverse environmental conditions on land, whilst numbers of genes encoding the E1-E4 conjugation system remained unexpanded.^9^ Our Cell Atlas revealed unique combinations of SUMO protease expression changes in response to each of the 3 stresses. For example, following flg22 elicitor treatment, OTS1 and DESI3A was specifically turned over, while ESD4, DESI2B and 3C increased in abundance. This observation suggests that the flg22 dependant SUMOylome is shaped by the increase in SCE1 dependant conjugation and ESD4, DESI2B and 3C deconjugation, followed by a decrease in OTS1 and DESI3A deconjugation. In contrast, salt stress reduced OTS1 and 2, DESI1 and DESI4A abundance, whilst SPF1 and ELS1 increased in a tissue-specific manner. This insight into the tissue/cell-specific SUMO “coding” system opens up new avenues for identifying stress-specific SUMOylated targets. There is correlative evidence for the neo-functionalization and gene expansion of SUMO proteases, through fusion of conserved catalytic domains with divergent sequences with the emergence of adaptive traits in the plant lineage.^9^ Given that we did not find a component level of specificity to different stresses within the conjugation machinery compared to the SUMO proteases, we postulate that target specificity in plants lies mainly in the SUMO protease gene family of their SUMO system.

## Resource availability

We provide a resource for all researchers working on plant SUMOylation which is located on the ePlant server (https://bar.utoronto.ca/~nprovart/SUMO_Map/). This interweb interface visualises the expression, and localisation of all SUMO components in unstressed and stressed plants demonstrated as a heatmap across root tissues. Seeds of all transgenic lines have been deposited in the Nottingham Arabidopsis Stock Centre. Additionally, through the pMDS1 and pMDS2 vectors we provide tools where others can simultaneously visualize gene expression and protein localisation of their gene of interest. Further information and requests for resources and reagents should be directed to and will be fulfilled by the lead contact Dr. Ari Sadanandom;

## Acknowledgments

We thank UKRI BBSRC for the strategic Longer Larger Grant; SUMO code (BB/V003534/1) to, AS, MJB, AJ, KSL, MdL, AB, LB and RB. We thank David Salt for developing the SUMO code proposal.

## Author contributions

AS, MJB, AJ, KSL, MdL, AB and RB conceived the ideas, analysed data, wrote the manuscript. JB, ShG, DR, conceived the ideas, performed most of the experiments, analysed data and help write the paper, KI, LC, SrG performed experiments and analysed data, ES, LB and KSO, analysed data and help generate figures. LB, DW and JA helped supervise experimental work and conceptualise the main themes of the manuscript.

## Supplemental Figure Legends

**Figure S1.**
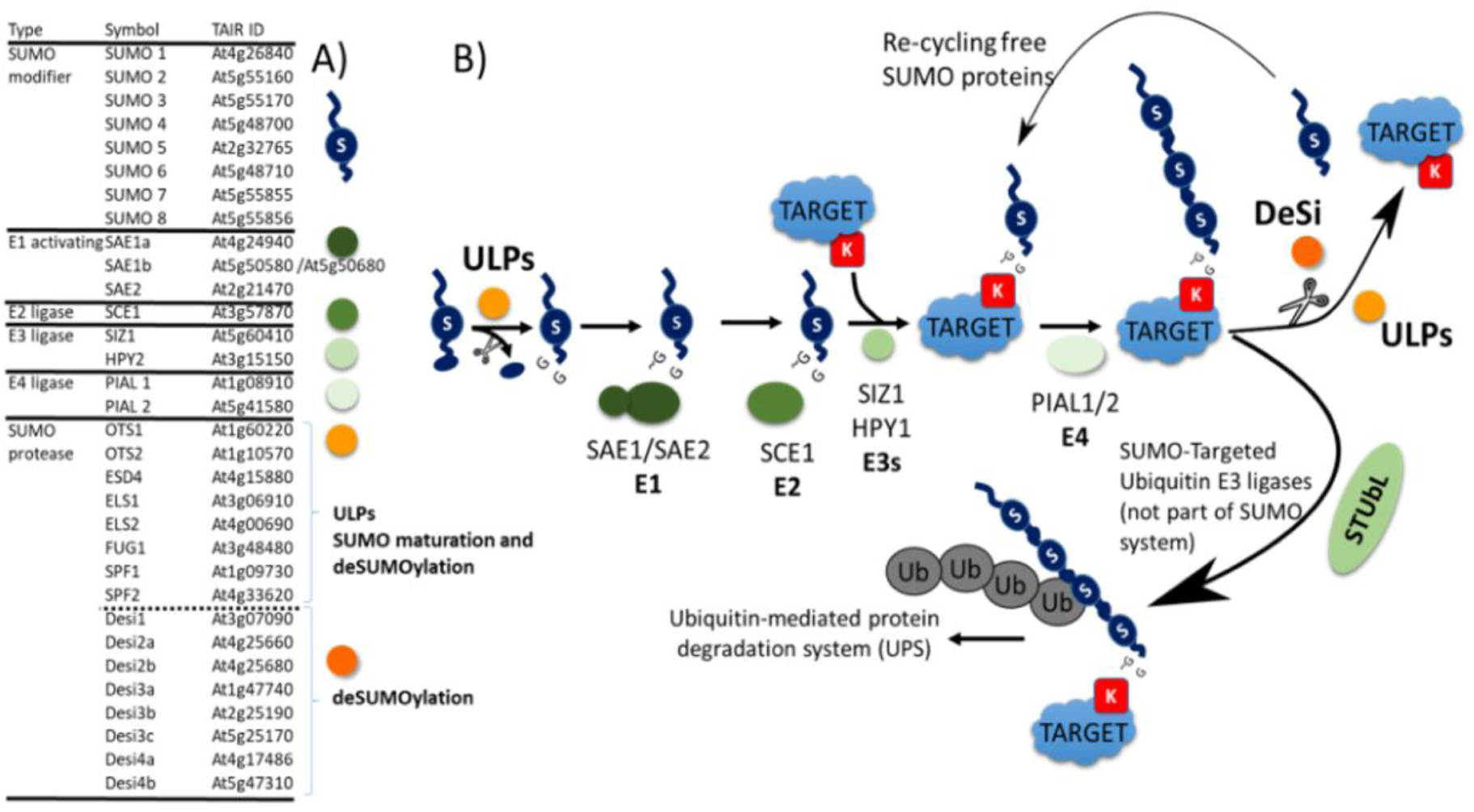
**A)** Table of known SUMO machinery genes in *Arabidopsis thaliana*; **B)** Schematic of SUMO System composition and functions in *Arabidopsis thaliana*. The ULP class of SUMO proteases process the precursor SUMO modifier at the C-terminus to expose the diglycine C-terminal end (-GG). The E1 (2 subunits; SAE1/SAE2) initiates the enzymatic conjugation cascade of the SUMO modifier, resulting in the conjugation of the SUMO modifier to the lysine (K) residues on substrates. ULP and DeSi class of SUMO proteases cleave SUMO off-targets. ULP= UBL-specific proteases, (UBL= Ubiquitin-like family protein modifiers); DeSi = DeSumoylating Isopeptidase; STUbl =SUMO-targeted ubiquitin ligases; Ub=Ubiquitin.

**Figure S2.**
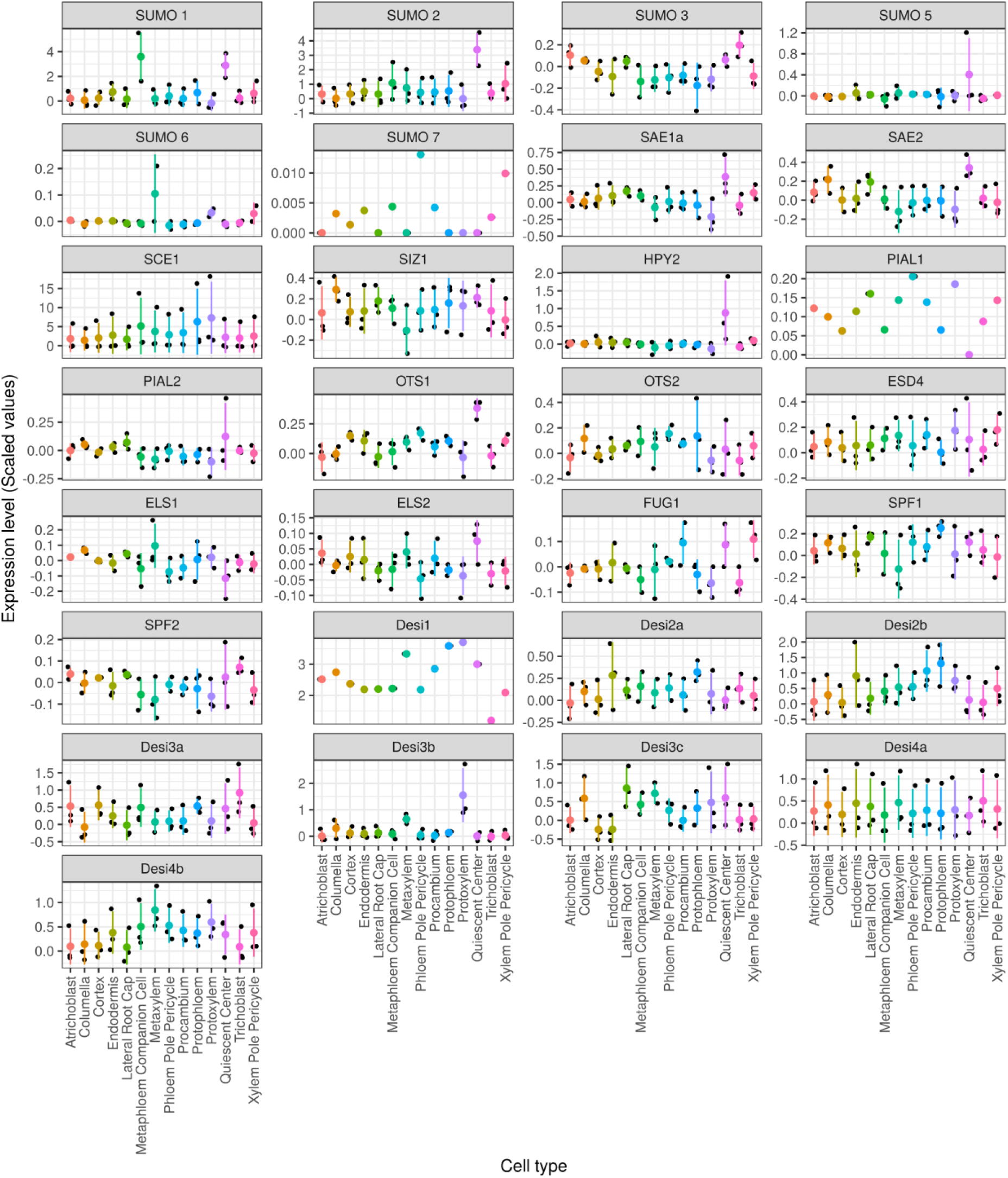
Expression levels of candidate genes representing the Arabidopsis SUMO machinery across various root cell types. The figure shows the expression levels of SUMO system genes from scRNAseq experiments across multiple cell types of the Arabidopsis roots. Each plot’s coloured bold dot and line represent the mean and standard deviation. Black dots for each cell type are the scaled and transformed expression values from 3 scRNAseq datasets.

**Figure S3.**
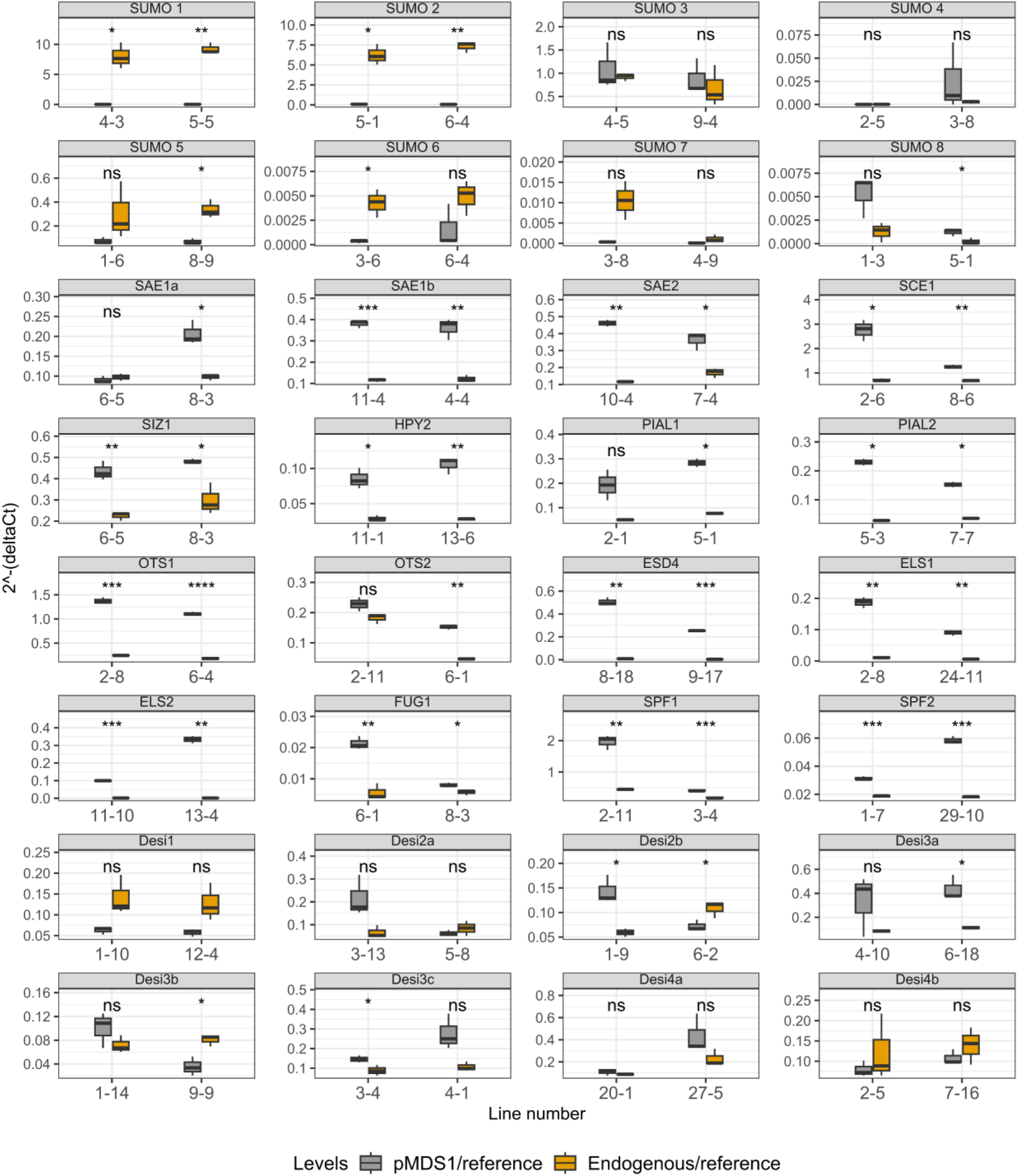
Expression levels of candidate genes representing the Arabidopsis SUMO machinery across independent transgenic lines developed for this study. Quantitative real-time PCR analysis was performed to assess the gene expression levels of the transgene (incorporated in the pMDS vector) in comparison to its endogenous counterpart in the transgenic lines of All 32 SUMO components. The plotted values show the 2^-(ΔCt) values for the endogenous gene vs pMDS1-fusion gene expression against the respective reference genes. The statistical significance of the difference between pMDS1 and endogenous gene as indicated by Student’s t-test is shown by the asterisk (*). ‘****’, ‘***’, ‘**’, ‘*’, and ‘ns’ represent a p-value cut-off of value < 0.0001, < 0.001, < 0.01, < 0.05, > 0.05, respectively.

**Figure S4.**
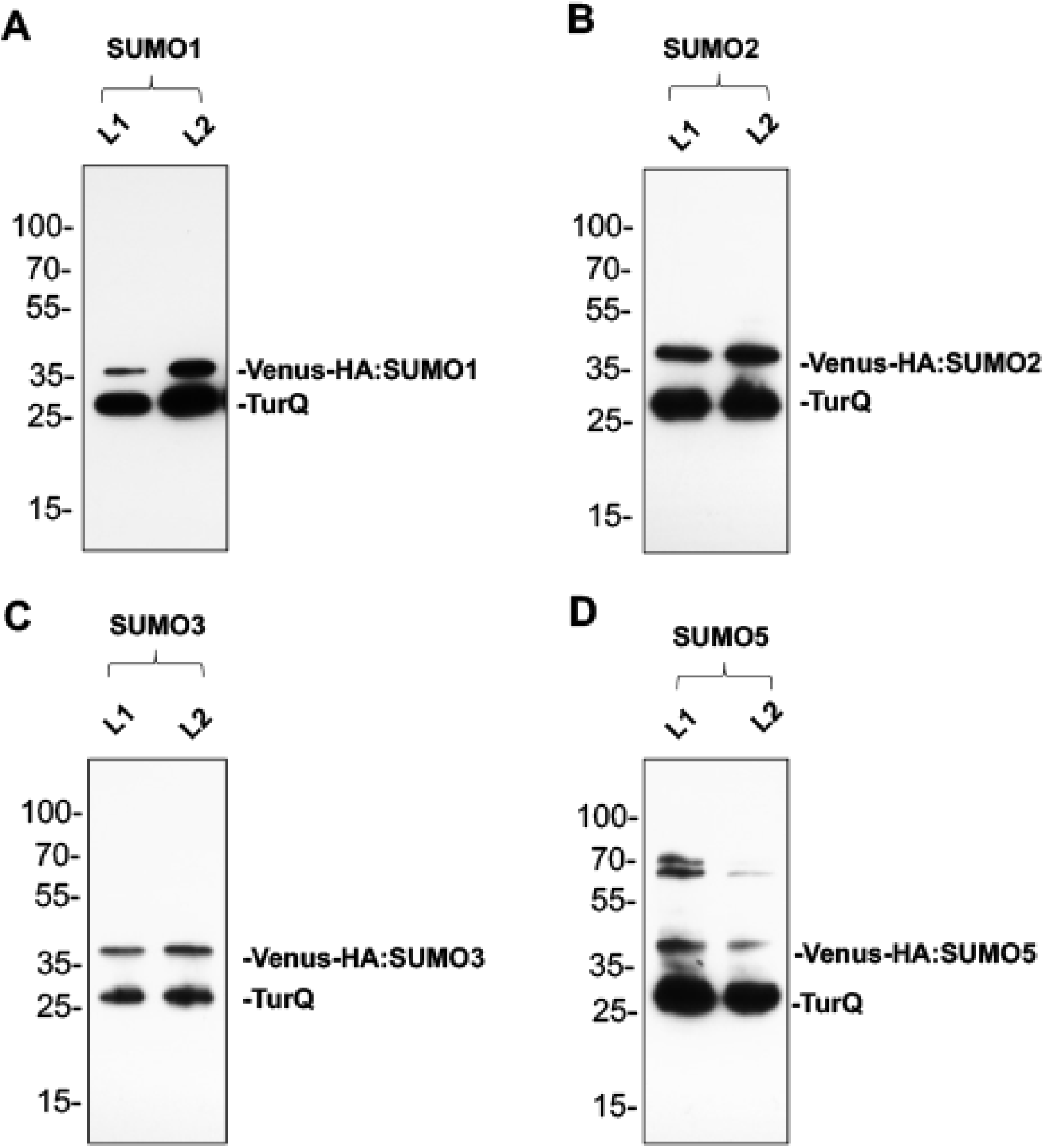
Immunoblots of the SUMO modifier expressing pMDS2 lines indicating efficient ribosomal skipping induced by the 2A peptides. **(A)** SUMO1 **(B)** SUMO2 **(C)** SUMO3 **(D)** SUMO5. Bands corresponding to mTurquoise are labelled as mTurq.

**Figure S5.**
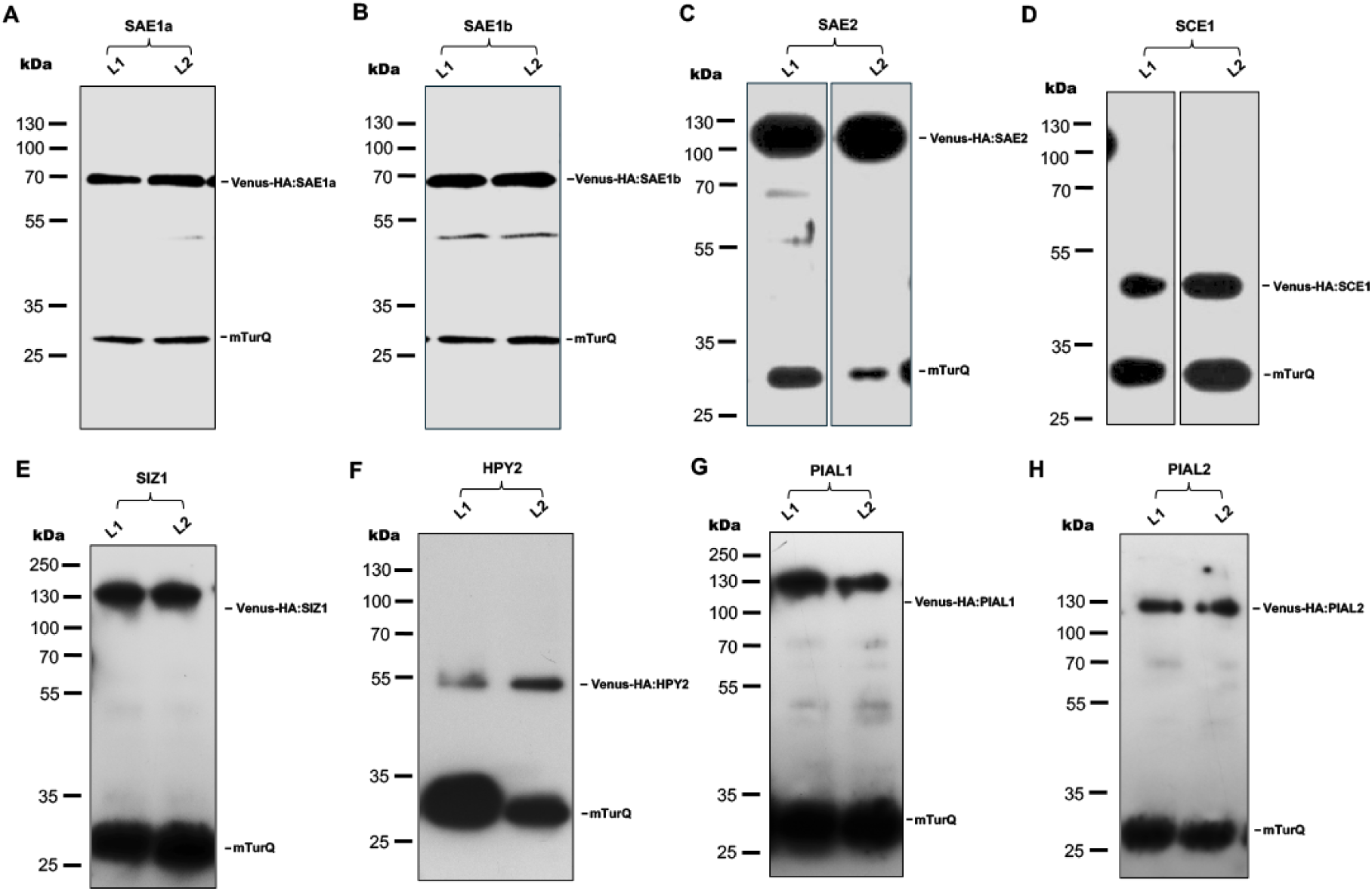
Immunoblots of the E1-E4 enzyme expressing pMDS1 lines indicating efficient ribosomal skipping induced by the 2A peptides. **(A)** SAE1a **(B)** SAE1b **(C)** SAE2 **(D)** SCE1 **(E)** SIZ1 **(F)** HPY2 **(G)** PIAL1 (**H)** PIAL2. Bands corresponding to mTurquoise are labelled as mTurq.

**Figure S6.**
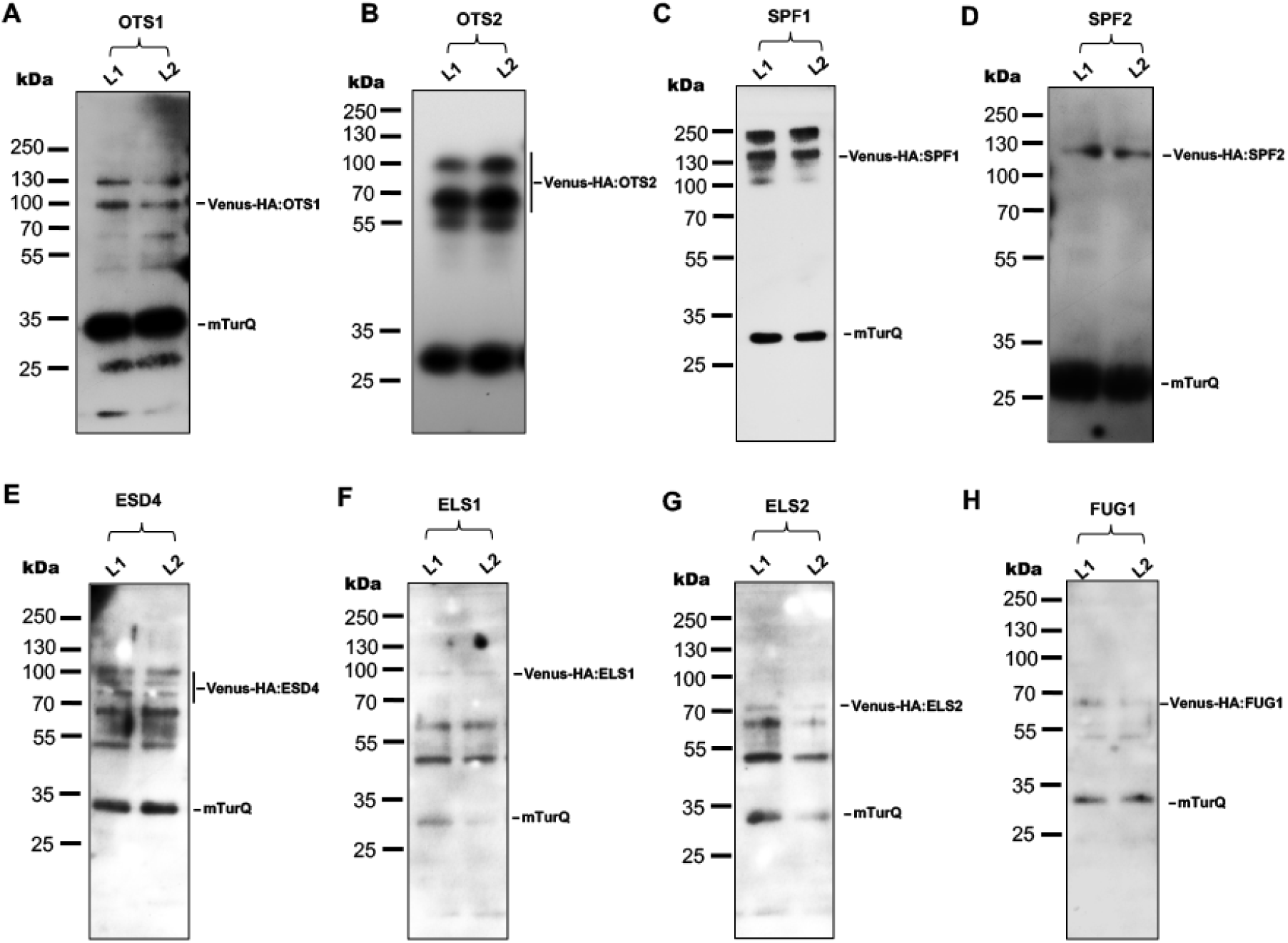
Immunoblots of the ULP SUMO protease expressing pMDS1 lines indicating efficient ribosomal skipping induced by the 2A peptides. (A) **OTS1** (B) **OTS2** (C) **SPF1** (D) SPF2 **(E)** ESD4 **(F)** ELS1 **(G)** ELS2 (**H)** FUG1. Bands corresponding to mTurquoise are labelled as mTurq.

**Figure S7.**
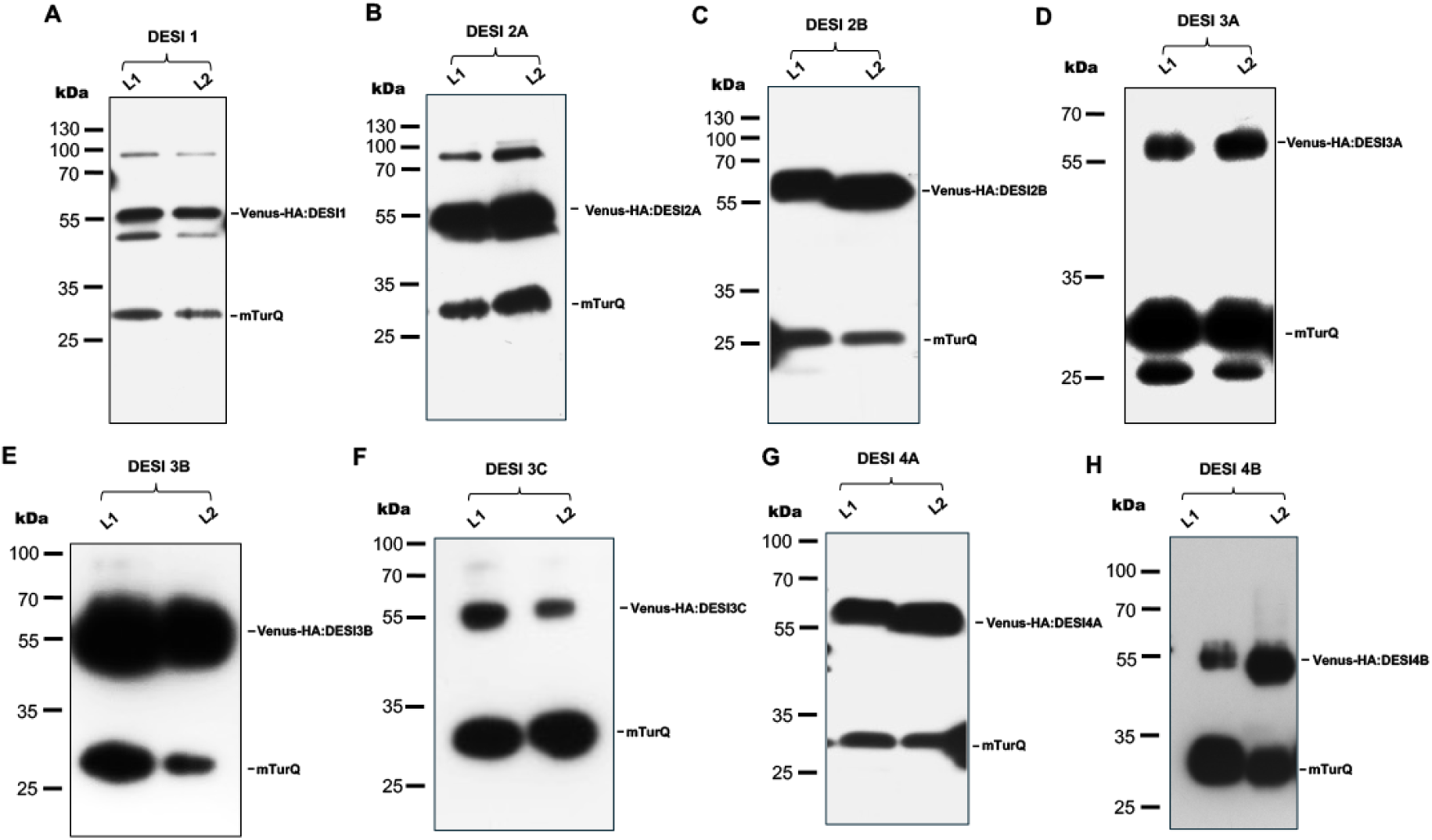
Immunoblots of the DESI SUMO protease expressing pMDS1 lines indicating efficient ribosomal skipping induced by the 2A peptides. **(A)** DESI1 and DESI 2A **(B)** DESI 2B **(C)** DESI 3A **(D)** DESI 3B (**E)** DESI 3C **(F)** DESI 4A **(G)** DESI 4B. Bands corresponding to mTurquoise are labelled as mTurq.

**Figure S8.**
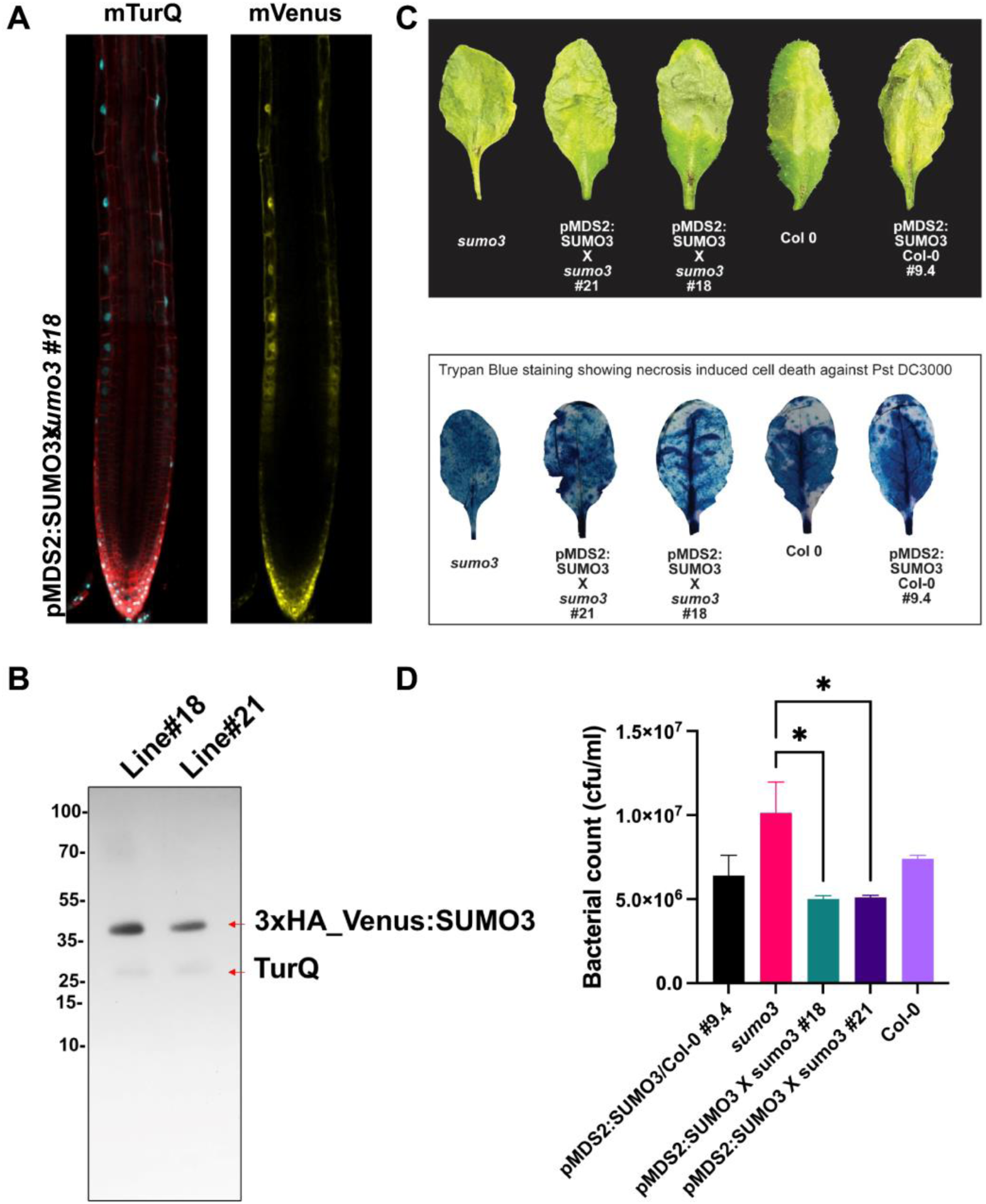
Complementation assay of *sumo3-1* mutant using pMDS2_SUMO3. (A) Confocal image of pMDS1:SUMO3 x *sumo3-1* showing gene expression (mTurQ) and protein level (mVenus) in epidermis. (B) Western blot analysis showing confirmation of complementation in pMDS1:SUMO3 x *sumo3-1* (Line 18 and 21). The blots were probed with α-GFP (C) Disease symptoms in leaf tissues infected with *Pseudomonas syringae pv. tomato DC3000* (upper panel), and trypan blue staining showing cell death against *Pst.* DC3000 (lower panel). (D) Bacterial count from 4-week-old Arabidopsis plants infected with virulent *Pst.* DC3000 at 3dpi. Error bars show standard error of three biological replicates. The asterisk indicates significant difference at p-value ≤ 0.05.

**Figure S9.**
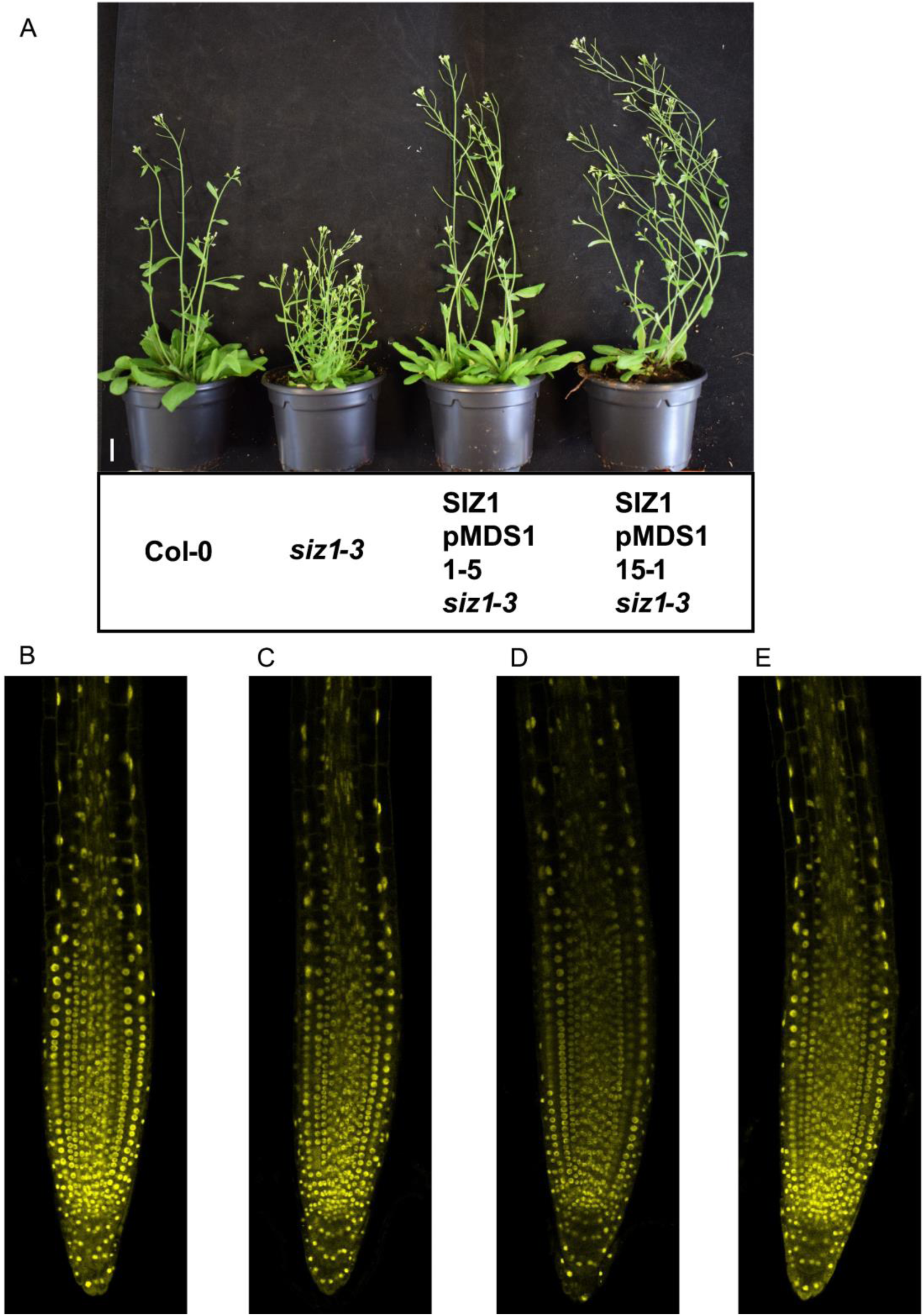
Complementation assay of *siz1-3* mutant using pMDS1_SIZ1. **(A)** Complementation of *siz1-3* mutant using the SIZ1 pMDS1 construct restores shoot growth**. (B and C)** mVenus signal of SIZ1 pMDS1 in Col-0. **(D and E)** mVenus signal of SIZ1 pMDS1 in *siz1-3*Scale bars represent 1cm **(A)** and 100μm **(B-E)**.

**Figure S10.**
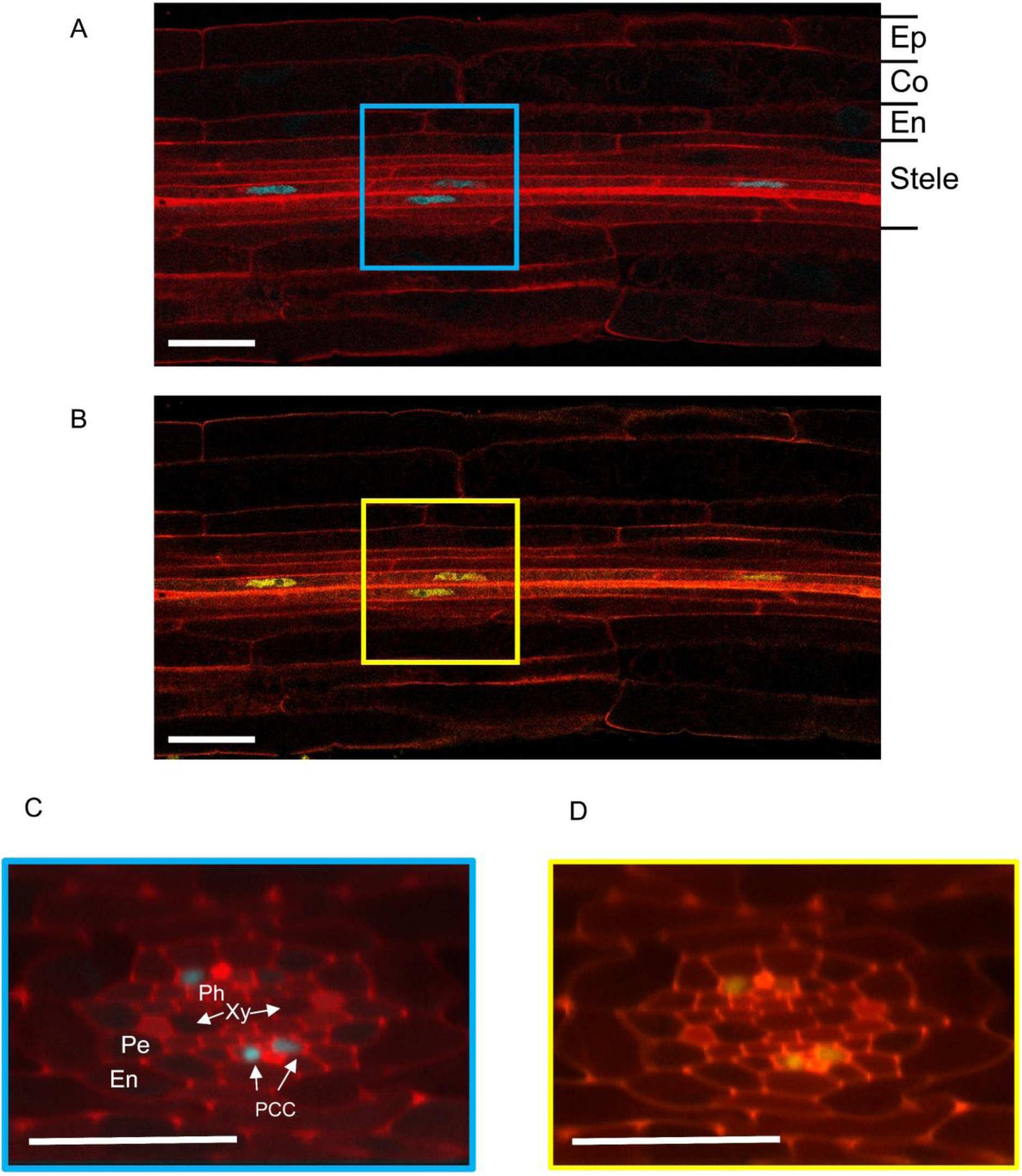
SUMO1 root cross sections indicated expression and localisation in the phloem pole companion cells. **(A and B)** Longitudinal sections of the localization of SUMO1 expression (A) and protein localization (B) in the root tip stele. Note the indication of the root cell type radial layout including, Epidermis (Ep), Cortex (Co), Endodermis (En) and Stele. **(C and D)** cross sections of localisation of SUMO1 expression (C) and protein localization (B) in the phloem pole companion cells. Note the indication of the Endodermis (En), Pericycle (Pe), Xylem (Xy), Phloem (Ph) and Phloem Pole Companion cells (PPC). Scale bar represent 25μm.

**Figure S11.**
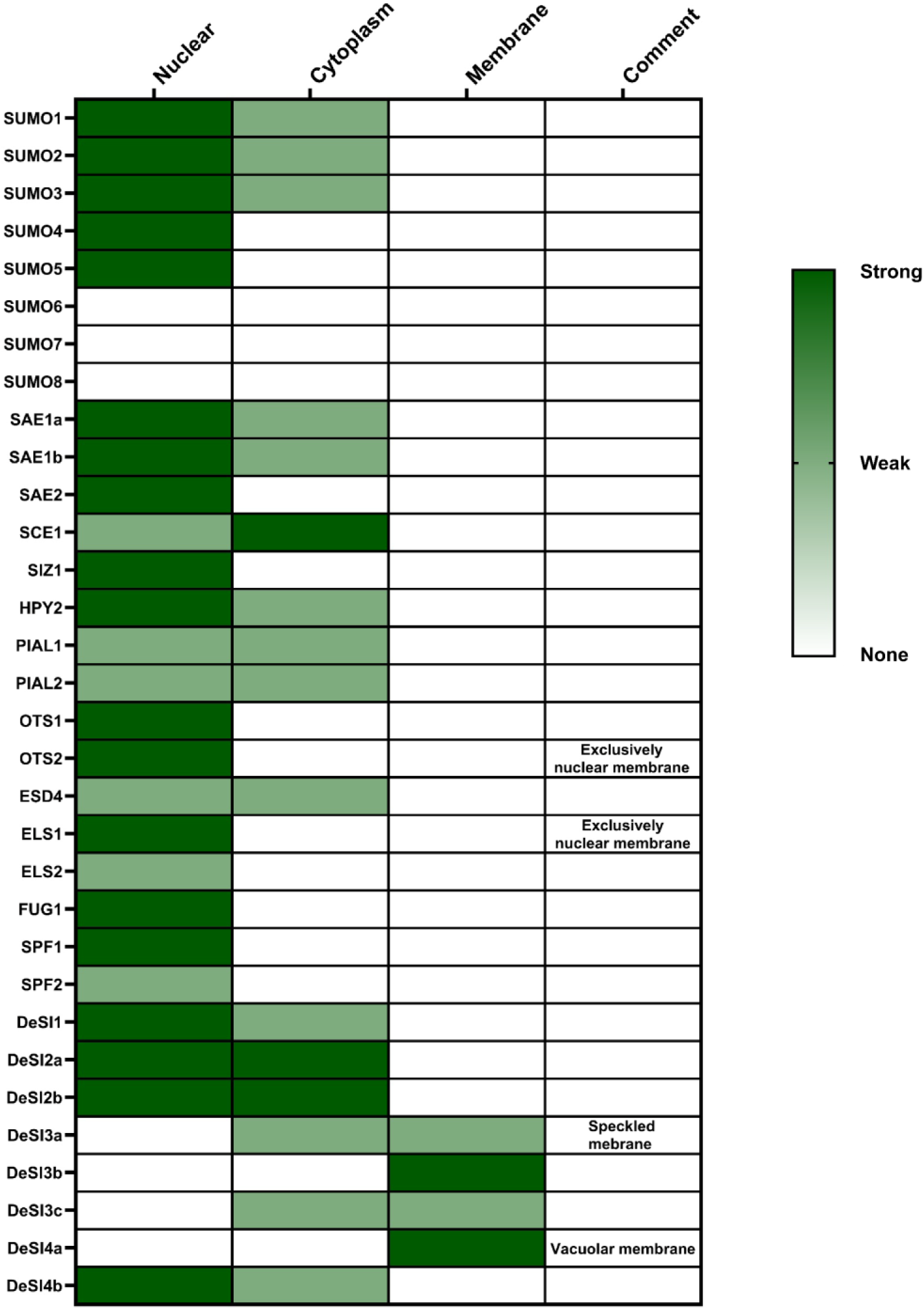
Subcellular location of all SUMO components in *A. thaliana* based on imaging data in the root tip. The columns in the heatmap describe different subcellular locations including nuclear, cytoplasmic and membrane. The last column is used in case protein location is more specific than the previous defined groups. The colours define the strength of localisation in a certain organelle, where white defines no protein, pale green is weakly localised to this organelle and dark green is strong localisation in this organelle. For example, SUMO1 is strongly localized in the nuclear with only some expression in the cytoplasm.

**Figure S12.**
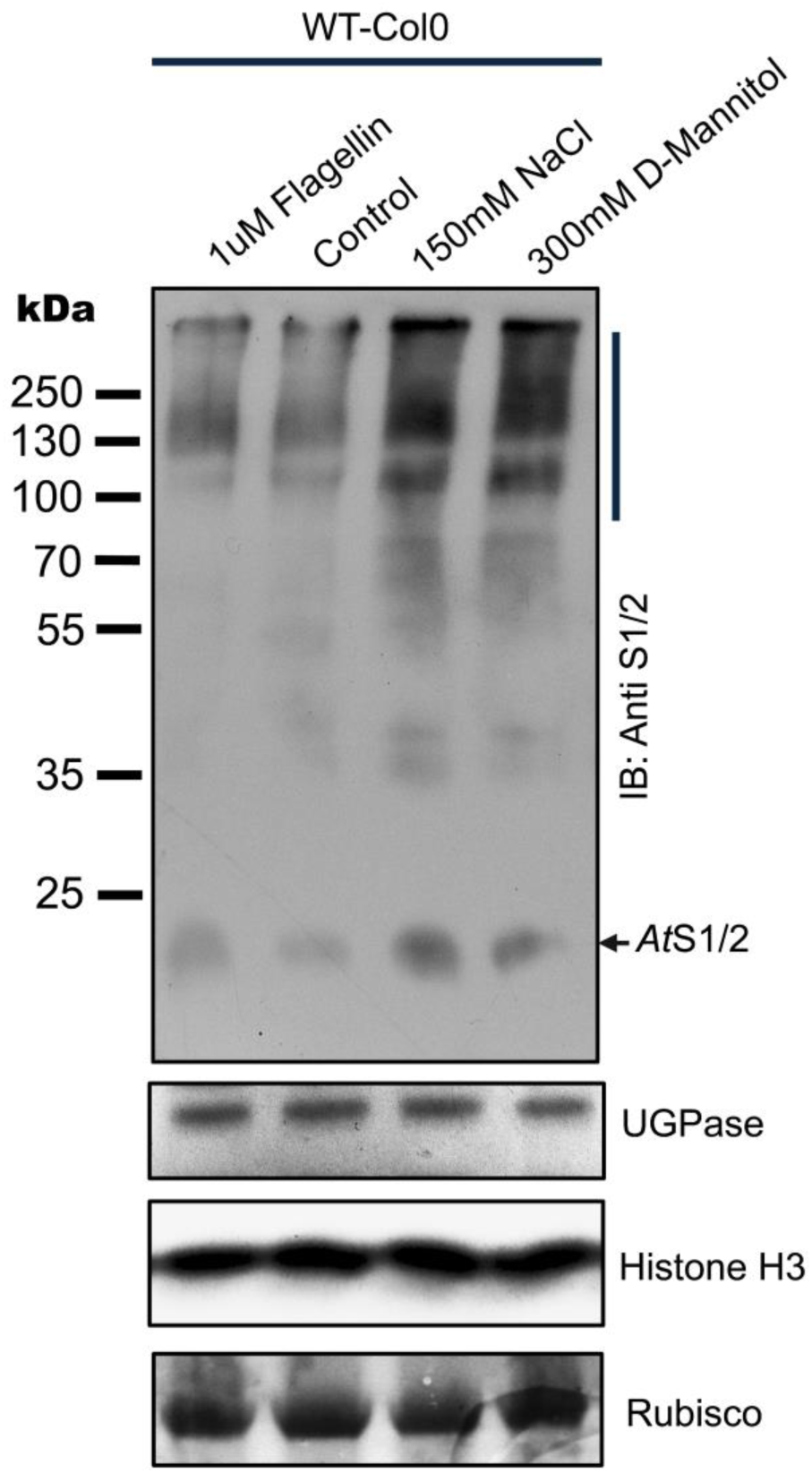
SUMO1/2 conjugates accumulate with osmotic stress. Immunoblot (IB) analysis of total protein (10 mg loaded for each lane) derived from wild-type Col-0 *Arabidopsis* seedlings grown for 12 d under long-day conditions and then subjected to the indicated stresses for 3 hours. Filters were probed with anti-SUMO1/2 (anti At S1/2) antibodies (J46). The vertical bar shows the laddering effect produced by the increased accumulation of SUMO1/2 conjugates. The arrowhead indicates free (nonconjugated) SUMO1/2. The bottom panels show immunostaining of UGPase and Ponceau staining of ribulose-1,5-bis-phosphate carboxylase/oxygenase (Rubisco) small subunit (RbscS), which served as loading controls. The experiment was repeated three times with similar results. Panel **A, B**, and **C** are different exposures of the same filter to highlight the laddering effect produced by the increased accumulation of SUMO1/2 conjugates.

**Figure S13.**
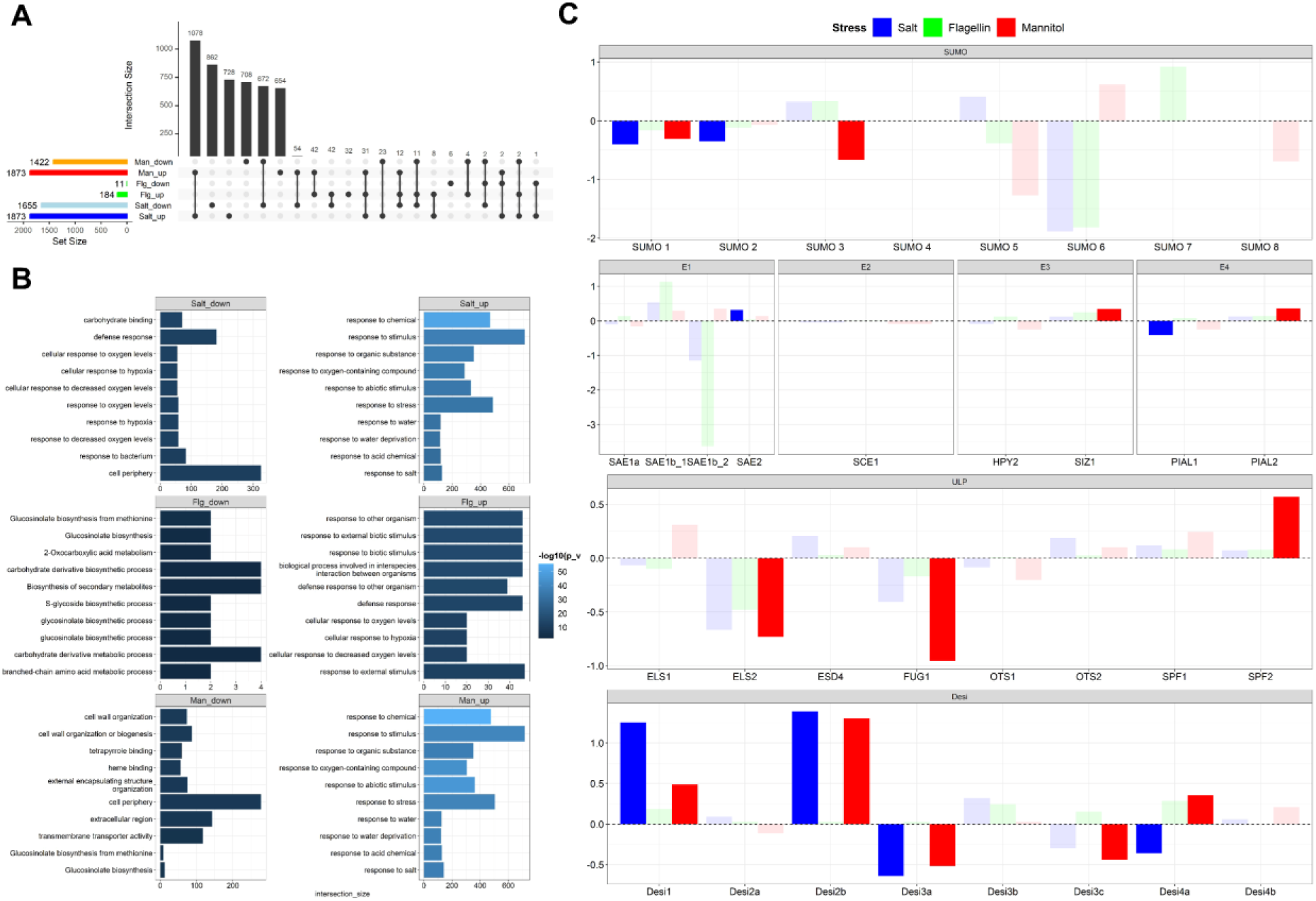
Transcriptomic profiles of the SUMOylation machinery under salt, flagellin and mannitol treatments. **A)** Summary of the DEGs in response to the stresses. The number of the up/down–regulated DEGs are shown in the sidebars. The number of common DEGs between the treatments (connecting dots) is shown in the black bars. **B)** Top 10 GO terms of the DEGs. The GO terms are arranged by most significant p_value **C)** Transcripts expression of SUMOylation genes. The Y-axis is Log2-fold change values. Colours represent the treatments; salt (blue), flagellin (green), and mannitol (red). Solid colours are the DEG significant at p_value < 0.05.

**Figure S14.**
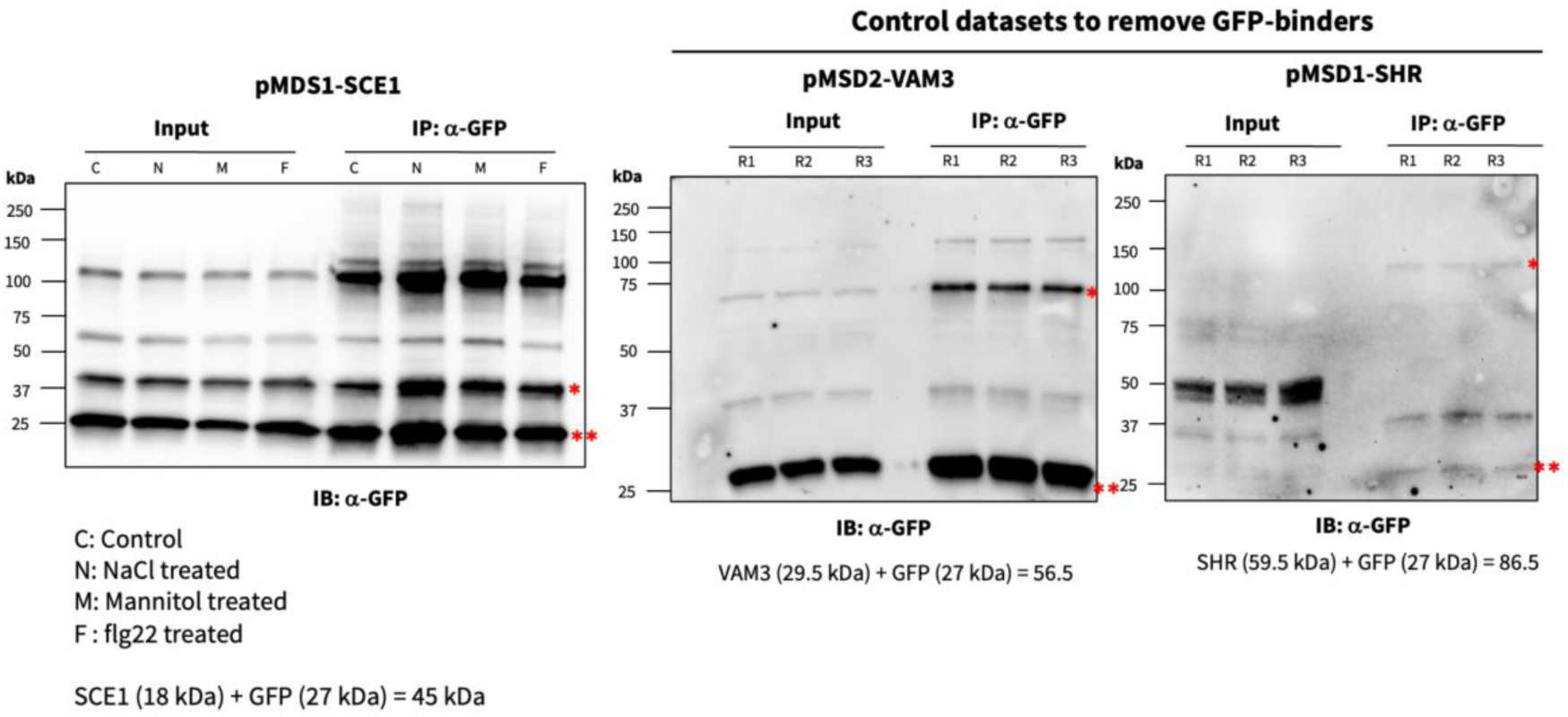
Immunoblot analysis of SCE1, VAM3, and SHR following immunoaffinity enrichment using anti-GFP magnetic beads. Single red asterisk indicates the Venus -3X HA fusion of the protein of interest (SCE1, VAM3 and SHR); while double red asterisk indicates mTurquoise band.

**Figure S15.**
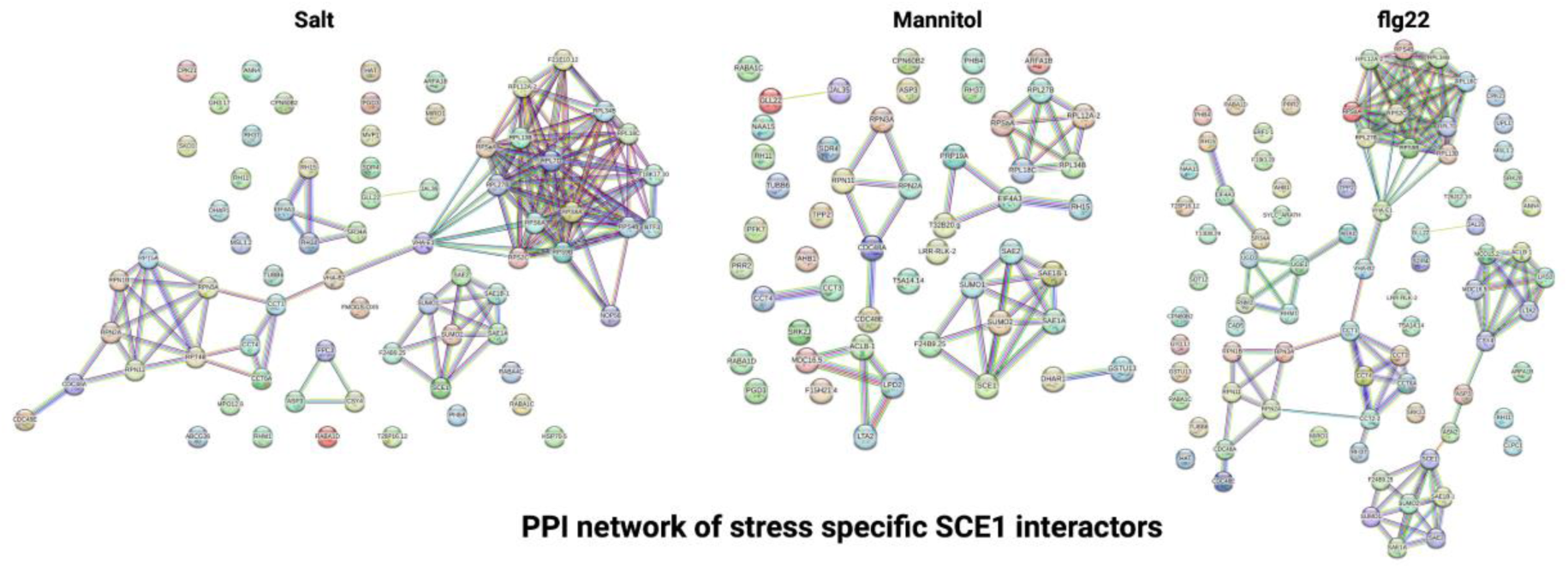
Visualization of protein-protein interaction (PPI) networks for SCE1 interactors under stress conditions (salt, mannitol, and flg22) generated using the STRING database (https://string-db.org).

## Star Methods

### Contact for reagent and resource sharing

Further information and requests for reagents and resources may be directed to, and will be fulfilled by the corresponding author, Dr. Ari Sadanandom.

### Materials availability

Destination plasmids generated in this study have been deposited to Addgene. The Arabidopsis lines generated in this study have been deposited to NASC. All other lines are available on request from the corresponding author (Tables S1).

### Data and code availability

Expression and protein abundance data of all 32 SUMO components and their response to stresses is available via ePlant.

Bulk RNA seq data available from NCIB GEO database (www.ncbi.nlm.nih.gov/geo/query/acc.cgi?acc=GSE275006).

Microscopy data used for quantification is deposited on Mendeley Data (DOI: 10.17632/fgjvs8589d.1).

### Experimental model and subject details

All lines generated in this study were in the Arabidopsis thaliana Col-0 background. For complementation lines the following mutant backgrounds were used: *sum3-1* (SALK_123673C) and *siz1-3* (SALK_034008). The seedlings were grown at 21°C. The growth conditions for individual experiments are listed in the method details section.

## Method details

### Generating transgenic material

The generation of a polycistronic vectors (pMDS1 and pMDS2) that enables simultaneous monitoring of both the transcription and translation of a gene via confocal imaging and western blot, involved several cloning steps:

For the pMDS1 (C-terminal fusion), first, the Venus fluorescence coding sequence was PCR amplified (primers: 863/864 and template DR5:Venus:N7) and inserted into a pGWB13 vector containing a 3xHA tag and Xba/PacI sites using Hot Fusion cloning. From this vector, the VENUS:3xHA sequence was amplified by PCR with the addition of a long glycine-serine linker sequence on the 5’ end. This PCR product was cloned into the HindIII site of pMCY2, a vector containing mCherry for seed selection and cell imaging marker. Next, the 2A:mTurquoise (primers 868/79) and N7 (primers: 869/ 870 and template DR5:Venus:N7) coding sequences were separately amplified by PCR and simultaneously cloned into the pMCY2 vector containing GSlinker:Venus:3xHA sequence using Hot Fusion into the Bsu36I site. The mTurquoise was re-educated to remove a SapI restriction site of its sequence (primers: 874, 875, 876 and 877) and reinserted into the vector using HindIII digestion and Hot Fusion reaction. The final plasmid containing GSlinker:Venus:3xHA:2A:mTurquoise:N7 was named pMDS1

For generation of the pMDS2 (N-terminal fusion), we amplified 3xHA from pMDS1 using primers 893 and 867, followed by insertion via hot fusion of the gel-purified fragment into a HindIII-digested pMCY2. The resulting vector was then linearized at a BSU36I site (introduced by primer 867) to enable insertion of the GSG-Linker:Venus fragment, which was amplified from pMDS1 using primers 880 and 881. Subsequently, this construct was linearized at the BSU36I site (introduced by 881) to allow insertion of the 2A peptide-mTurquoise fragment, which was amplified from pMDS1 using primers 886, 888, and 870. The final pMDS2 plasmid contains a single BSU36I site that can be used to generate an N-terminal 3xHA:GSG:Venus fusion protein.

### Making of the pMDS1and pMDS2 contructs

Gene specific primers were designed to clone the promoter (∼2kb or up to the previous ORF in CIS) and genomic region of the 32 SUMO components into the pMDS1 vector using the Hot Fusion approach,^40^ (Table S2) at SapI sites for pMDS1 and SapI and Bsu36I for pMDS2. Only for SUMO1 construct a shorter promoter was used (add length) as the 2kb promoter did not express. The final plasmid was transformed in *Agrobacterium tumefaciens GV3101* strain containing pSOUP helper plasmid by electroporation and then transformed into *Arabidopsis thaliana* ecotype Col-0 by the floral dip method.

### Plant materials and growth conditions

Seeds were surface sterilized for 4 min in 30% (v/v) bleach containing 0.005% Triton X-100. Followed by five washes with sterile water and then stratified at 4 °C for 48 h in the dark. Seeds were germinated on media containing half-strength Murashige and Skoog (MS) media (Sigma), 0.4% sucrose and 1% Bacto-agar at pH 5.7. Seedlings were grown vertically for 6 days under day temperature of 23 °C and night temperature of 18 °C with a 16-h photoperiod (150 μmol m−2 s−1). All transgenic material and constructs are listed in Table S1.

### Confocal settings SUMO Cell Atlas (Figure 2)

T3 plants of the different SUMO components were imaged using a Leica SP8 confocal microscope (Leica Microsystems) using a 20x objective and sequential scanning using TileScan with automated merging. The mTurqouise signal was excited at 422nm with a 422 diode and light captured between 452-505nm using a (Hybrid GaAsp/APD (HyD)). For mVenus signal, we used an Argon laser at 516nm and light captured between 520 and 570 using a (Hybrid GaAsp/APD (HyD)). mCherry signal was excited with a 561nm DPSS laser and light between 590 and 731 nm was collected using a Two Photo Multipliers Tube (PMT) detector.

To understand the relative expression of these variably expressed components three laser powers for both the 422 diode and the argon laser were used: 2%, 14% and 26%, while the Gain stayed the same (150 V).

To create the split images, TileScan images were used and roots straightened with Fiji’s Straighten function. This allowed us to make clear splits through the longitudinal middle of each root. One half of the root was then chosen, in which the nuclei line up in the different cell-type in mTurquoise channel. This half was then split into 3 channels and afterwards mTurqouise and mCherry were fused back together. One half would then be flipped horizontally to mirror the other half.

### Confocal settings for stress atlas (Figure 3-5)

For imaging the SUMO stress atlas, the same confocal settings as for Figure 2 were used, but instead of imaging in three different laser powers, only the optimum was chosen per gene, and this was not changed during the experiment and repeats. Experiments were repeated twice with eight roots per treatment. The median experiment out of three and the four median roots within that experiment were chosen for detailed tissue specific analysis. This was based on whole image fluorescence of both the mTurqouise and mVenus channel.

### Mannitol stress

6-day old *Arabidopsis* seedlings were transferred from 1/2MS plates to plates either containing 300mM Mannitol or Mock plates. After 3 hours, seedlings were mounted on a slide and scanned on the confocal microscope.

### Salt stress

Five-day-old Arabidopsis seedlings grown on 1/2 MS medium were transferred to a setup containing liquid 1/2 MS supplemented with 150mM NaCl, ensuring that only the roots were exposed to the salt treatment while the aerial parts remained untreated. After 3 hours of salt exposure, the seedlings were mounted on slides and imaged using a confocal microscope. A parallel experimental setup containing only liquid 1/2 MS was used as a reference control.

### Flagellin stress

Five-day-old Arabidopsis seedlings grown on 1/2 MS medium were transferred to a setup containing liquid 1/2 MS supplemented with 1uM Flagellin 22, ensuring that only the roots were exposed to the salt treatment while the aerial parts remained untreated. After 3 hours of salt exposure, the seedlings were mounted on slides and imaged using a confocal microscope. A parallel experimental setup containing only liquid 1/2 MS was used as a reference control.

### RNA Isolation and Quantitative RT-PCR

For the quantification of the expression level of endogenous and exogenous genes in the reporter lines, roots of 6-day old seedlings (∼100 roots per sample) were harvested. Total RNA was extracted using RNeasy Plant Mini Kit (Qiagen). Total level of RNA was then measured using NanoDropTM 1000 Spectrophotometer (Thermo Scientific) and normalised before cDNA synthesis. cDNA was synthesised using Invitrogen SuperScript® II Reverse Transcriptase. Forward primers for qRT-PCR of endogenous gene expression were designed, when possible, to span an exon junction as to avoid amplification of contaminated genomic DNA. The reverse primer was designed by using the 3’ UTR region as this was not included in the construct design. Exogenous gene expression primers used the Venus-N7 region as indication of transgene expression level. A full list of primers is shown in Table S3. qRT PCR was performed using the Analytik Jena qTower³ 84 G Real-Time PCR System (Nottingham) and Qiagen Rotor gene Q system (Durham). Actin2 (AT3G18780) was used as housekeeping gene in case of high expression. Low gene expression used PP2A as housekeeping gene. Three biological replicates and three technical replicates were used in this experiment. The difference (ΔCt) in the expression of trans-gene or the endogenous gene was calculated with reference to the housekeeping gene and is plotted as 2^-(ΔCt). The significance between expression of trans- and endogenous genes was calculated by a T-test, P-value < 0.05.

### RNA isolation (for bulk RNA seq)

For the RNA-seq stress dataset, roots of 6-day old seedlings (∼120 roots per sample) were harvested 3 hours after stress treatment. Total RNA was extracted using RNeasy Plant Mini Kit (Qiagen). Extracted RNA was send to Novogene Europe where a poly-A non-stranded library prep was done followed by Illumina PE150 sequencing to 3GB depth conducted on the NovaSeq 6000 S4 platform.

### Western Blotting

*Arabidopsis thaliana* seedlings of the appropriate reporter line were treated with either salt, mannitol and flagellin for 3 hours after which 50 roots were harvested and flash frozen in liquid nitrogen. Total protein was extracted using Laemmli buffer containing: 50mM Tris-HCl (pH6.8), 2% SDS, 10% glycerol, 147mM B-mercaptoethanol, 12.5mM EDTA, 2mg bromophenol blue made up in deionized water. After protein extraction, the samples were separated by SDS/Page and blotted onto PVDF membrane. PVDF membrane was then blocked with 5% (w/v) milk in TBST. The membrane was then either probed with anti-HA antibody (3F10, Roche) or anti-UGPase (Agrisera), followed by anti-Rat HRP(Abcam) or anti-Rabbit HRP (Abcam). The membranes were treated with SuperSignal™ West Pico PLUS Chemiluminescent Substrate (Thermo Scientific), and luminescence detected using an x-ray cassette and fixer and developer.

### Immunoprecipitation assay

To assess the level of efficiency in mediating ribosomal skipping by the 2A peptide in the pMDS1/pMDS2 vector system, we performed immunoprecipitation assays employing anti-GFP pulldowns. Arabidopsis thaliana seedlings were grown in Gamborgs B5 media with added B5 vitamin and sucrose. Roots were harvested and total protein extracted using the following extraction buffer: 50 mM Tris-HCl (pH 8.5), 150 mM NaCl, 1mM EDTA, 0.1% [v/v] SDS, 0.5% [w/v] Sodium deoxycholate, 1.0 % [v/v] Igepal CA-630 (Nonidet P40), 50mM KCl , 50mM NEM (N-ethyl maleimide) and protease inhibitor cocktail (one tablet per 10ml of buffer). Total protein extract was incubated with anti-GFP beads (Miltenyi Biotech) on a rotator at 4°C at 20-25 rpm speed for 30 minutes. Afterwards the extract was loaded into a micro column (Miltenyi Biotech) in a magnetic stand. Beads were then washed 4 times with ice cold IP extraction buffer buffer. 80μl of pre-heated (98°C) Laemmli buffer was used to elute the protein. The eluted immunoprecipitated fraction was finally probed with anti-GFP antibody (Abcam, ab 6556) to detect the various forms of the target fusion protein and the transcriptional reporter mTurquoise. Efficient ribosomal skipping induced by the 2A peptide will result in two polypeptides that include: POI-GFP-3XHA and N7-mTurquose, where POI stands for <u>P</u>rotein <u>O</u>f Interest.

### Immunoprecipitation (IP) and LC-MS/MS Analysis

Proteins were extracted from the root tissues of pMDS1-SCE1 plants using three biological replicates for immunoprecipitation experiments. As a negative control, root tissues from pMDS1-VAM3 and pMDS1-SHR were utilized to account for non-specific binding to GFP, as well as to exclude interactions with SUMO pathway-unrelated proteins, such as VAM3 and SHR. The protein extraction buffer (EB) contained 50 mM Tris-Cl (pH 7.5), 125 mM NaCl, 1.5 mM MgCl2, 1 mM EDTA, 5% (v/v) glycerol, 0.1% (v/v) Tween-20, 0.2% (v/v) Igepal CA-630, and 0.5% (w/v) digitonin, supplemented with freshly added protease inhibitors (Complete Mini EDTA-free, Roche; 1 tablet per 10 mL solution). Root tissues (3 g) were ground to a fine powder under liquid nitrogen using a mortar and pestle. The powdered tissue was mixed with 6 mL of extraction buffer, and the lysate was clarified by centrifugation at 14,000×g for 10 minutes.

The supernatants were incubated with 50 µL of anti-GFP magnetic beads (Miltenyi Biotec, Cat No. 130-091-125) on an end-to-end rotator at 4°C for 2 hours. Bead-bound lysates were passed through micro-columns (Miltenyi Biotec, Cat No. 130-042-701) mounted on the µMACS™ Separator. The beads were washed three times with chilled IP wash buffer (EB without digitonin), and the immunocomplexes were eluted using 50 µL of pre-heated (95°C) 1× SDS-loading buffer. Eluted proteins were resolved on 4-15% gradient SDS-PAGE gels and analyzed by immunoblotting with an anti-GFP antibody (Abcam: ab6556) to confirm SCE1 enrichment.

The immunoprecipitated eluates were further resolved on a 12% SDS-PAGE gel, electrophoresed for 20 minutes to allow protein entry into the resolving gel, and stained with colloidal Coomassie brilliant blue. Destaining was performed with a solution containing 50% water, 45% methanol, and 5% glacial acetic acid for 30 minutes. Protein bands were excised into ∼1 mm³ pieces using a sterile blade, transferred to 1.5 mL tubes, and washed three times with a solution of 50% acetonitrile (ACN) and 50 mM ammonium bicarbonate (ABC) to remove the stain. The gel pieces were dehydrated in 100% ACN for 10 minutes. Disulfide bonds were reduced with 10 mM dithiothreitol (DTT) in 50 mM ABC at 37°C for 1 hour, followed by alkylation with 55 mM iodoacetamide in 50 mM ABC at room temperature in the dark for 45 minutes. The gel pieces were again dehydrated with 100% ACN for 10 minutes.

After removing residual ACN, the gel pieces underwent in-gel digestion with trypsin/Lys-C mix (Promega, V5071) at a protease-to-protein ratio of 1:25 (w/w) at 37°C for 16 hours. Peptides were extracted in 0.1% formic acid (FA), dried in a speed-vac, and desalted using Pierce™ Peptide Desalting Spin Columns (Thermo Fisher Scientific, Cat No. 89851) following the manufacturer’s protocol. Desalted peptides were dissolved in 0.1% FA for analysis using a Q Exactive Orbitrap Mass Spectrometer.

### Bulk RNA seq analysis and statistics

For bulk seq RNA analysis, the fastq files were quality control checked by FastQC (https://www.bioinformatics.babraham.ac.uk/projects/fastqc/) and the adaptors and low-quality reads were removed by Trimmomatic.^41^ High quality reads were then mapped to Araport11,^42^ the duplicated reads were discarded by Picard (https://broadinstitute.github.io/picard/) and the read abundant were quantified by HTSeq.^43^ Differentially expressed genes (DEGs) were analysed by Deseq2.^44^ The transcripts that expressed at p_value < 0.05, fold-change ≥ 1 and ≤ –1 were considered as significantly up– and down–regulated DEGs.

### Single cell sequencing analysis

scRNA-seq datasets of wild-type Arabidopsis root samples under control conditions were used from following studies: (GSE123013),^45^ (GSE123818)^46^ and (GSE152766).^47^ Total 15 samples (except sc_9 and sc_10 from GSE152766) were chosen for further analysis. Individual biological samples from Denyer et al. 2019 and Shahan et al. 2022 were integrated into respective seurat objects (https://doi.org/10.1016/j.cell.2021.04.048).^48^ Scaled values from respective Seurat objects for each SUMO gene were used for comparisons.

### SUMO Cell atlas

Fiji software was used for image analysis. As three different settings were used to capture the extent of the variety of fluorescence signal, we divided the relative level of fluorescence in 6 different bins (Table S4).

### Stress Atlas and Tissue specific analysis

Fiji software was used for the tissue specific data analysis. A box was drawn around the four tissue types measured: epidermis, cortex, endodermis and stele. Raw fluorescence was measured and divided by the area of measurement to account for differences in area size. Fluorescence in two zones was measured, meristem and elongation zone. The meristem zone was defined by the point where the cortex cell becomes twice the size of width in length. The elongation zone is measured from the end of meristem to the start of root hair bulging. The measured fluorescence under stress vs control conditions was compared for significance using a paired t-test with bonferroni method to adjust the p-values. For each tissue type, the number of sumo system genes that show significantly different levels in their transcript or protein abundance were counted based on their p-value and the log2 fold change under stress vs control i.e. upregulated when log2 fold change > 0 and p < 0.05, downregulated when log2 fold change < 0 and p < 0.05 and unchanged for p ≥ 0.05.

### LC-MS/MS Analysis

The resulting MS/MS data were processed using Proteome Discoverer 3.1 software to identify potential interactors of SCE1. Venn diagrams were generated using the Venny 2.1 online tool (https://bioinfogp.cnb.csic.es/tools/venny/). Protein-protein interaction networks were constructed using the STRING database (https://string-db.org) and Gene Ontology (GO) analysis of stress-specific SCE1 interactors was performed using the ShinyGo 0.81 online tool (http://bioinformatics.sdstate.edu/go/). Arabidopsis thaliana root single-cell,^49^ was used to extract the expression values of SCE interactors identified from Mannitol, Salt, and Flagellin treatments across different cell clusters. The cell cluster (cell lineages or cell types) and their pseudo-time trajectory nomenclature were followed as specified in Plant sc-Atlas (https://bioit3.irc.ugent.be/plant-sc-atlas/) and each expression value being the average expression value of that gene within the specified cell cluster. The data matrix for each treatment was organized with rows representing individual genes and columns representing cell clusters (see cell cluster names on the figure). The dataset was standardised across rows to centre the data and ensure uniform scaling, thereby enabling comparison between different cell clusters (see scale bar on the figure). MATLAB’s built-in clustergram function was employed to perform hierarchical clustering on both rows and columns, using default distance metric and linkage method. The resulting clustergram was visualised as a heatmap with dendrograms to depict the hierarchical relationships.

**Table S1:**
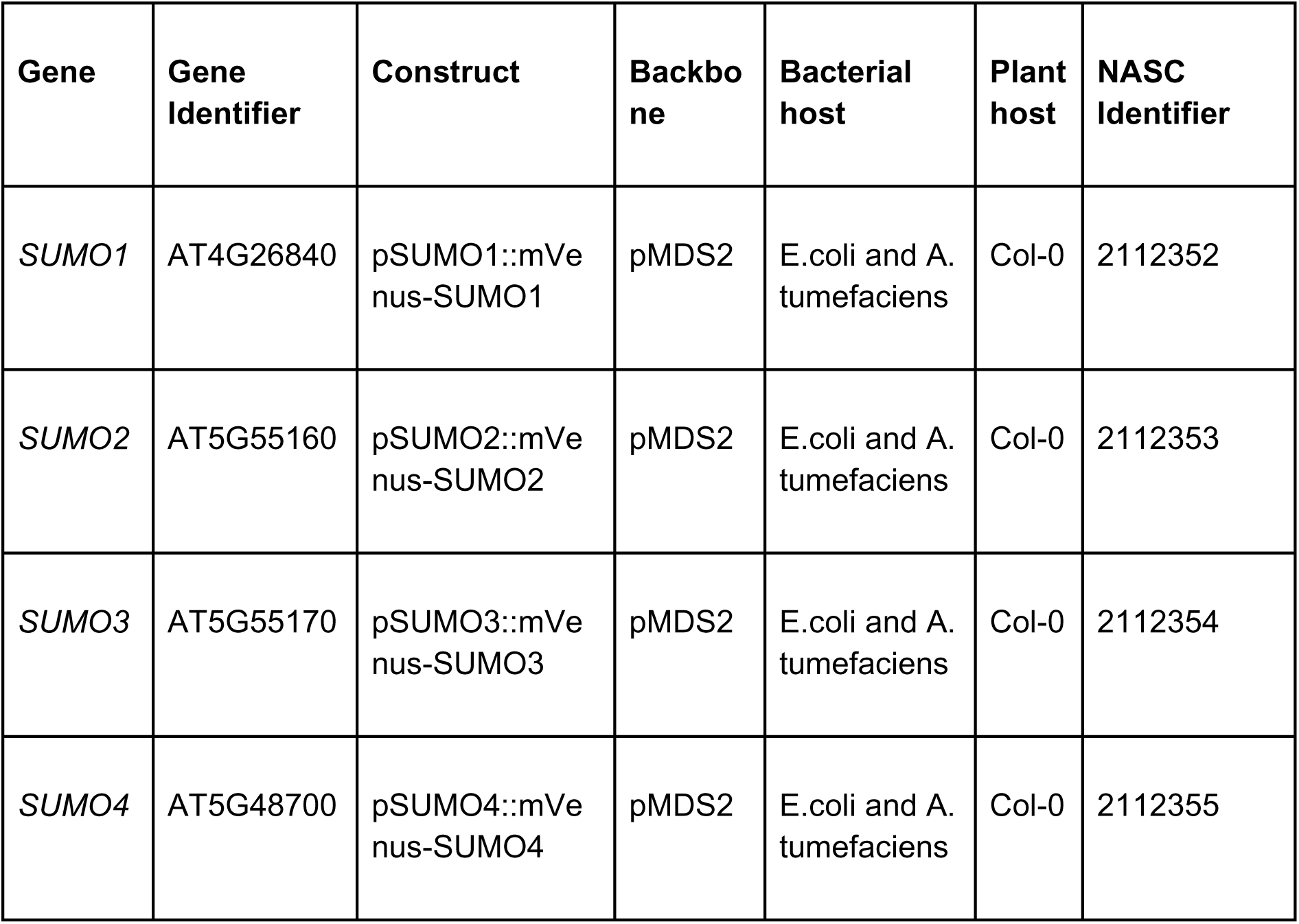
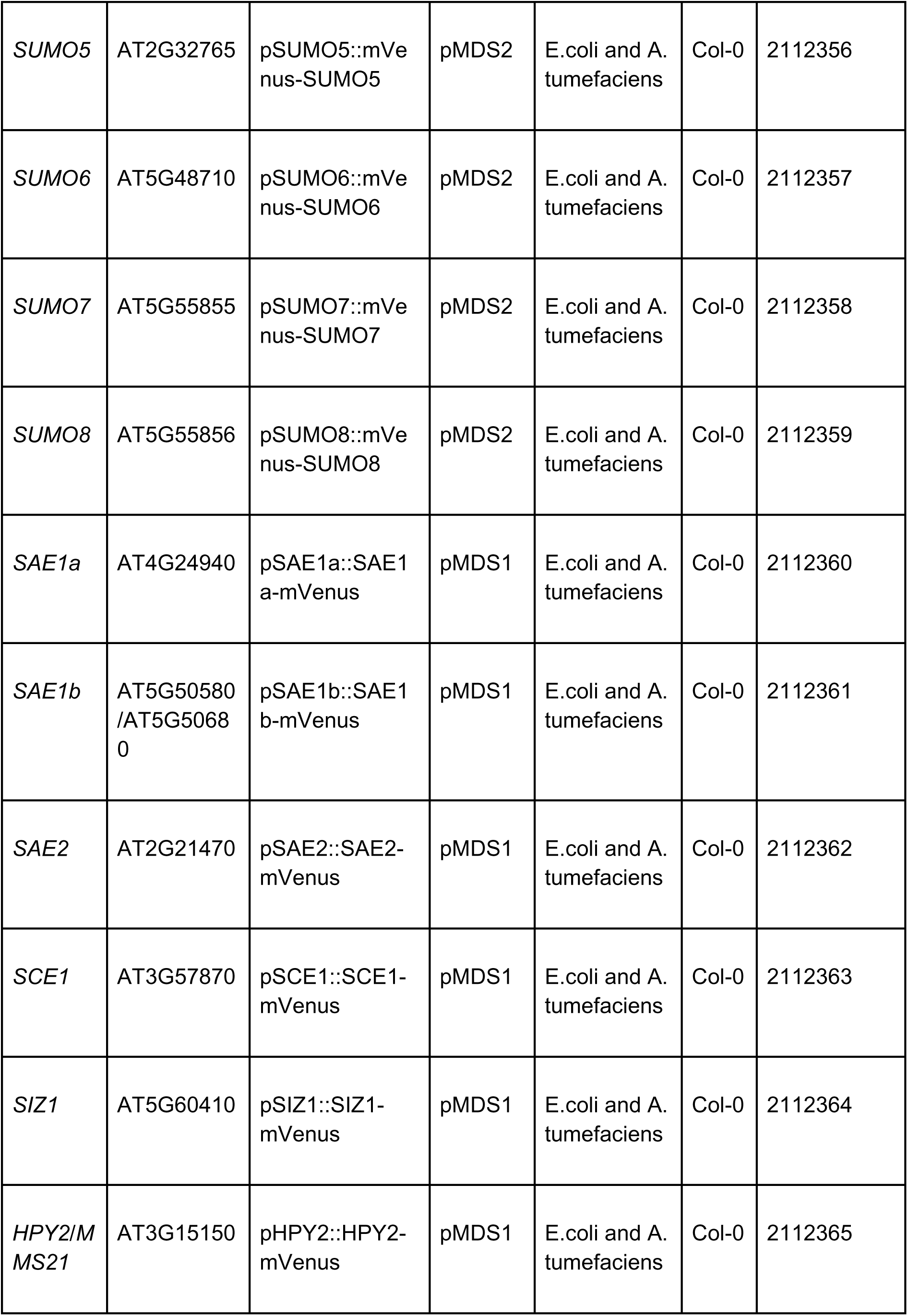
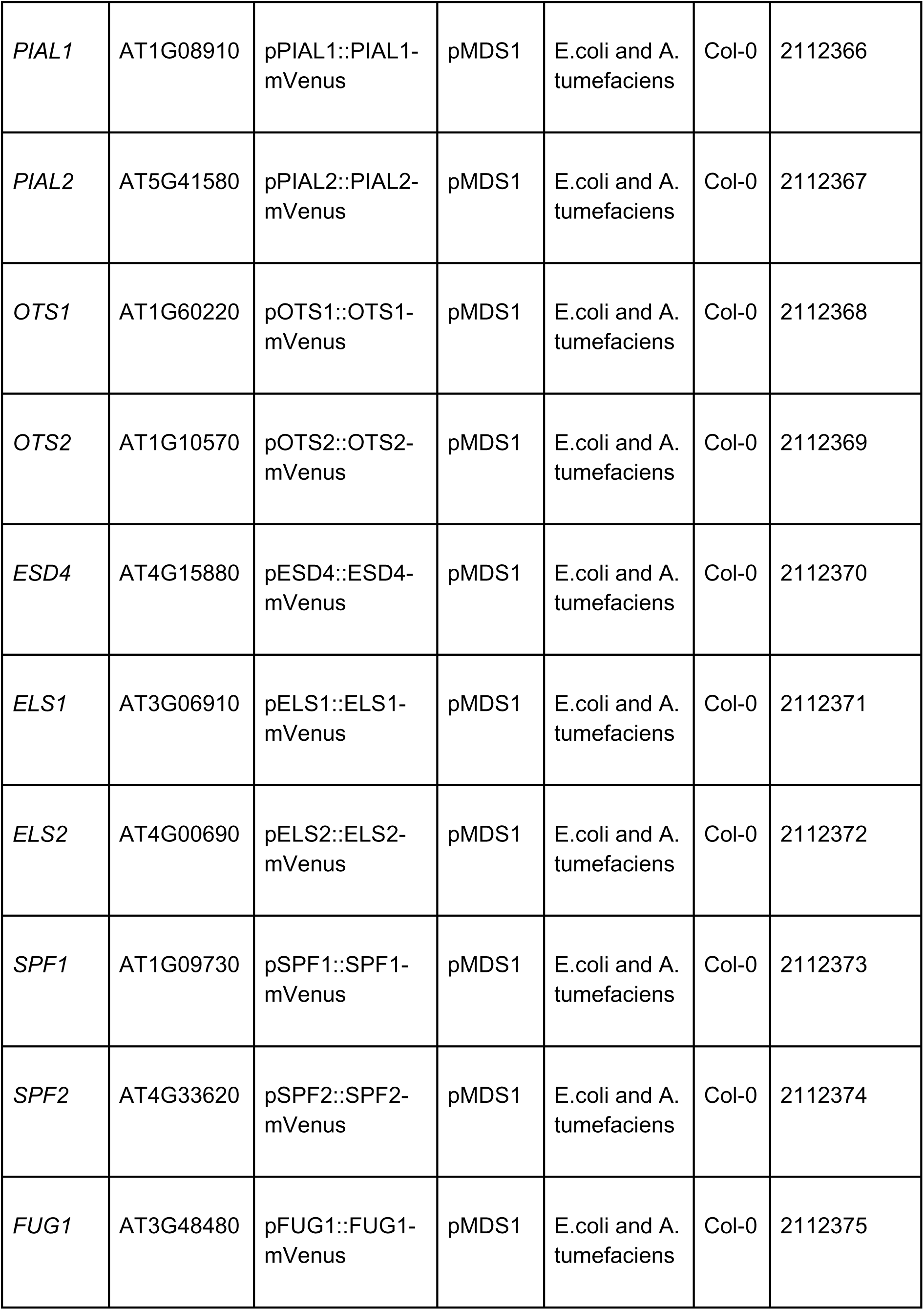
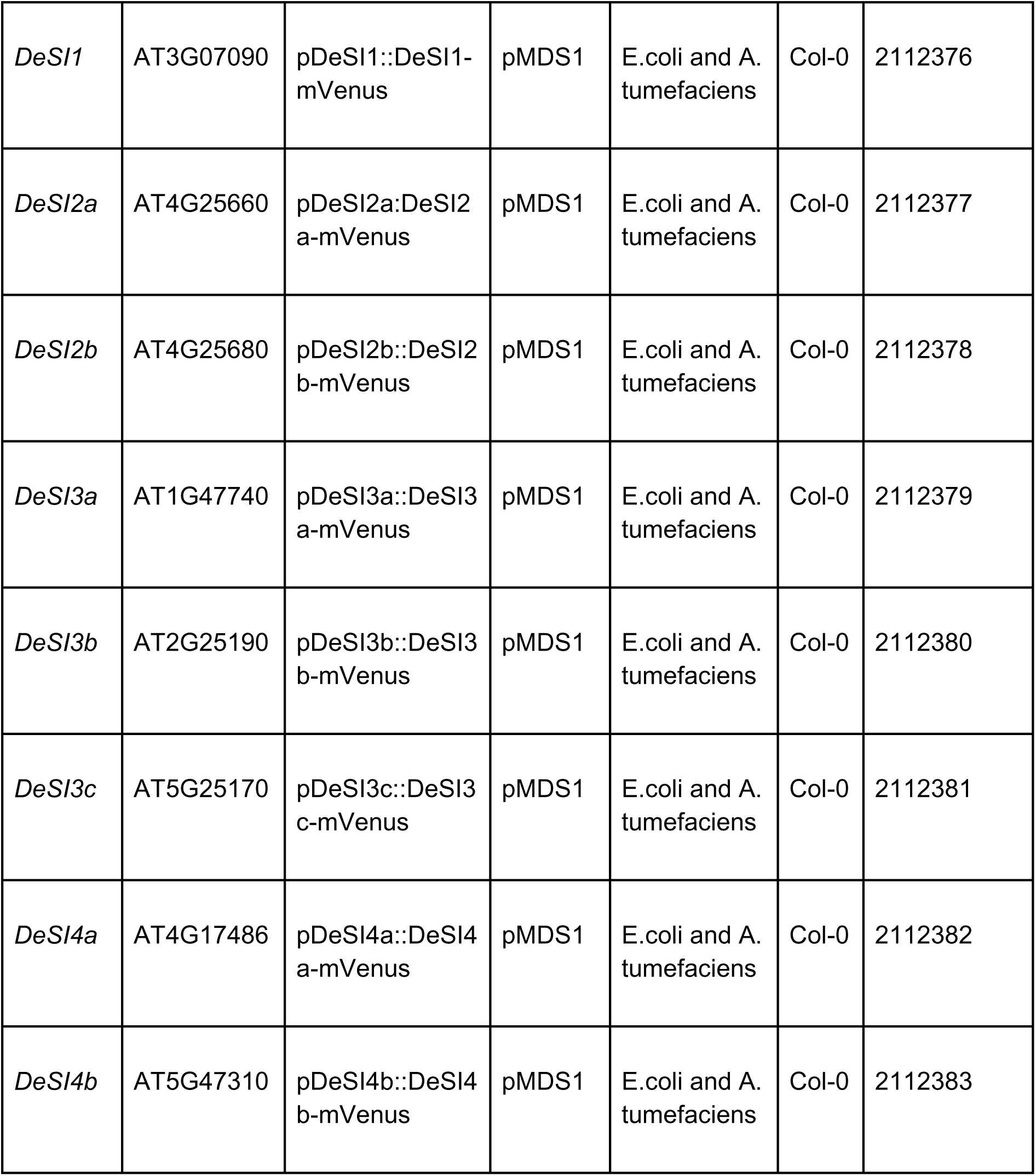
List of all constructs and plant lines made in this paper with NASC code.

**Table S2:**
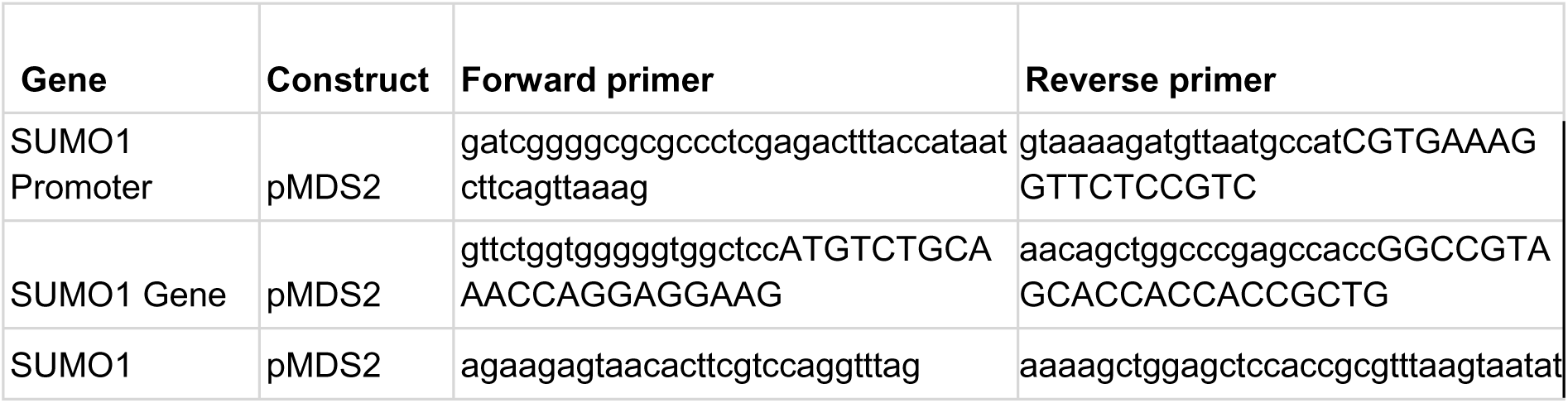
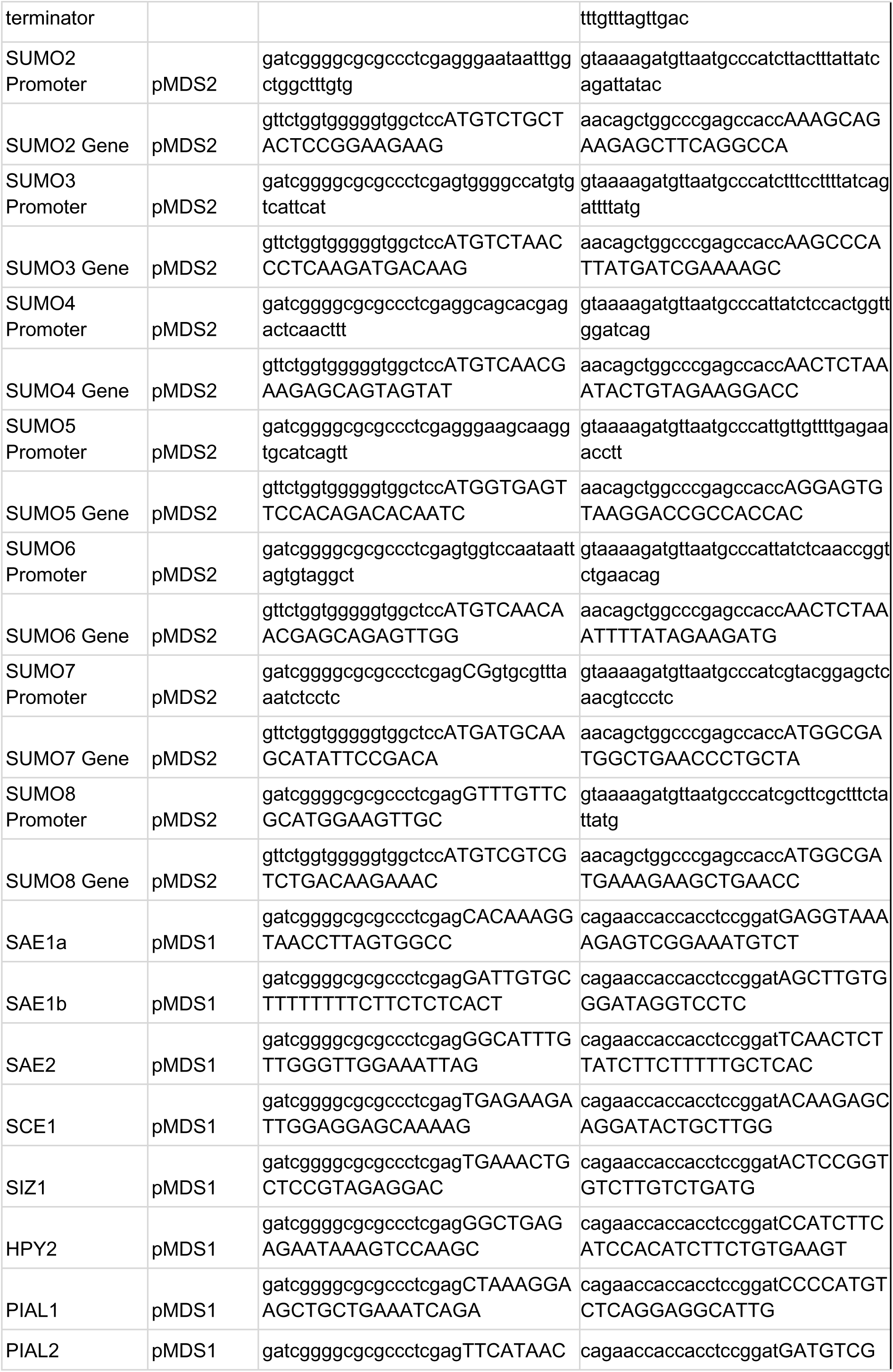
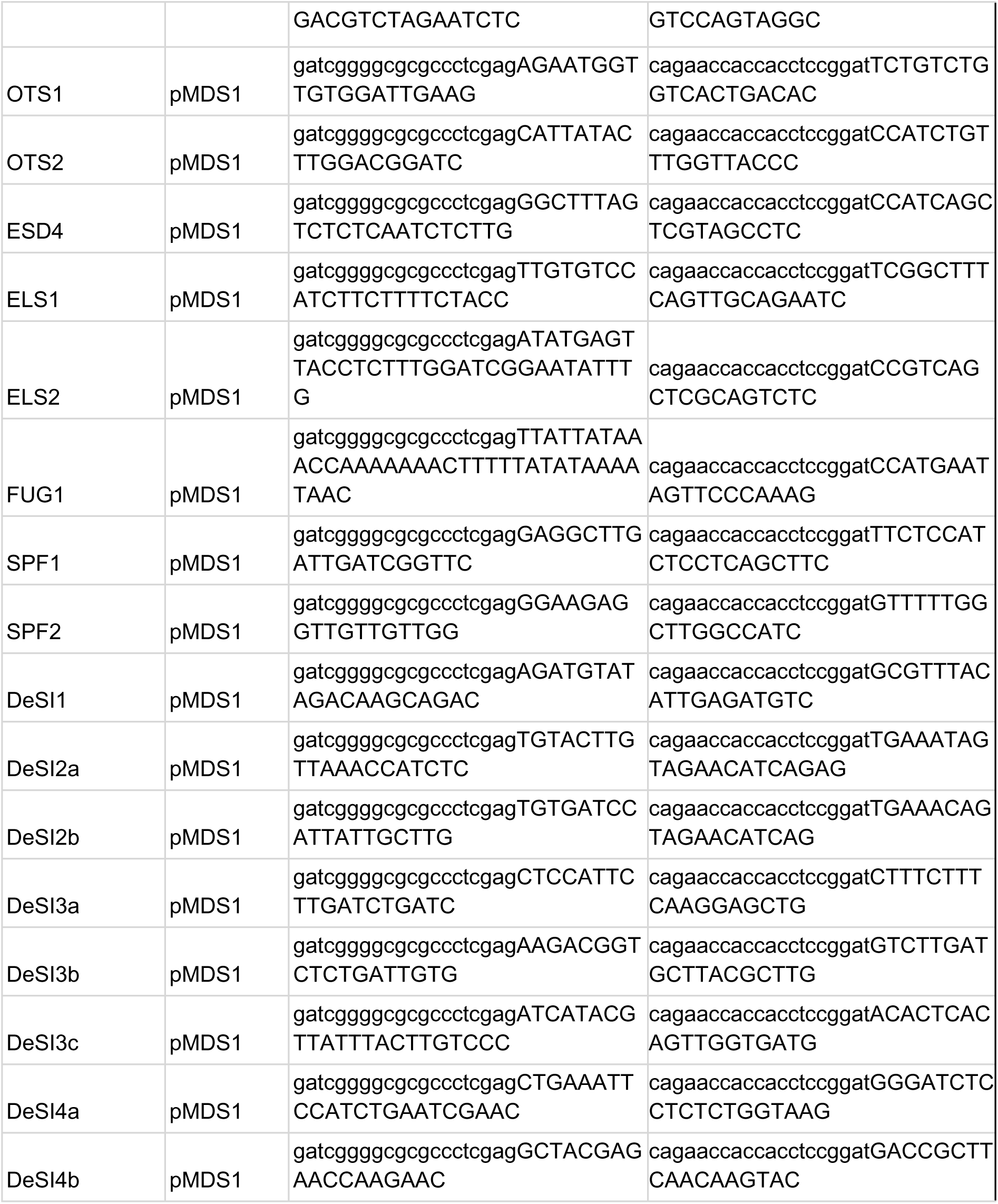
List of cloning primers for pMDS1 and pMDS2.

**Table S3:**
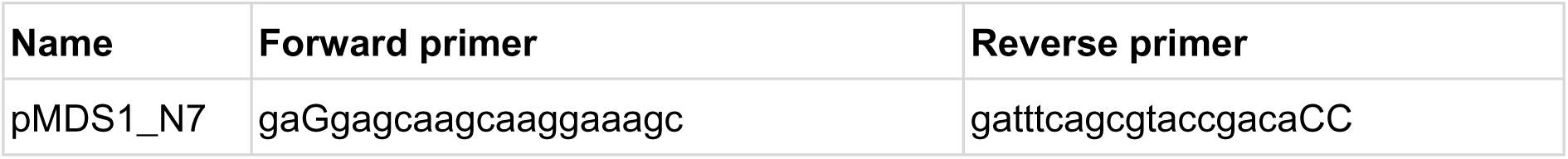
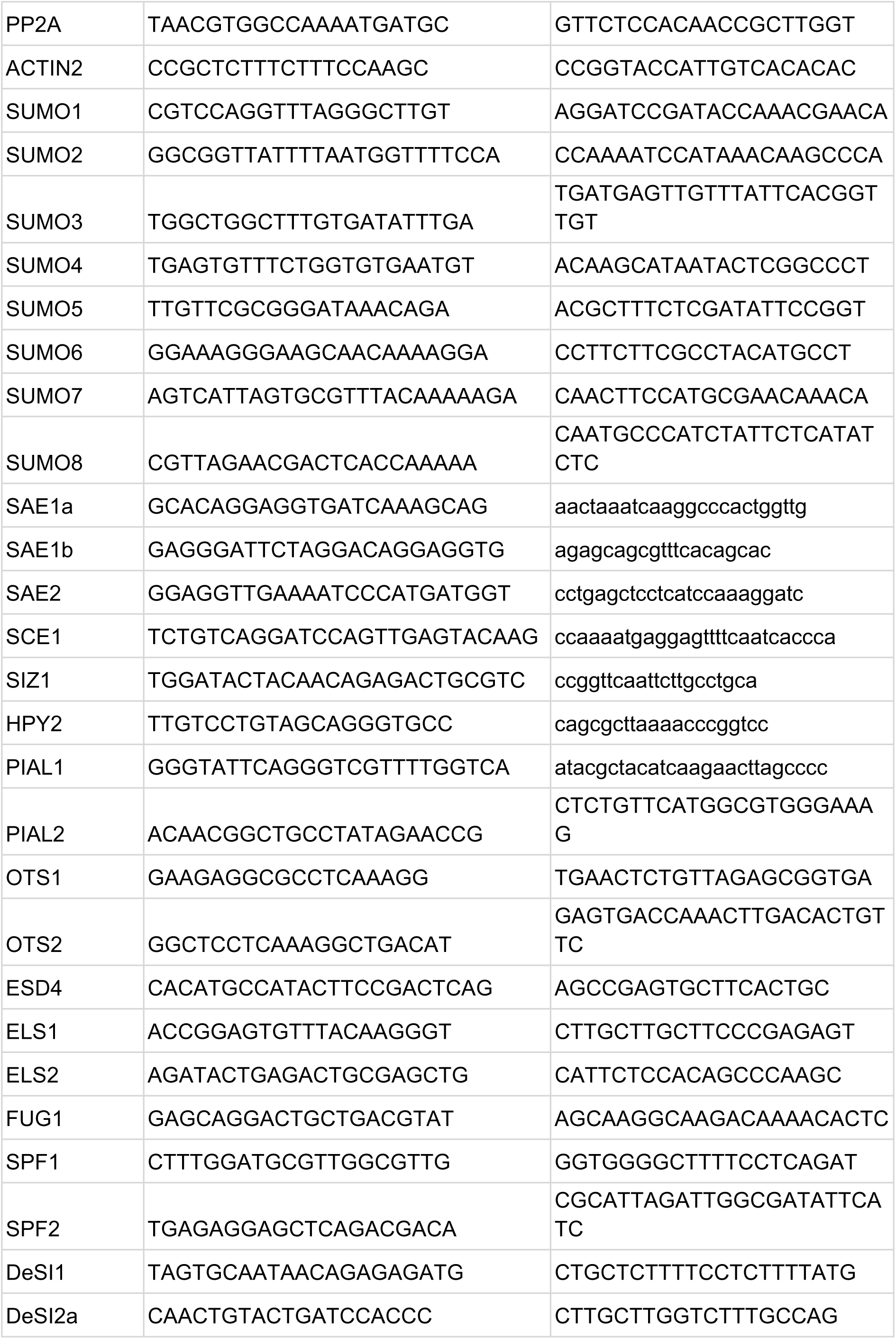
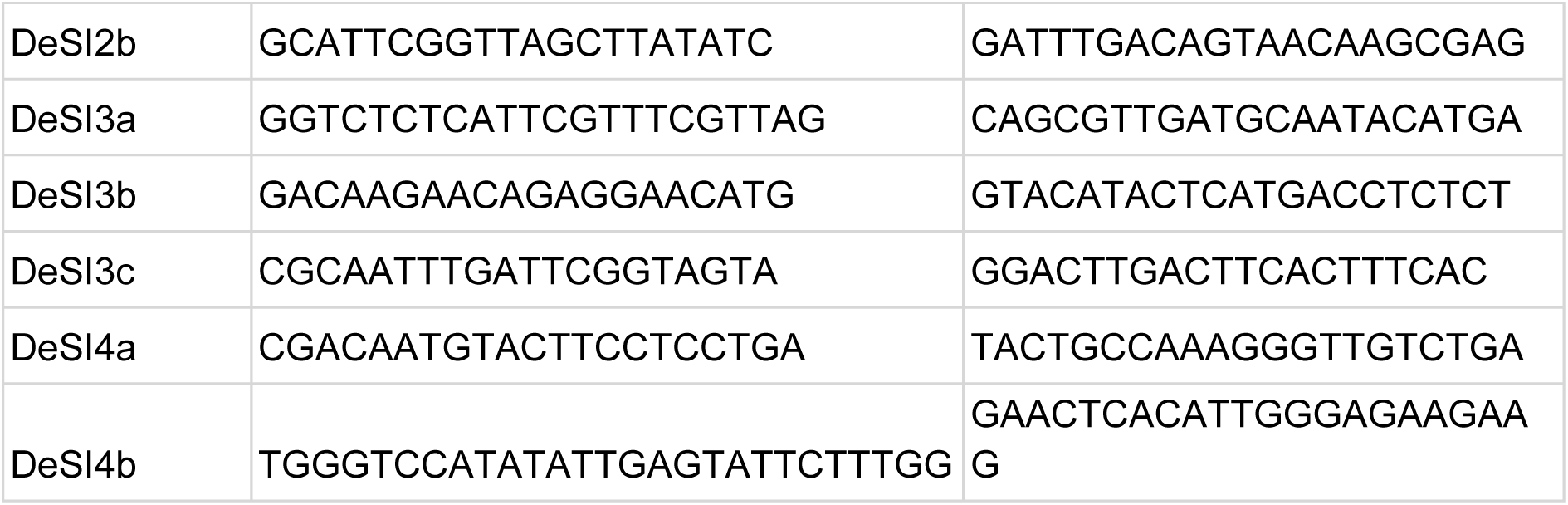
Primer details used for Real Time quantitative PCR.

**Table S4:**
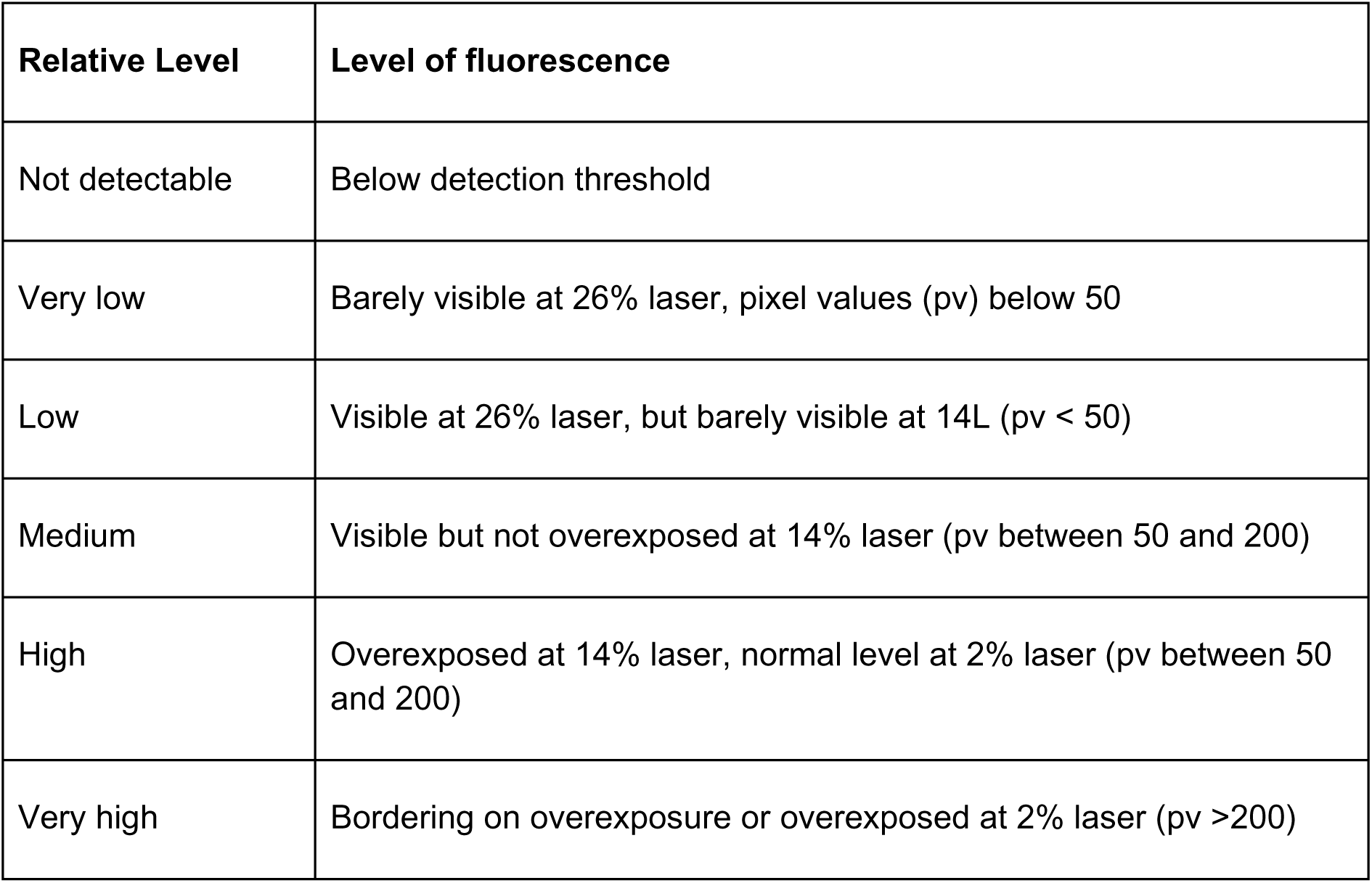
Relative fluorescence was scored in 6 different bins from not detectable to very high level of fluorescence.

---

**Table S5:** Salt SCE1 interactors.

| Protein Name | Symbol | Ensembl Gene ID | Species | Description |
| --- | --- | --- | --- | --- |
| FMOGS-OX5 | FMO GS-OX5 | AT1G12140 | Arabidopsis thaliana | flavin-monooxygenase glucosinolate S-oxygenase 5 [Source:NCBI gene (formerly Entrezgene);Acc:837766] |
| HSP70-5 | Hsp70b | AT1G16030 | Arabidopsis thaliana | heat shock protein 70B [Source:NCBI gene (formerly Entrezgene);Acc:838174] |
| BTF3 | BTF3 | AT1G17880 | Arabidopsis thaliana | basic transcription factor 3 [Source:NCBI gene (formerly Entrezgene);Acc:838367] |
| DHAR1 | DHAR1 | AT1G19570 | Arabidopsis thaliana | dehydroascorbate reductase [Source:NCBI gene (formerly Entrezgene);Acc:838544] |
| RPN3A | EMB2719 | AT1G20200 | Arabidopsis thaliana | PAM domain (PCI/PINT associated module) protein [Source:NCBI gene (formerly Entrezgene);Acc:838609] |
| RPT4B | AT1G45000 | AT1G45000 | Arabidopsis thaliana | AAA-type ATPase family protein [Source:NCBI gene (formerly Entrezgene);Acc:841065] |
| GLL22 | GLL22 | AT1G54000 | Arabidopsis thaliana | GDSL-like Lipase/Acylhydrolase superfamily protein [Source:NCBI gene (formerly Entrezgene);Acc:841838] |
| MVP1 | MVP1 | AT1G54030 | Arabidopsis thaliana | GDSL-like Lipase/Acylhydrolase superfamily protein [Source:NCBI gene (formerly Entrezgene);Acc:841841] |
| NOP56 | NOP56 | AT1G56110 | Arabidopsis thaliana | NOP56-like pre RNA processing ribonucleoprotein [Source:NCBI gene (formerly Entrezgene);Acc:842063] |
| ABCG36 | PEN3 | AT1G59870 | Arabidopsis thaliana | ABC-2 and Plant PDR ABC-type transporter family protein [Source:NCBI gene (formerly Entrezgene);Acc:842281] |
| RPL34B | RPL34 | AT1G69620 | Arabidopsis thaliana | ribosomal protein L34 [Source:NCBI gene (formerly Entrezgene);Acc:843298] |
| P40 | P40 | AT1G72370 | Arabidopsis thaliana | 40s ribosomal protein SA [Source:NCBI gene (formerly Entrezgene);Acc:843569] |
| RHM1 | RHM1 | AT1G78570 | Arabidopsis thaliana | rhamnose biosynthesis 1 [Source:NCBI gene (formerly Entrezgene);Acc:844193] |
| SKD1 | SKD1 | AT2G27600 | Arabidopsis thaliana | AAA-type ATPase family protein [Source:NCBI gene (formerly Entrezgene);Acc:817306] |
| RPN2A | AT2G32730 | AT2G32730 | Arabidopsis thaliana | 26S proteasome regulatory complex, non-ATPase subcomplex, Rpn2/Psmd1 subunit [Source:NCBI gene (formerly Entrezgene);Acc:817833] |
| RPL12A | AT2G37190 | AT2G37190 | Arabidopsis thaliana | Ribosomal protein L11 family protein [Source:NCBI gene (formerly Entrezgene);Acc:818295] |
| ANN4 | ANNAT4 | AT2G38750 | Arabidopsis thaliana | annexin 4 [Source:NCBI gene (formerly Entrezgene);Acc:818457] |
| RPS2C | AT2G41840 | AT2G41840 | Arabidopsis thaliana | Ribosomal protein S5 family protein [Source:NCBI gene (formerly Entrezgene);Acc:818784] |
| RH37 | AT2G42520 | AT2G42520 | Arabidopsis thaliana | P-loop containing nucleoside triphosphate hydrolases superfamily protein [Source:NCBI gene (formerly Entrezgene);Acc:818852] |
| CSY4 | ATCS | AT2G44350 | Arabidopsis thaliana | Citrate synthase family protein [Source:NCBI gene (formerly Entrezgene);Acc:819042] |
| CCT6A | AT3G02530 | AT3G02530 | Arabidopsis thaliana | TCP-1/cpn60 chaperonin family protein [Source:NCBI gene (formerly Entrezgene);Acc:821160] |
| RPT5A | RPT5A | AT3G05530 | Arabidopsis thaliana | regulatory particle triple-A ATPase 5A [Source:NCBI gene (formerly Entrezgene);Acc:819718] |
| CDC48A | CDC48 | AT3G09840 | Arabidopsis thaliana | cell division cycle 48 [Source:NCBI gene (formerly Entrezgene);Acc:820142] |
| CPN60B2 | Cpn60beta2 | AT3G13470 | Arabidopsis thaliana | TCP-1/cpn60 chaperonin family protein [Source:NCBI gene (formerly Entrezgene);Acc:820549] |
| RPL7D | AT3G13580 | AT3G13580 | Arabidopsis thaliana | Ribosomal protein L30/L7 family protein [Source:NCBI gene (formerly Entrezgene);Acc:820560] |
| PPC3 | PPC3 | AT3G14940 | Arabidopsis thaliana | phosphoenolpyruvate carboxylase 3 [Source:NCBI gene (formerly Entrezgene);Acc:820723] |
| JAL35 | JR1 | AT3G16470 | Arabidopsis thaliana | Mannose-binding lectin superfamily protein [Source:NCBI gene (formerly Entrezgene);Acc:820895] |
| CCT4 | AT3G18190 | AT3G18190 | Arabidopsis thaliana | TCP-1/cpn60 chaperonin family protein [Source:NCBI gene (formerly Entrezgene);Acc:821346] |
| EIF4A3 | EIF4A-III | AT3G19760 | Arabidopsis thaliana | eukaryotic initiation factor 4A-III [Source:NCBI gene (formerly Entrezgene);Acc:821513] |
| CCT1 | TCP-1 | AT3G20050 | Arabidopsis thaliana | T-complex protein 1 alpha subunit [Source:NCBI gene (formerly Entrezgene);Acc:821544] |
| RPL27B | AT3G22230 | AT3G22230 | Arabidopsis thaliana | Ribosomal L27e protein family [Source:NCBI gene (formerly Entrezgene);Acc:821787] |
| PHB4 | PHB4 | AT3G27280 | Arabidopsis thaliana | prohibitin 4 [Source:NCBI gene (formerly Entrezgene);Acc:822347] |
| SDR4 | SDR4 | AT3G29250 | Arabidopsis thaliana | NAD(P)-binding Rossmann-fold superfamily protein [Source:NCBI gene (formerly Entrezgene);Acc:822580] |
| DAYSLEEPER | DAYSLEEPER | AT3G42170 | Arabidopsis thaliana | BED zinc finger and hAT dimerization domain-containing protein DAYSLEEPER [Source:NCBI gene (formerly Entrezgene);Acc:3769417] |
| BBC1 | BBC1 | AT3G49010 | Arabidopsis thaliana | breast basic conserved 1 [Source:NCBI gene (formerly Entrezgene);Acc:824062] |
| SR34A | SR34a | AT3G49430 | Arabidopsis thaliana | SER/ARG-rich protein 34A [Source:NCBI gene (formerly Entrezgene);Acc:824105] |
| RH11 | AT3G58510 | AT3G58510 | Arabidopsis thaliana | DEA(D/H)-box RNA helicase family protein [Source:NCBI gene (formerly Entrezgene);Acc:825020] |
| CPK21 | CPK21 | AT4G04720 | Arabidopsis thaliana | calcium-dependent protein kinase 21 [Source:NCBI gene (formerly Entrezgene);Acc:825807] |
| VHA-E1 | TUF | AT4G11150 | Arabidopsis thaliana | vacuolar ATP synthase subunit E1 [Source:NCBI gene (formerly Entrezgene);Acc:826716] |
| RABA1D | RABA1d | AT4G18800 | Arabidopsis thaliana | RAB GTPase homolog A1D [Source:NCBI gene (formerly Entrezgene);Acc:827614] |
| RPN1B | RPN1B | AT4G28470 | Arabidopsis thaliana | 26S proteasome regulatory subunit S2 1B [Source:NCBI gene (formerly Entrezgene);Acc:828964] |
| RPS6 | RPS6 | AT4G31700 | Arabidopsis thaliana | ribosomal protein S6 [Source:NCBI gene (formerly Entrezgene);Acc:829298] |
| VAB2 | VAB2 | AT4G38510 | Arabidopsis thaliana | ATPase, V1 complex, subunit B protein [Source:NCBI gene (formerly Entrezgene);Acc:830008] |
| CDC48E | AtCDC48C | AT5G03340 | Arabidopsis thaliana | ATPase, AAA-type, CDC48 protein [Source:NCBI gene (formerly Entrezgene);Acc:831870] |
| UAP56B | UAP56b | AT5G11200 | Arabidopsis thaliana | DEAD/DEAH box RNA helicase family protein [Source:NCBI gene (formerly Entrezgene);Acc:830990] |
| ASP3 | ASP3 | AT5G11520 | Arabidopsis thaliana | aspartate aminotransferase 3 [Source:NCBI gene (formerly Entrezgene);Acc:831024] |
| TUBB6 | TUB6 | AT5G12250 | Arabidopsis thaliana | beta-6 tubulin [Source:NCBI gene (formerly Entrezgene);Acc:831100] |
| ARFA1B | ARFA1B | AT5G14670 | Arabidopsis thaliana | ADP-ribosylation factor A1B [Source:NCBI gene (formerly Entrezgene);Acc:831319] |
| RPS9B | AT5G15200 | AT5G15200 | Arabidopsis thaliana | Ribosomal protein S4 [Source:NCBI gene (formerly Entrezgene);Acc:831372] |
| RPS8A | AT5G20290 | AT5G20290 | Arabidopsis thaliana | Ribosomal protein S8e family protein [Source:NCBI gene (formerly Entrezgene);Acc:832151] |
| MIRO1 | MIRO1 | AT5G27540 | Arabidopsis thaliana | MIRO-related GTP-ase 1 [Source:NCBI gene (formerly Entrezgene);Acc:832814] |
| RPL18C | AT5G27850 | AT5G27850 | Arabidopsis thaliana | Ribosomal protein L18e/L15 superfamily protein [Source:NCBI gene (formerly Entrezgene);Acc:832848] |
| PDP3 | AT5G40340 | AT5G40340 | Arabidopsis thaliana | Tudor/PWWP/MBT superfamily protein [Source:NCBI gene (formerly Entrezgene);Acc:834032] |
| PGD3 | AT5G41670 | AT5G41670 | Arabidopsis thaliana | 6-phosphogluconate dehydrogenase family protein [Source:NCBI gene (formerly Entrezgene);Acc:834169] |
| RABA1C | RABA1c | AT5G45750 | Arabidopsis thaliana | RAB GTPase homolog A1C [Source:NCBI gene (formerly Entrezgene);Acc:834614] |
| RABA4C | RABA4C | AT5G47960 | Arabidopsis thaliana | RAB GTPase homolog A4C [Source:NCBI gene (formerly Entrezgene);Acc:834847] |
| DEA(D/H)-box RNA helicase family protein | A0A1P8ATJ6 | AT1G51380 | Arabidopsis thaliana | DEA(D/H)-box RNA helicase family protein |
| Nascent polypeptide-associated complex subunit beta | Q9CAT7 | AT1G73230 | Arabidopsis thaliana | Nascent polypeptide-associated complex subunit beta |
| Probable fructokinase-1 | Q9SID0 | AT2G31390 | Arabidopsis thaliana | Probable fructokinase-1 |
| Coatomer subunit beta' | A0A1I9LTK3 | AT3G15980 | Arabidopsis thaliana | Coatomer subunit beta' |
| 40S ribosomal protein S4 | F4K5C7 | AT5G07090 | Arabidopsis thaliana | 40S ribosomal protein S4 |
| Glutamate--tRNA ligase, cytoplasmic | O82462 | AT5G26710 | Arabidopsis thaliana | Glutamate--tRNA ligase, cytoplasmic |
| Histone H4 | P59259 | AT5G59970 | Arabidopsis thaliana | Histone H4 |
| Indole-3-acetic acid-amido synthetase GH3.17 | Q9FZ87 | At1g28130 | Arabidopsis thaliana | Indole-3-acetic acid-amido synthetase GH3.17 |
| Mov34/MPN/PAD-1 family protein | A8MR12 | At5g23540 | Arabidopsis thaliana | Mov34/MPN/PAD-1 family protein |

**Table S6:**
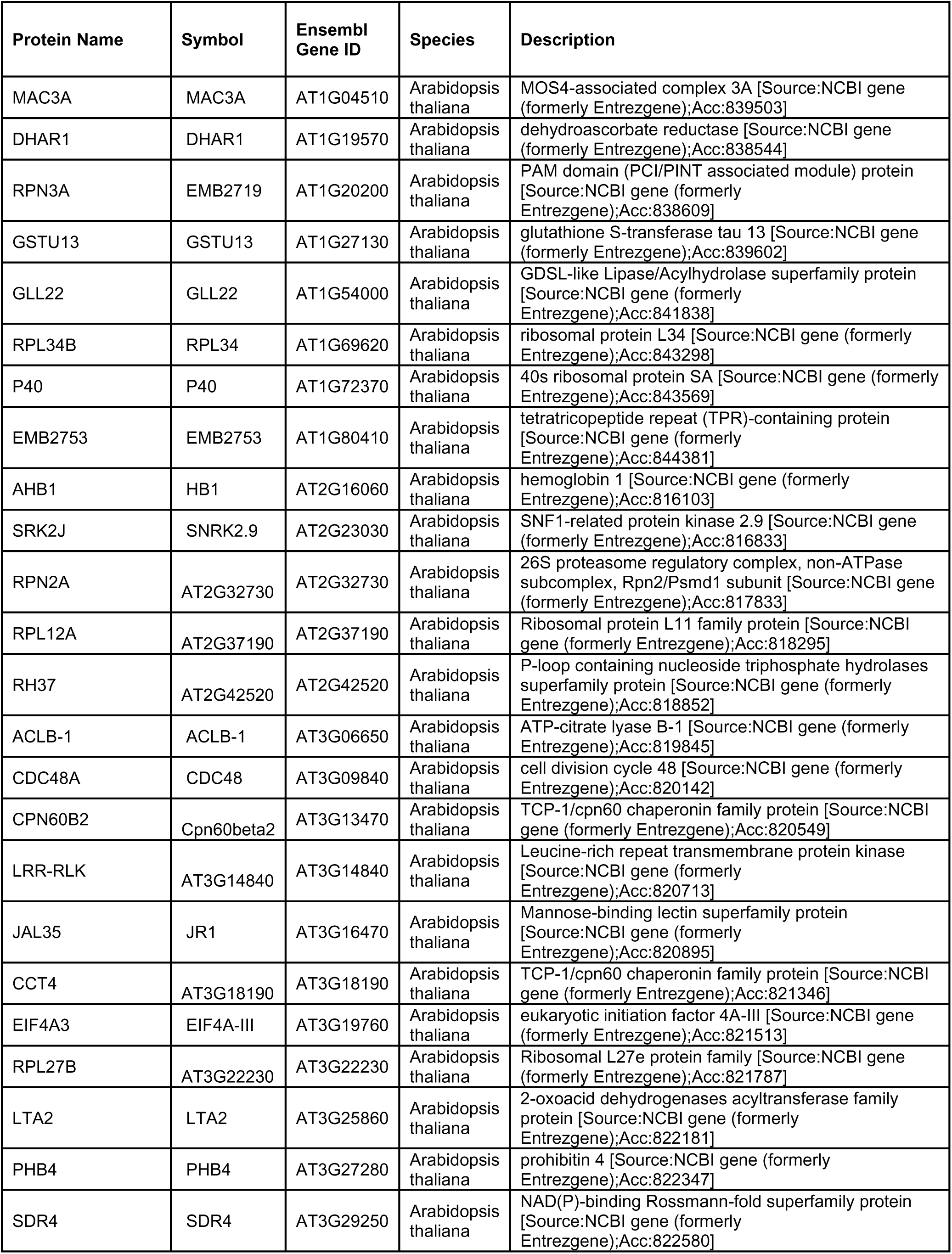
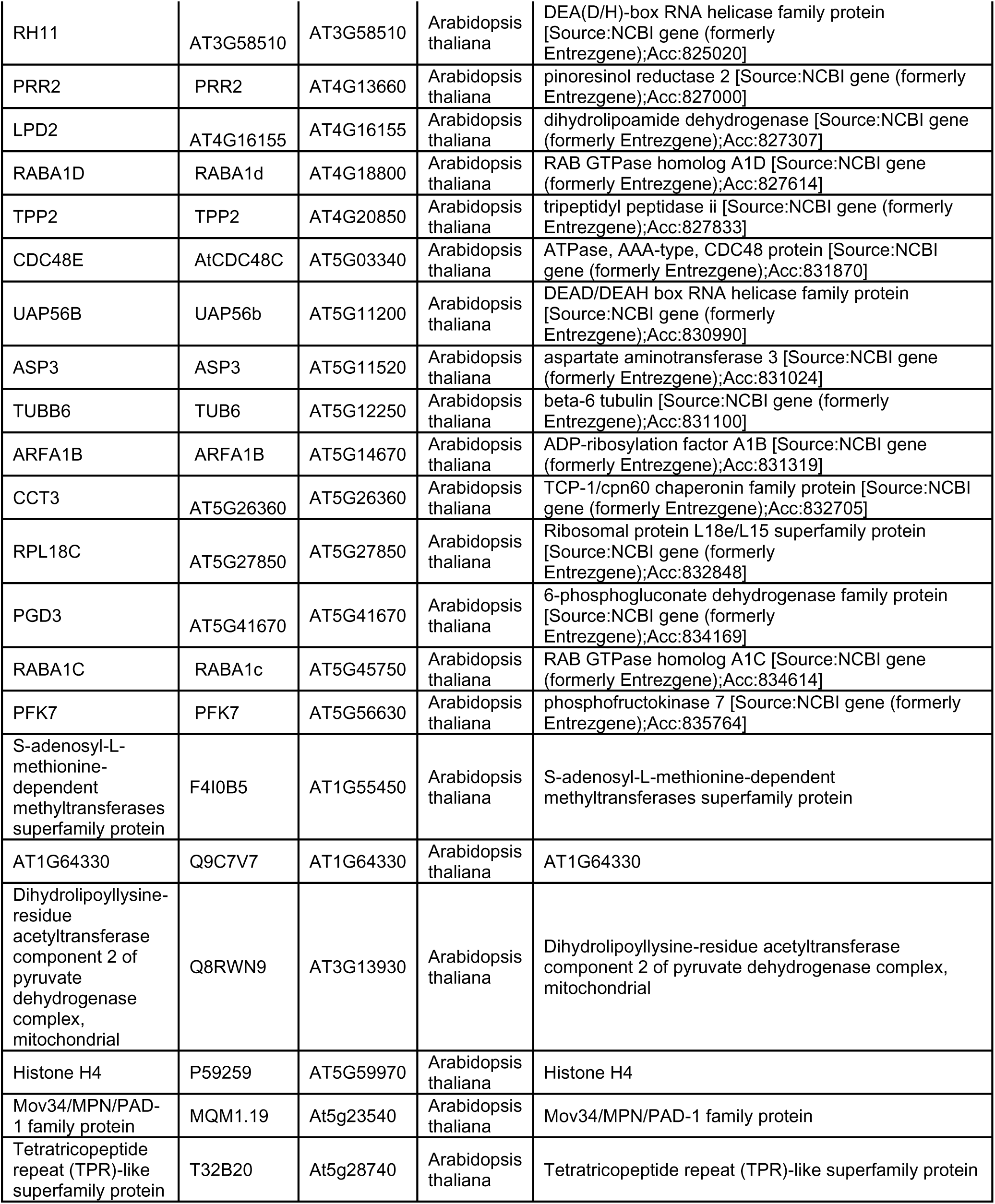
Mannitol SCE1 interactors.

**Table S7:** Flagellin22 SCE1 interactors.

| Protein Name | Symbol | Ensembl Gene ID | Species | Description |
| --- | --- | --- | --- | --- |
| UGE1 | UGE1 | AT1G12780 | Arabidopsis thaliana | UDP-D-glucose/UDP-D-galactose 4-epimerase 1 [Source:NCBI gene (formerly Entrezgene);Acc:837834] |
| RPN3A | EMB2719 | AT1G20200 | Arabidopsis thaliana | PAM domain (PCI/PINT associated module) protein [Source:NCBI gene (formerly Entrezgene);Acc:838609] |
| GSTU13 | GSTU13 | AT1G27130 | Arabidopsis thaliana | glutathione S-transferase tau 13 [Source:NCBI gene (formerly Entrezgene);Acc:839602] |
| RHM2 | MUM4 | AT1G53500 | Arabidopsis thaliana | NAD-dependent epimerase/dehydratase family protein [Source:NCBI gene (formerly Entrezgene);Acc:841785] |
| GLL22 | GLL22 | AT1G54000 | Arabidopsis thaliana | GDSL-like Lipase/Acylhydrolase superfamily protein [Source:NCBI gene (formerly Entrezgene);Acc:841838] |
| UPL1 | UPL1 | AT1G55860 | Arabidopsis thaliana | LOW protein: E3 ubiquitin ligase-like protein [Source:NCBI gene (formerly Entrezgene);Acc:842036] |
| SRK2B | SNRK2.10 | AT1G60940 | Arabidopsis thaliana | SNF1-related protein kinase 2.10 [Source:NCBI gene (formerly Entrezgene);Acc:842385] |
| RPL34B | RPL34 | AT1G69620 | Arabidopsis thaliana | ribosomal protein L34 [Source:NCBI gene (formerly Entrezgene);Acc:843298] |
| P40 | P40 | AT1G72370 | Arabidopsis thaliana | 40s ribosomal protein SA [Source:NCBI gene (formerly Entrezgene);Acc:843569] |
| RHM1 | RHM1 | AT1G78570 | Arabidopsis thaliana | rhamnose biosynthesis 1 [Source:NCBI gene (formerly Entrezgene);Acc:844193] |
| EMB2753 | EMB2753 | AT1G80410 | Arabidopsis thaliana | tetratricopeptide repeat (TPR)-containing protein [Source:NCBI gene (formerly Entrezgene);Acc:844381] |
| SOT12 | SOT12 | AT2G03760 | Arabidopsis thaliana | sulfotransferase 12 [Source:NCBI gene (formerly Entrezgene);Acc:814903] |
| AHB1 | HB1 | AT2G16060 | Arabidopsis thaliana | hemoglobin 1 [Source:NCBI gene (formerly Entrezgene);Acc:816103] |
| SRK2J | SNRK2.9 | AT2G23030 | Arabidopsis thaliana | SNF1-related protein kinase 2.9 [Source:NCBI gene (formerly Entrezgene);Acc:816833] |
| RPN2A | AT2G32730 | AT2G32730 | Arabidopsis thaliana | 26S proteasome regulatory complex, non-ATPase subcomplex, Rpn2/Psmd1 subunit [Source:NCBI gene (formerly Entrezgene);Acc:817833] |
| RPL12A | AT2G37190 | AT2G37190 | Arabidopsis thaliana | Ribosomal protein L11 family protein [Source:NCBI gene (formerly Entrezgene);Acc:818295] |
| ANN4 | ANNAT4 | AT2G38750 | Arabidopsis thaliana | annexin 4 [Source:NCBI gene (formerly Entrezgene);Acc:818457] |
| RPS2C | AT2G41840 | AT2G41840 | Arabidopsis thaliana | Ribosomal protein S5 family protein [Source:NCBI gene (formerly Entrezgene);Acc:818784] |
| RH37 | AT2G42520 | AT2G42520 | Arabidopsis thaliana | P-loop containing nucleoside triphosphate hydrolases superfamily protein [Source:NCBI gene (formerly Entrezgene);Acc:818852] |
| CSY4 | ATCS | AT2G44350 | Arabidopsis thaliana | Citrate synthase family protein [Source:NCBI gene (formerly Entrezgene);Acc:819042] |
| CCT6A | AT3G02530 | AT3G02530 | Arabidopsis thaliana | TCP-1/cpn60 chaperonin family protein [Source:NCBI gene (formerly Entrezgene);Acc:821160] |
| ACLB-1 | ACLB-1 | AT3G06650 | Arabidopsis thaliana | ATP-citrate lyase B-1 [Source:NCBI gene (formerly Entrezgene);Acc:819845] |
| CDC48A | CDC48 | AT3G09840 | Arabidopsis thaliana | cell division cycle 48 [Source:NCBI gene (formerly Entrezgene);Acc:820142] |
| CPN60B2 | Cpn60beta2 | AT3G13470 | Arabidopsis thaliana | TCP-1/cpn60 chaperonin family protein [Source:NCBI gene (formerly Entrezgene);Acc:820549] |
| RPL7D | AT3G13580 | AT3G13580 | Arabidopsis thaliana | Ribosomal protein L30/L7 family protein [Source:NCBI gene (formerly Entrezgene);Acc:820560] |
| LRR-RLK | AT3G14840 | AT3G14840 | Arabidopsis thaliana | Leucine-rich repeat transmembrane protein kinase [Source:NCBI gene (formerly Entrezgene);Acc:820713] |
| JAL35 | JR1 | AT3G16470 | Arabidopsis thaliana | Mannose-binding lectin superfamily protein [Source:NCBI gene (formerly Entrezgene);Acc:820895] |
| CCT4 | AT3G18190 | AT3G18190 | Arabidopsis thaliana | TCP-1/cpn60 chaperonin family protein [Source:NCBI gene (formerly Entrezgene);Acc:821346] |
| EIF4A3 | EIF4A-III | AT3G19760 | Arabidopsis thaliana | eukaryotic initiation factor 4A-III [Source:NCBI gene (formerly Entrezgene);Acc:821513] |
| CCT1 | TCP-1 | AT3G20050 | Arabidopsis thaliana | T-complex protein 1 alpha subunit [Source:NCBI gene (formerly Entrezgene);Acc:821544] |
| RPL27B | AT3G22230 | AT3G22230 | Arabidopsis thaliana | Ribosomal L27e protein family [Source:NCBI gene (formerly Entrezgene);Acc:821787] |
| LTA2 | LTA2 | AT3G25860 | Arabidopsis thaliana | 2-oxoacid dehydrogenases acyltransferase family protein [Source:NCBI gene (formerly Entrezgene);Acc:822181] |
| PHB4 | PHB4 | AT3G27280 | Arabidopsis thaliana | prohibitin 4 [Source:NCBI gene (formerly Entrezgene);Acc:822347] |
| SDR4 | SDR4 | AT3G29250 | Arabidopsis thaliana | NAD(P)-binding Rossmann-fold superfamily protein [Source:NCBI gene (formerly Entrezgene);Acc:822580] |
| DAYSLEEPER | DAYSLEEPER | AT3G42170 | Arabidopsis thaliana | BED zinc finger and hAT dimerization domain-containing protein DAYSLEEPER [Source:NCBI gene (formerly Entrezgene);Acc:3769417] |
| BBC1 | BBC1 | AT3G49010 | Arabidopsis thaliana | breast basic conserved 1 [Source:NCBI gene (formerly Entrezgene);Acc:824062] |
| SR34A | SR34a | AT3G49430 | Arabidopsis thaliana | SER/ARG-rich protein 34A [Source:NCBI gene (formerly Entrezgene);Acc:824105] |
| RH11 | AT3G58510 | AT3G58510 | Arabidopsis thaliana | DEA(D/H)-box RNA helicase family protein [Source:NCBI gene (formerly Entrezgene);Acc:825020] |
| CPK21 | CPK21 | AT4G04720 | Arabidopsis thaliana | calcium-dependent protein kinase 21 [Source:NCBI gene (formerly Entrezgene);Acc:825807] |
| VHA-E1 | TUF | AT4G11150 | Arabidopsis thaliana | vacuolar ATP synthase subunit E1 [Source:NCBI gene (formerly Entrezgene);Acc:826716] |
| PRR2 | PRR2 | AT4G13660 | Arabidopsis thaliana | pinorexinol reductase 2 [Source:NCBI gene (formerly Entrezgene);Acc:827000] |
| ARA1 | ARA1 | AT4G16130 | Arabidopsis thaliana | arabinose kinase [Source:NCBI gene (formerly Entrezgene);Acc:827299] |
| LPD2 | AT4G16155 | AT4G16155 | Arabidopsis thaliana | dihydrolipoamide dehydrogenase [Source:NCBI gene (formerly Entrezgene);Acc:827307] |
| RABA1D | RABA1d | AT4G18800 | Arabidopsis thaliana | RAB GTPase homolog A1D [Source:NCBI gene (formerly Entrezgene);Acc:827614] |
| TPP2 | TPP2 | AT4G20850 | Arabidopsis thaliana | tripeptidyl peptidase ii [Source:NCBI gene (formerly Entrezgene);Acc:827833] |
| RPN1B | RPN1B | AT4G28470 | Arabidopsis thaliana | 26S proteasome regulatory subunit S2 1B [Source:NCBI gene (formerly Entrezgene);Acc:828964] |
| CAD5 | CAD5 | AT4G34230 | Arabidopsis thaliana | cinnamyl alcohol dehydrogenase 5 [Source:NCBI gene (formerly Entrezgene);Acc:829572] |
| VAB2 | VAB2 | AT4G38510 | Arabidopsis thaliana | ATPase, V1 complex, subunit B protein [Source:NCBI gene (formerly Entrezgene);Acc:830008] |
| CDC48E | AtCDC48C | AT5G03340 | Arabidopsis thaliana | ATPase, AAA-type, CDC48 protein [Source:NCBI gene (formerly Entrezgene);Acc:831870] |
| UAP56B | UAP56b | AT5G11200 | Arabidopsis thaliana | DEAD/DEAH box RNA helicase family protein [Source:NCBI gene (formerly Entrezgene);Acc:830990] |
| ASP3 | ASP3 | AT5G11520 | Arabidopsis thaliana | aspartate aminotransferase 3 [Source:NCBI gene (formerly Entrezgene);Acc:831024] |
| TUBB6 | TUB6 | AT5G12250 | Arabidopsis thaliana | beta-6 tubulin [Source:NCBI gene (formerly Entrezgene);Acc:831100] |
| ARFA1B | ARFA1B | AT5G14670 | Arabidopsis thaliana | ADP-ribosylation factor A1B [Source:NCBI gene (formerly Entrezgene);Acc:831319] |
| RPS9B | AT5G15200 | AT5G15200 | Arabidopsis thaliana | Ribosomal protein S4 [Source:NCBI gene (formerly Entrezgene);Acc:831372] |
| UGD3 | UGD3 | AT5G15490 | Arabidopsis thaliana | UDP-glucose 6-dehydrogenase family protein [Source:NCBI gene (formerly Entrezgene);Acc:831402] |
| CCT2 | AT5G20890 | AT5G20890 | Arabidopsis thaliana | TCP-1/cpn60 chaperonin family protein [Source:NCBI gene (formerly Entrezgene);Acc:832213] |
| CCT3 | AT5G26360 | AT5G26360 | Arabidopsis thaliana | TCP-1/cpn60 chaperonin family protein [Source:NCBI gene (formerly Entrezgene);Acc:832705] |
| MIRO1 | MIRO1 | AT5G27540 | Arabidopsis thaliana | MIRO-related GTP-ase 1 [Source:NCBI gene (formerly Entrezgene);Acc:832814] |
| RPL18C | AT5G27850 | AT5G27850 | Arabidopsis thaliana | Ribosomal protein L18e/L15 superfamily protein [Source:NCBI gene (formerly Entrezgene);Acc:832848] |
| RABA1C | RABA1c | AT5G45750 | Arabidopsis thaliana | RAB GTPase homolog A1C [Source:NCBI gene (formerly Entrezgene);Acc:834614] |
| ERF1-1 | ERF1-1 | AT5G47880 | Arabidopsis thaliana | eukaryotic release factor 1-1 [Source:NCBI gene (formerly Entrezgene);Acc:834839] |
| CLPC1 | CLPC1 | AT5G50920 | Arabidopsis thaliana | CLPC homologue 1 [Source:NCBI gene (formerly Entrezgene);Acc:835165] |
| ASN2 | ASN2 | AT5G65010 | Arabidopsis thaliana | asparagine synthetase 2 [Source:NCBI gene (formerly Entrezgene);Acc:836625] |
| Leucine--tRNA ligase, cytoplasmic | F4I116 | AT1G09620 | Arabidopsis thaliana | Leucine--tRNA ligase, cytoplasmic |
| At1g23130/T26J12_10 | O49304 | AT1G23130 | Arabidopsis thaliana | At1g23130/T26J12_10 |
| S-adenosyl-L-methionine-dependent methyltransferases superfamily protein | F4I0B5 | AT1G55450 | Arabidopsis thaliana | S-adenosyl-L-methionine-dependent methyltransferases superfamily protein |
| Probable nucleoredoxin 1 | O80763 | AT1G60420 | Arabidopsis thaliana | Probable nucleoredoxin 1 |
| Probable fructokinase-1 | Q9SID0 | AT2G31390 | Arabidopsis thaliana | Probable fructokinase-1 |
| Translation initiation factor eIF2B subunit epsilon | O64760 | AT2G34970 | Arabidopsis thaliana | Translation initiation factor eIF2B subunit epsilon |
| Dihydrolipoyllysine-residue acetyltransferase component 2 of pyruvate dehydrogenase complex, mitochondrial | Q8RWN9 | AT3G13930 | Arabidopsis thaliana | Dihydrolipoyllysine-residue acetyltransferase component 2 of pyruvate dehydrogenase complex, mitochondrial |
| Coatomer subunit beta' | A0A1I9LTK3 | AT3G15980 | Arabidopsis thaliana | Coatomer subunit beta' |
| 40S ribosomal protein S4 | F4K5C7 | AT5G07090 | Arabidopsis thaliana | 40S ribosomal protein S4 |
| Dihydrolipoyllysine-residue succinyltransferase component of 2-oxoglutarate dehydrogenase complex 1, mitochondrial | Q9FLQ4 | AT5G55070 | Arabidopsis thaliana | Dihydrolipoyllysine-residue succinyltransferase component of 2-oxoglutarate dehydrogenase complex 1, mitochondrial |
| Histone H4 | P59259 | AT5G59970 | Arabidopsis thaliana | Histone H4 |
| Indole-3-acetic acid-amido synthetase GH3.17 | Q9FZ87 | At1g28130 | Arabidopsis thaliana | Indole-3-acetic acid-amido synthetase GH3.17 |
| Mov34/MPN/PAD-1 family protein | A8MR12 | At5g23540 | Arabidopsis thaliana | Mov34/MPN/PAD-1 family protein |

